# A SARS-CoV-2-Human Protein-Protein Interaction Map Reveals Drug Targets and Potential Drug-Repurposing

**DOI:** 10.1101/2020.03.22.002386

**Authors:** David E. Gordon, Gwendolyn M. Jang, Mehdi Bouhaddou, Jiewei Xu, Kirsten Obernier, Matthew J. O’Meara, Jeffrey Z. Guo, Danielle L. Swaney, Tia A. Tummino, Ruth Huettenhain, Robyn M. Kaake, Alicia L. Richards, Beril Tutuncuoglu, Helene Foussard, Jyoti Batra, Kelsey Haas, Maya Modak, Minkyu Kim, Paige Haas, Benjamin J. Polacco, Hannes Braberg, Jacqueline M. Fabius, Manon Eckhardt, Margaret Soucheray, Melanie J. Bennett, Merve Cakir, Michael J. McGregor, Qiongyu Li, Zun Zar Chi Naing, Yuan Zhou, Shiming Peng, Ilsa T. Kirby, James E. Melnyk, John S. Chorba, Kevin Lou, Shizhong A. Dai, Wenqi Shen, Ying Shi, Ziyang Zhang, Inigo Barrio-Hernandez, Danish Memon, Claudia Hernandez-Armenta, Christopher J.P. Mathy, Tina Perica, Kala B. Pilla, Sai J. Ganesan, Daniel J. Saltzberg, Rakesh Ramachandran, Xi Liu, Sara B. Rosenthal, Lorenzo Calviello, Srivats Venkataramanan, Jose Liboy-Lugo, Yizhu Lin, Stephanie A. Wankowicz, Markus Bohn, Phillip P. Sharp, Raphael Trenker, Janet M. Young, Devin A. Cavero, Joseph Hiatt, Theodore L. Roth, Ujjwal Rathore, Advait Subramanian, Julia Noack, Mathieu Hubert, Ferdinand Roesch, Thomas Vallet, Björn Meyer, Kris M. White, Lisa Miorin, Oren S. Rosenberg, Kliment A Verba, David Agard, Melanie Ott, Michael Emerman, Davide Ruggero, Adolfo García-Sastre, Natalia Jura, Mark von Zastrow, Jack Taunton, Alan Ashworth, Olivier Schwartz, Marco Vignuzzi, Christophe d’Enfert, Shaeri Mukherjee, Matt Jacobson, Harmit S. Malik, Danica G. Fujimori, Trey Ideker, Charles S. Craik, Stephen Floor, James S. Fraser, John Gross, Andrej Sali, Tanja Kortemme, Pedro Beltrao, Kevan Shokat, Brian K. Shoichet, Nevan J. Krogan

## Abstract

An outbreak of the novel coronavirus SARS-CoV-2, the causative agent of COVID-19 respiratory disease, has infected over 290,000 people since the end of 2019, killed over 12,000, and caused worldwide social and economic disruption^1,2^. There are currently no antiviral drugs with proven efficacy nor are there vaccines for its prevention. Unfortunately, the scientific community has little knowledge of the molecular details of SARS-CoV-2 infection. To illuminate this, we cloned, tagged and expressed 26 of the 29 viral proteins in human cells and identified the human proteins physically associated with each using affinity-purification mass spectrometry (AP-MS), which identified 332 high confidence SARS-CoV-2-human protein-protein interactions (PPIs). Among these, we identify 67 druggable human proteins or host factors targeted by 69 existing FDA-approved drugs, drugs in clinical trials and/or preclinical compounds, that we are currently evaluating for efficacy in live SARS-CoV-2 infection assays. The identification of host dependency factors mediating virus infection may provide key insights into effective molecular targets for developing broadly acting antiviral therapeutics against SARS-CoV-2 and other deadly coronavirus strains.

## MAIN

The current pandemic of COVID-19 (<u>Co</u>rona<u>vi</u>rus <u>D</u>isease-2019), a respiratory disease that has led to over 290,000 confirmed cases and 12,000 fatalities in over 100 countries since its emergence in late 2019^3,4^, is caused by a novel virus strain, SARS-CoV-2, an enveloped, positive-sense, single-stranded RNA betacoronavirus of the family *Coronaviridae*. Coronaviruses infecting humans historically included several mild common cold viruses e.g. hCoV-OC43, HKU, 229E^5^. However, over the past two decades, highly pathogenic human coronaviruses have emerged, including SARS-CoV in 2002 and 2003 with 8,000 cases worldwide and a death rate of ∼10%, and MERS-CoV in 2012, which caused 2,500 confirmed cases and a fatality rate of 36%^6^. Infection with these highly pathogenic coronaviruses can result in Acute Respiratory Distress Syndrome (ARDS), which may lead to long-term reduction in lung function, arrhythmia, and death. Compared to MERS or SARS^7,8^, SARS-CoV-2 appears to spread more efficiently, making it difficult to contain and increasing its pandemic potential. To devise therapeutic strategies to counteract SARS-CoV-2 infection, it is crucial to develop a comprehensive understanding of how this coronavirus hijacks the host during the course of infection, and to apply this knowledge towards developing both new drugs and repurposing existing ones.

So far, no clinically available antiviral drugs have been developed for SARS-CoV, SARS-CoV-2 or MERS-CoV. Clinical trials are ongoing for treatment of COVID-19 with the nucleotide analog RNA-dependent RNA Polymerase (RdRP) inhibitor remdesivir^9–11^, and recent data suggests a new nucleotide analog may be effective against SARS-CoV-2 infection in laboratory animals^12^. Clinical trials on several vaccine candidates are also underway^13^, as well as trials of repurposed host-directed compounds inhibiting the human protease TMPRSS2^14^. We believe there is great potential in systematically exploring the host dependencies of the SARS-CoV-2 virus to identify other host proteins already targeted with existing drugs. Therapies targeting the host-virus interface, where mutational resistance is arguably less likely, could potentially present durable, broad-spectrum treatment modalities^15^. Unfortunately, our minimal knowledge of the molecular details of SARS-CoV-2 infection precludes a comprehensive evaluation of small molecule candidates for host-directed therapies. We sought to address this knowledge gap by systematically mapping the interaction landscape between SARS-CoV-2 proteins and human proteins.

### Cloning and expression of predicted SARS-CoV-2 proteins

Sequence analysis of SARS-CoV-2 isolates suggests that the 30kb genome encodes as many as 14 open reading frames (Orfs). The 5’ Orf1a / Orf1ab encodes polyproteins, which are auto-proteolytically processed into 16 non-structural proteins (Nsp1-16) which form the replicase / transcriptase complex (RTC). The RTC consists of multiple enzymes, including the papain-like protease (Nsp3), the main protease (Nsp5), the Nsp7-Nsp8 primase complex, the primary RNA-dependent RNA polymerase (Nsp12), a helicase/triphosphatase (Nsp13), an exoribonuclease (Nsp14), an endonuclease (Nsp15), and N7- and 2’O-methyltransferases (Nsp10/Nsp16)^1,16,17^. At the 3’ end of the viral genome, as many as 13 Orfs are expressed from nine predicted sub-genomic RNAs. These include four structural proteins: Spike (S), Envelope (E), Membrane (M) and Nucleocapsid (N)^17^, and nine putative accessory factors (Fig. 1a)^1,16^. In genetic composition, the SARS-CoV-2 genome is very similar to SARS-CoV: each has an Orf1ab encoding 16 predicted Nsps and each has the four typical coronavirus structural proteins. However, they differ in their complement of 3’ open reading frames: SARS-CoV-2 possesses an Orf3b and Orf10 with limited detectable protein homology to SARS-CoV^16^, and its Orf8 is intact while SARS-CoV encodes Orf8a and Orf8b (Fig. 1b)^1,16,18^.

**Figure 1:**
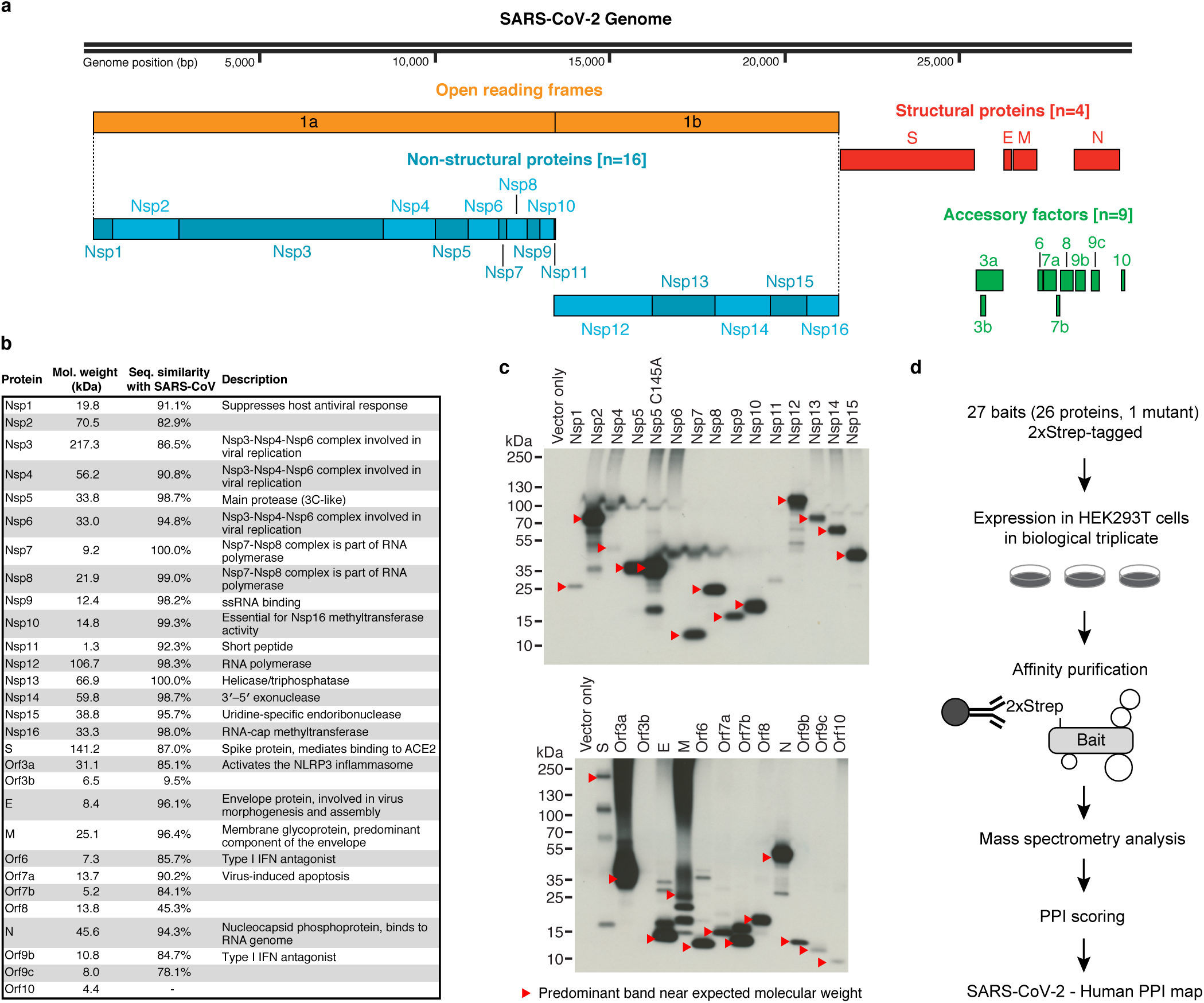
AP-MS Workflow for Identification of SARS-CoV-2 Host Protein-Protein Interactions. (**a**) SARS-CoV-2 genome annotation. (**b**) Table of the SARS-CoV-2 proteins, including molecular weight, sequence similarity with the SARS-CoV homolog, and inferred function based on the SARS-CoV homolog. (**c**) Immunoblot detection of 2xStrep tag demonstrates expression of each bait in input samples, as indicated by red arrowhead. (**d**) Experimental workflow for expressing each 2xStrep tagged SARS-CoV-2 fusion protein in biological triplicate in HEK293T cells, followed by affinity purification-mass spectrometry, and PPI scoring to identify 332 high confidence protein-protein interactions.

Mature Nsps and all predicted proteins expressed from other SARS-CoV-2 Orfs (27 proteins plus one mutant) were codon optimized and cloned into a mammalian expression vector with a 2xStrep tag fused to either the N- or C-terminus^19^. Protein expression plasmids were transfected into human HEK293T cells, which were incubated for 40 hours before lysis in a mild detergent buffer. Viral proteins were affinity purified using MagStrep beads. Beads were washed, on-bead digestion was performed overnight, and peptides were desalted and analyzed by protein mass spectrometry. High confidence interactors were identified using SAINTexpress and the MiST algorithm^19,20^.

To verify viral protein expression, we performed an anti-Strep western blot on input cell lysate, and with the exception of Nsp4, Nsp6, Nsp11, and Orf3b, we observed bands consistent with predicted protein sizes (24 of 28 constructs). Despite the lack of detection via western blot we were able to detect expression of viral peptides Nsp4, Nsp6, and Orf3b in the proteomic analysis. The fourth construct not confirmed by western blot, the small peptide Nsp11, had a predicted molecular mass of 4.8 kDa (including tag) but an apparent mass of approximately 30 kDa (Fig. 1c). SARS-CoV-2 Orf3b has limited homology to SARS-CoV Orf3b^16^, and its restricted expression as measured by both western blot and AP-MS may indicate that it is either not a bonafide protein coding gene or that it requires co-expression of other SARS-CoV-2 proteins for robust expression. Ultimately we proceeded with analysis of protein interaction data from 27 baits, and excluded Orf7b, which had high prey detection background (Fig. 1d).

### Global analysis of SARS-CoV-2 host interacting proteins

Our affinity purification-mass spectrometry analysis identified 332 protein interactions between SARS-CoV-2 proteins and human proteins (Extended Data Fig. 1, Supplementary Tables 1, 2; also see Fig. 3). We studied the interacting human proteins in regards to their cell biology, anatomical expression patterns, expression changes during SARS-CoV-2 infection^21^ and in relation to other maps of pathogen interacting proteins^19,22–30^(Fig. 2a). For each of the viral proteins, we performed Gene Ontology enrichment analysis (Fig. 2b, Extended Data Fig. 2), identifying the major cell biological processes of the interacting proteins, including lipoprotein metabolism (S), nuclear transport (Nsp7), and biogenesis of ribonucleoprotein (Nsp8). To discover potential binding interfaces, we performed an enrichment for domain families within the interacting proteins of each viral bait (Extended Data Fig. 3). As examples, we note the enrichment of DNA polymerase domains in the interactors of Nsp1 and the enrichment of bromodomains and extra-terminal domain (BET) family for interactors of E (see also Figs. 3, 4). In line with the latter, the interactors of E are also enriched in genes annotated for binding to acetylated histones (Fig. 2b).

**Figure 2:**
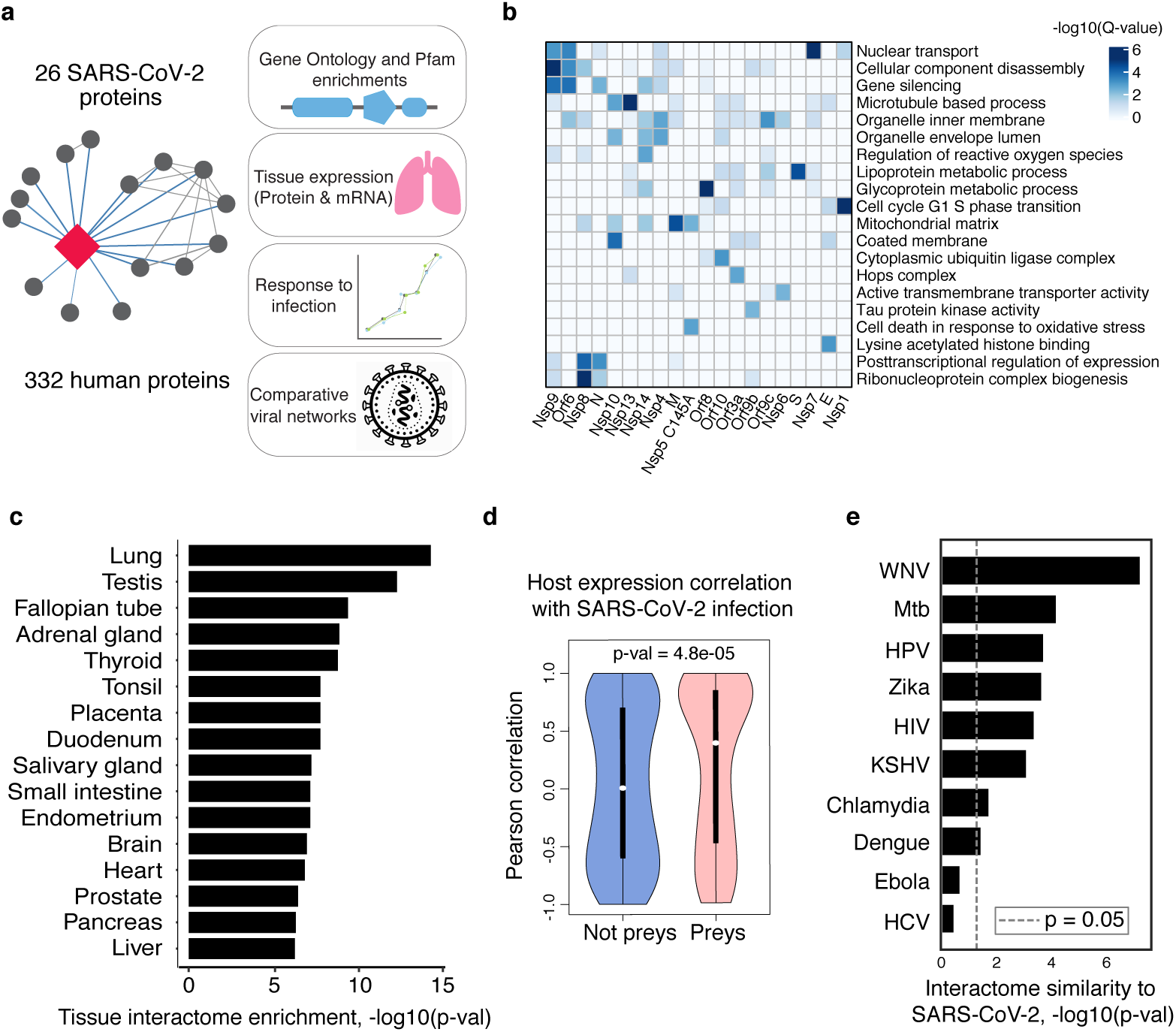
Global Analysis of SARS-CoV-2 Protein Interactions. (**a**) Overview of global analyses performed. (**b**) Gene Ontology (GO) enrichment analysis performed on the human interacting proteins of each viral protein (Methods). The top GO term of each viral protein was selected for visualization. (**c**) Degree of differential protein expression for the human interacting proteins across human tissues. We obtained protein abundance values for the proteome in 29 human tissues and calculated the median level of abundance for the set of human interacting proteins (top 16 tissues shown). This median value was then compared with the distribution of abundance values for the full proteome in each tissue and summarized as a Z-score from which a p-value was calculated and adjusted for multiple tests. (**d**) Distribution of correlation of protein level changes during SARS-CoV-2 infection for pairs of viral-human proteins. **(e)** Significance of the overlap of human interacting proteins between SARS-CoV-2 and other pathogens.

**Figure 3:**
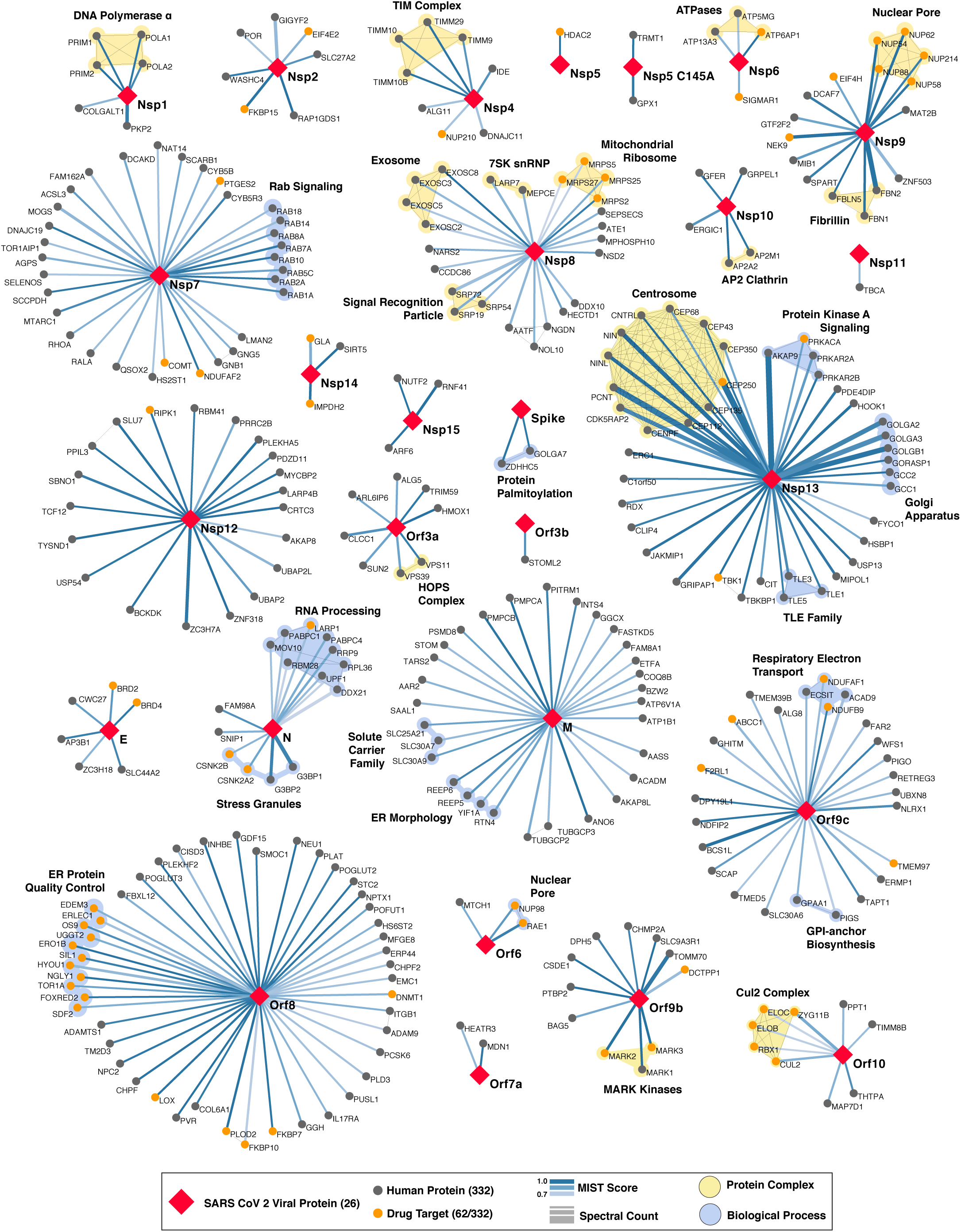
SARS-CoV-2 Protein-Protein Interaction Network. In total, 332 high confidence interactions are represented between 26 SARS-CoV-2 proteins and their human interactors. Red diamonds represent a SARS-CoV-2 viral protein, interacting human host proteins are represented with circles, with drug targets in orange. Edge color is proportional to MiST score and edge thickness proportional to spectral counts. Physical interactions among host proteins are noted as thin black lines, protein complexes are highlighted in yellow, and proteins sharing the same biological process are highlighted in blue.

**Figure 4:**
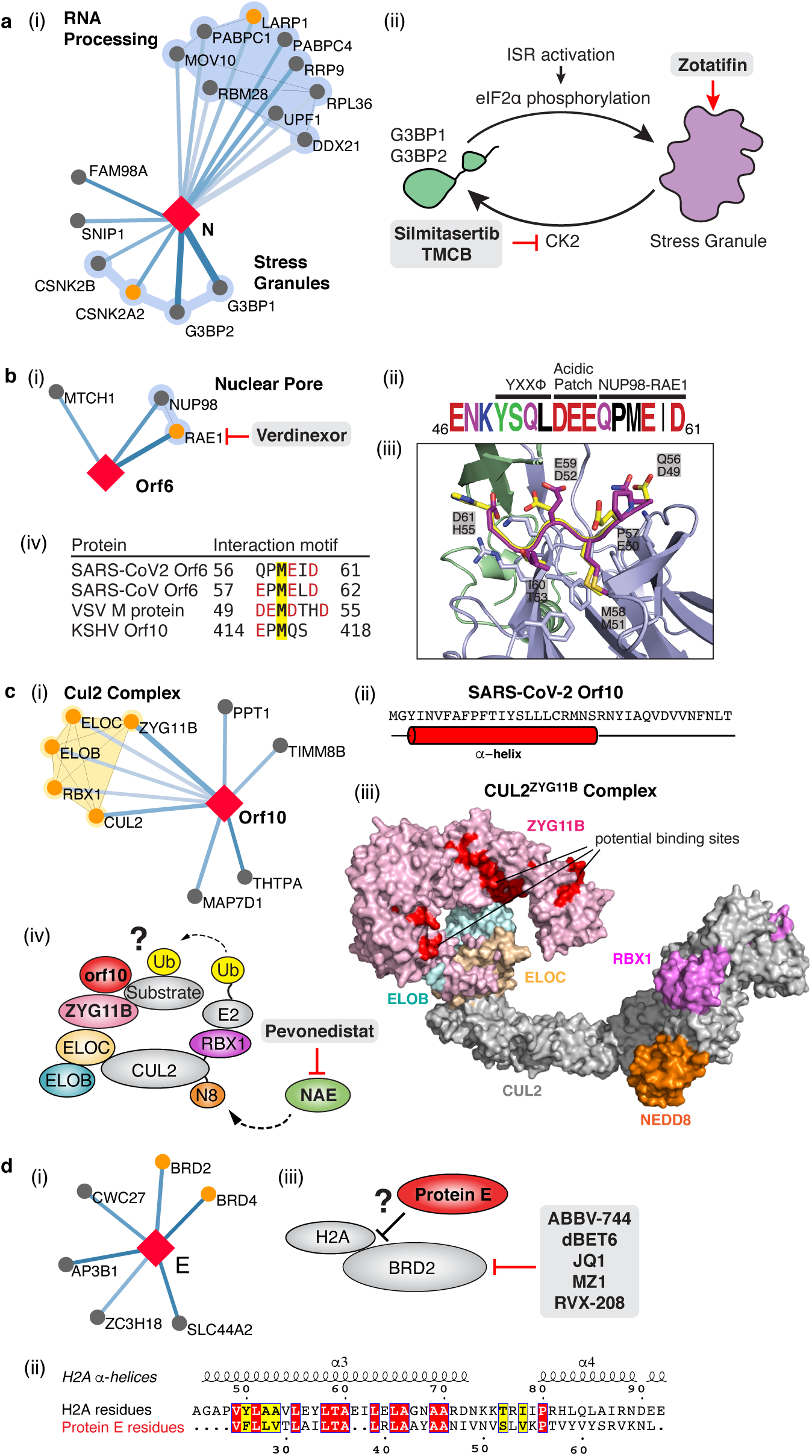
The SARS-CoV-2 interactome reveals novel aspects of SARS-CoV-2 biology that can be targeted pharmacologically. (**a**) Protein N targets stress granule proteins. (i) Protein N interactome. (ii) Model for therapeutic targeting of N interactions in the formation of stress granules (SGs). SGs are known to exhibit antiviral activity, with the integrative stress response (ISR) inducing eIF2α phosphorylation and SG formation, and Casein kinase II (CK2) disrupting and preventing the formation of SGs. By activating SG formation, or inhibiting CK2, the cellular environment could potentially shift to a more antiviral state. (**b**) Orf6 interacts with an interferon-inducible mRNA nuclear export complex. (i) Orf6 interactome including small molecule inhibitors for RAE. (ii) Annotated C-terminal sequence of SARS-CoV-2 Orf6, highlighting previously described trafficking motifs and the putative NUP98-RAE1 binding sequence. Colors indicate chemical properties of amino acids: polar (G,S,T,Y,C, green), neutral (Q,N, purple), basic (K, R, H, blue), acidic (D, E, red), and hydrophobic (A, V, L, I, P, W, F, M, black). (iii) SARS-CoV-2 Orf6 carboxy-terminal peptide modeled into the binding site of the VSV M protein-NUP98-RAE1 complex (PDB ID: 4OWR). Orf6 shown in dark purple, M protein in yellow, NUP98 in green, and RAE1 in light purple. Orf6 and M protein residues labeled. RAE1 hydrophobic residues contacting the key methionine and basic patch residues of RAE1 and NUP98 are shown. (iv) Putative NUP98-RAE1 interaction motifs present in proteins from several viral species. The consensus motif consists of negatively charged residues (red) surrounding a conserved methionine (yellow). (**c**) Orf10 interacts with the CUL2^ZYG11B^ complex. (i) Orf10 interactome. (ii) The secondary structure of Orf10 contains an alpha helix motif. (iii) Surface representation of the homology model for CUL2^ZYG11B^ complex, residues that are conserved amongst ZYG11B orthologues from various species are indicated in red are likely protein interaction surfaces for binding substrates and other proteins. (iv) A possible model of how Orf10 binds to the CUL2^ZYG11B^ complex to hijack the complex for ubiquitination or viral restriction factors and how it can be targeted pharmacologically. (**d**) Envelope (E) interacts with bromodomain proteins. (i) E interactome. (ii) Sequence alignment of highlighted regions of E and Histone 2A (H2A). The positions with identical and similar amino acid residues are highlighted in red and yellow, respectively. Note the greater hydrophobicity of E may indicate a part of the alignment represents a transmembrane segment. (iii) Model of how E might mimic the BRD2 native interaction partner Histone 2A and how BRD2 can be targeted pharmacologically.

While the cells used for these AP-MS experiments, HEK-293T kidney cells, are permissive to SARS-CoV-2 infection^31^, they do not represent the primary physiological site of infection. Therefore, we asked whether the host proteins bound by SARS-CoV-2 might be specifically relevant to the virus’s typical environment, lung tissue. We tested if the interacting human proteins were preferentially highly expressed, at the protein level, in any of 29 human tissues^32^, which identified the lung as the tissue with the highest expression of the prey proteins relative to the average proteome (Fig. 2c). In accordance to this, when compared to overall RefSeq gene expression in the lung (median=3.198 TPM), the interacting proteins were more highly expressed (median=25.52 TPM, p=0.0007; t-test, Extended Data Fig. 4) and were also specifically enriched in lung expression relative to other tissues (Extended Data Fig. 5). These data support the hypothesis that SARS-CoV-2 preferentially hijacks proteins available in lung tissue and the degree of tissue specific expression may facilitate the drug targeting of host factors with fewer systemic side-effects; this observation is particularly compelling given its derivation from HEK293 cells, which are themselves not lung derived. To further study the relevance of our map, we analyzed the recently reported protein abundance changes during SARS-CoV-2 infection^33^. We calculated, when possible, the correlation between changes in abundance of viral proteins and their human interaction partners across the 4 timepoints reported. Interacting pairs tended to have stronger correlated changes than other pairs of viral-human proteins (Fig. 2d, KS test p-value=4.8e−05), arguing that the AP-MS derived interactions are of high relevance for the target tissue and the infection context. We compared our SARS-CoV-2 interaction map with those derived for 10 other pathogens (Fig. 2e), identifying West Nile Virus (WNV)^23^ and Mycobacterium tuberculosis (Mtb)^27^ as having the most similar host protein interaction partners. The association with Mtb is particularly interesting considering it also infects lung tissue.

Finally, we studied the evolutionary properties of the host proteins. Successful virus spread in a diverse population in theory could be facilitated by a reliance on conserved host molecular components, therefore we analyzed the conservation of the human proteins identified in the SARS-CoV-2 interactome. Relative to a control sample of genes, the 332 SARS-CoV-2 interacting human proteins had depleted missense and premature stop mutations in gnomAD^34^, indicating that they have reduced genetic variation in human populations (Extended Data Fig. 6).

### The SARS-CoV-2 interactome reveals novel aspects of SARS-CoV-2 biology

Our study highlighted interactions between SARS-CoV-2 proteins and human proteins with a range of functions including DNA replication (Nsp1), epigenetic and gene expression regulators (Nsp5, Nsp8, Nsp13, E), vesicle trafficking (Nsp6, Nsp7, Nsp10, Nsp13, Nsp15, Orf3a, E, M, Orf8), lipid modification (Spike), RNA processing and regulation (Nsp8, N), ubiquitin ligases (Orf10), signaling (Nsp8, Nsp13, N, Orf9b), nuclear transport machinery (Nsp9, Nsp15, Orf6), cytoskeleton (Nsp1, Nsp13), mitochondria (Nsp4, Nsp8, Orf9c), and extracellular matrix (Nsp9) (Fig. 3).

A prominent number of interactions were related to lipid modifications and vesicle trafficking. Interestingly, the Spike protein (S) interacts with the GOLGA7-ZDHHC5 acyl-transferase complex, which likely mediates palmitoylation on its cytosolic tail (see also Appendix)^35^. Palmitoylation has been reported to facilitate membrane fusion by SARS-CoV Spike and suggests a potential target for therapeutic inhibition^36^. Interestingly, ZDHHC5 also has a published role in allowing anthrax toxin to enter cells, suggesting that inhibition of this enzyme could have broad utility^37^. Host interactions of Nsp8 (signal recognition particle), Orf8 (endoplasmic reticulum quality control), M (ER structural morphology proteins), Nsp13 (golgins) may facilitate the dramatic reconfiguration of ER/Golgi trafficking during coronavirus infection, and interactions in peripheral compartments by Nsp6 and M (vacuolar ATPase), Nsp7 (Rabs), Nsp10 (AP2), E (AP3), and Orf3a (HOPS) may also modify endomembrane compartments to favor coronavirus replication.

We identified protein-protein interactions with the main protease Nsp5, using both wild-type and catalytic dead (C145A) constructs. For wild-type Nsp5, we identified one high-confidence interaction, the epigenetic regulator histone deacetylase 2 (HDAC2), and predicted a cleavage site between the HDAC domain and the nuclear localization sequence, suggesting that Nsp5 may inhibit HDAC2 transport into the nucleus (Extended Data Fig. 7), potentially impacting the published functions of HDAC2 in mediating inflammation and interferon response^38,39^. We also identified an interaction of Nsp5 (C145A) with tRNA methyltransferase 1 (TRMT1), which is responsible for synthesis of the dimethylguanosine (m2,2G) base modification on both nuclear and mitochondrial tRNAs^40^. We predict TRMT1 is also cleaved by Nsp5, removing its zinc finger and nuclear localization signal and likely resulting in an exclusively mitochondrial localization (Extended Data Fig. 7).

### SARS-CoV-2 interacts with multiple innate immune pathways

We identified a number of cellular proteins implicated in innate immune signaling that are targeted by several SARS-CoV-2 viral proteins. Interestingly, we found that Nsp13 interacts with two key players of IFN signaling pathway including TANK-binding kinase 1 (TBK1) and TANK-binding kinase 1-binding protein 1 (TBKBP1/SINTBAD). SINTBAD acts as a critical adaptor protein between TBK1 and IKKi and therefore mediates induction of IRF-dependent transcription^41^. Further, Nsp13 interacts with multiple proteins of the TLE family, which are known to modulate NF-κB inflammatory response^42–44^. RNF41/Nrdp1, an E3 ubiquitin ligase is targeted by Nsp15 protein which promotes activation of TBK1 and IRF3 and therefore increases type I interferon production^45^. Two other E3 ubiquitin ligases, TRIM59 and MIB1 regulate antiviral innate immune signaling and are usurped by Orf3a and Nsp9, respectively^46,47^. Orf9c protein was found to interact with multiple proteins that modulate IkB kinase and NF-kB signaling pathway including NLRX1, F2RL1, NDFIP2^48–50^. We also found that Orf9b interacts with a mitochondrial import receptor, Tom70, which acts as an essential adaptor linking MAVS to TBK1/IRF3, resulting in the activation of IRF-3^51^.

N targets stress granule protein G3BP1, an essential antiviral protein which is known to induce innate immune response through multiple mechanisms^52–54^. Common among *coronaviridae* is the manipulation of stress granules (SG) and related RNA biology, possibly leading to suppression of stress granules and host translation shutoff^55^. This functionality seems to benefit viral replication, as stress granules are inhibitory to replication of MERS-CoV^56^ and other viruses^57^. The SARS-CoV-2 nucleocapsid (N) interactome includes many host mRNA binding proteins, including the SG related factors G3BP1/2, the mTOR translational repressors LARP1, and the protein kinases CK2 (Fig. 4a). SGs are induced by protein kinase R (PKR)-mediated phosphorylation of eIF2α upon viral dsRNA recognition^57^. Promoting G3BP aggregation via the eIF4A inhibitor Zotatafin^58,59^ or reducing SG disassembly by Silmitasertib inhibition of CK2^60^ warrant investigation for treatment of SARS-CoV-2. The mTOR inhibitor rapamycin disrupts the binding of LARP1 to mTORC1^61^ and has been shown to reduce MERS infection by ∼60% *in vitro*^*62*^, another drug that could be tested for repurposing.

Orf6 of SARS-CoV has been shown to play a role in antagonizing host interferon signaling^63^; we identified a novel, high-confidence interaction between SARS-CoV-2 Orf6 and NUP98-RAE1, an interferon-inducible mRNA nuclear export complex^64^ that is hijacked or degraded by multiple viruses, including VSV, Influenza-A, KSHV, and Polio, and is a restriction factor for Influenza-A infection^58,60,62,65^. The X-ray structure of VSV M protein complexed with NUP98-RAE1^66^ reveals key binding interactions that include a buried methionine residue on the M-protein packing into a hydrophobic pocket in RAE1, as well as neighboring acidic residues interacting with a basic patch on the NUP98-RAE1 complex (Fig 4b). These binding features are also present in a conserved motif in the C-terminal region of SARS-CoV-2 Orf6 (Fig. 4b, Extended Data Fig. 8), providing a structural hypothesis for the observed SARS-CoV-2–NUP98-RAE1 interaction. Moreover, a peptide containing the binding region of the VSV M protein was previously shown to outcompete RNA binding to NUP98-RAE1, suggesting a role in interfering with mRNA export^66^. These observations suggest a viral strategy to target the RNA nuclear export activity of RAE1, potentially revealing a mode of interferon antagonism by SARS-CoV-2.

### The novel Orf10 of SARS-CoV-2 interacts with a Cullin ubiquitin ligase complex

Viruses commonly hijack ubiquitination pathways for replication and pathogenesis^67^. The novel Orf10 of SARS-CoV-2 interacts with multiple members of a Cullin 2 (CUL2) RING E3 ligase complex (Fig. 4c), specifically the CUL2^ZYG11B^ complex. ZYG11B, a substrate adapter of CUL2 that targets substrates with exposed N-terminal glycines for degradation^65^, is the highest scoring protein in the Orf10 interactome suggesting its direct interaction with Orf10. Orf10 may bind to the CUL2^ZYG11B^ complex and hijack it for ubiquitination and degradation of restriction factors. The ubiquitin transfer to a substrate requires neddylation of CUL2 via NEDD8-activating enzyme (NAE), a druggable target that can be inhibited by the small molecule Pevonedistat^68^ (Fig. 4c).

### SARS-CoV-2 envelope interacts with bromodomain proteins

Surprisingly, we find that the transmembrane protein E binds to the bromodomain-containing proteins BRD2 and BRD4 (Fig. 4d, Extended Data Fig. 9), potentially disrupting BRD-histone binding by mimicking histone structure. BRD2 is a member of the bromodomain and extra-terminal (BET) domain family whose members bind acetylated histones to regulate gene transcription^69^. The N-terminus of histone 2A shares local sequence similarity over an alpha-helix of approximately 15 residues, some of which are in a transmembrane segment, of Protein E (Fig. 4d). Moreover, this matching region of the histone is spanned by acetylated lysine residues shown to bind BRD2^70^. This analysis may suggest that Protein E mimics the histone to disrupt its interaction with BRD2, thus inducing changes in host’s protein expression that are beneficial to the virus.

For a more comprehensive overview of the virus-host interactions we detected, see Supplemental Discussion.

### Identification of existing drugs targeting SARS-CoV-2 human host factors

To identify small molecules targeting human proteins in the SARS-CoV-2 interactome, we sought ligands known to interact with the human proteins, often directly but also by pathway and complexes, drawing on chemoinformatics databases and analyses (Methods). Molecules were prioritized by the statistical significance of the interaction between the human and viral proteins; by their status as approved drugs, investigational new drugs (INDs, “clinical” in Table 1a,b), or as preclinical candidates; by their apparent selectivity; and by their availability (for purchase availability notes, see Supplemental Tables 3 and 4). Chemoinformatics searches yielded 15 approved drugs, four investigational new drugs (clinical), and 18 pre-clinical candidates (Table 1a), while specialist knowledge revealed 12 approved drugs, 10 investigational new drugs (clinical), and 10 preclinical candidates (Table 1b). Of the 332 human targets that interact with the viral bait proteins with high significance, 63 have drugs/INDs/preclinical molecules that modulate them (Fig. 3). If we reduce our protein interaction score threshold slightly, we find an additional four human targets, revealing a total of 67 human targets (Supplementary Tables 3 and 4). The drug-human protein associations may be overlaid on top of our protein interaction network, highlighting potentially druggable host interactions (Fig. 5a).

**Table 1a.**
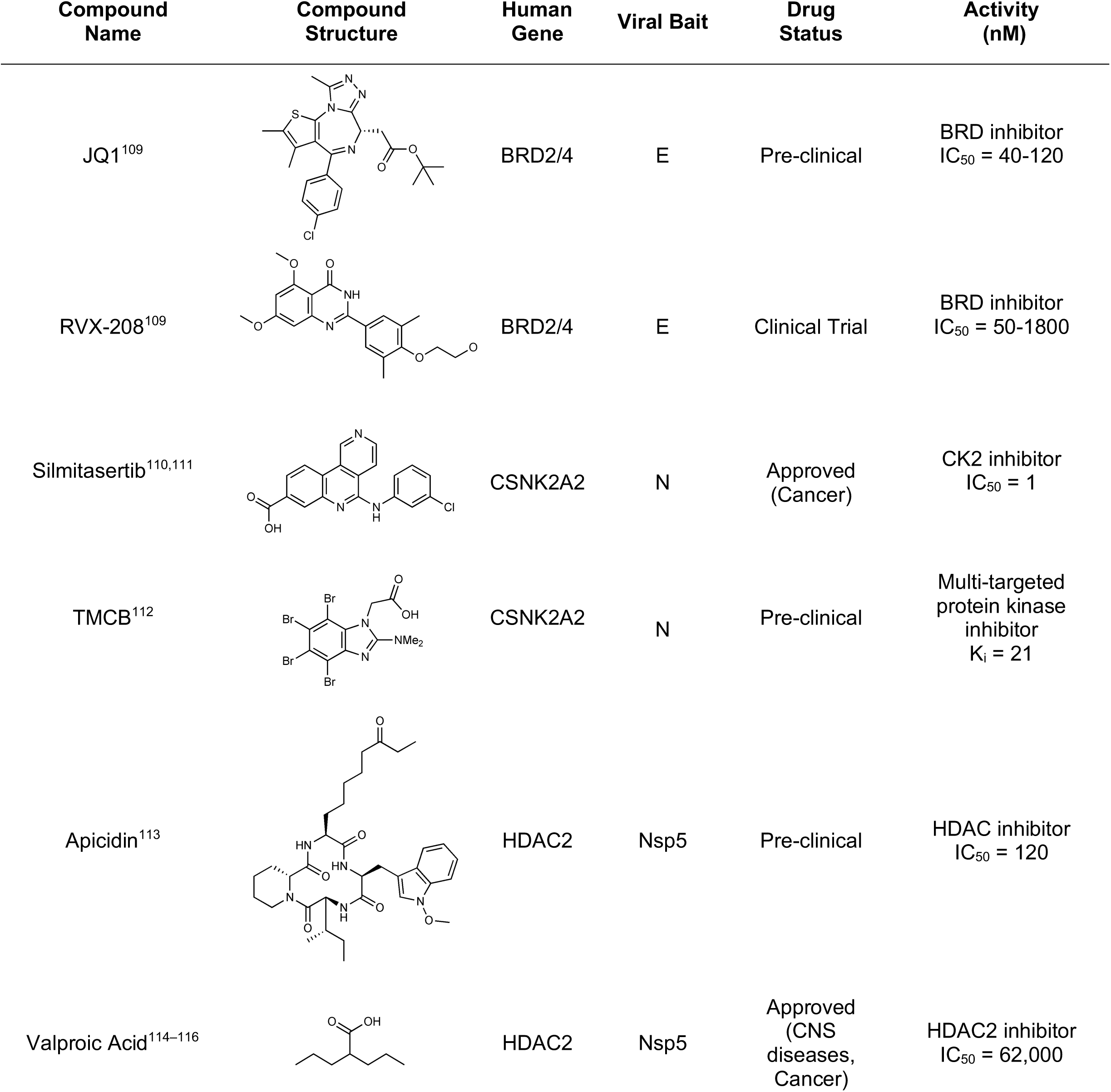
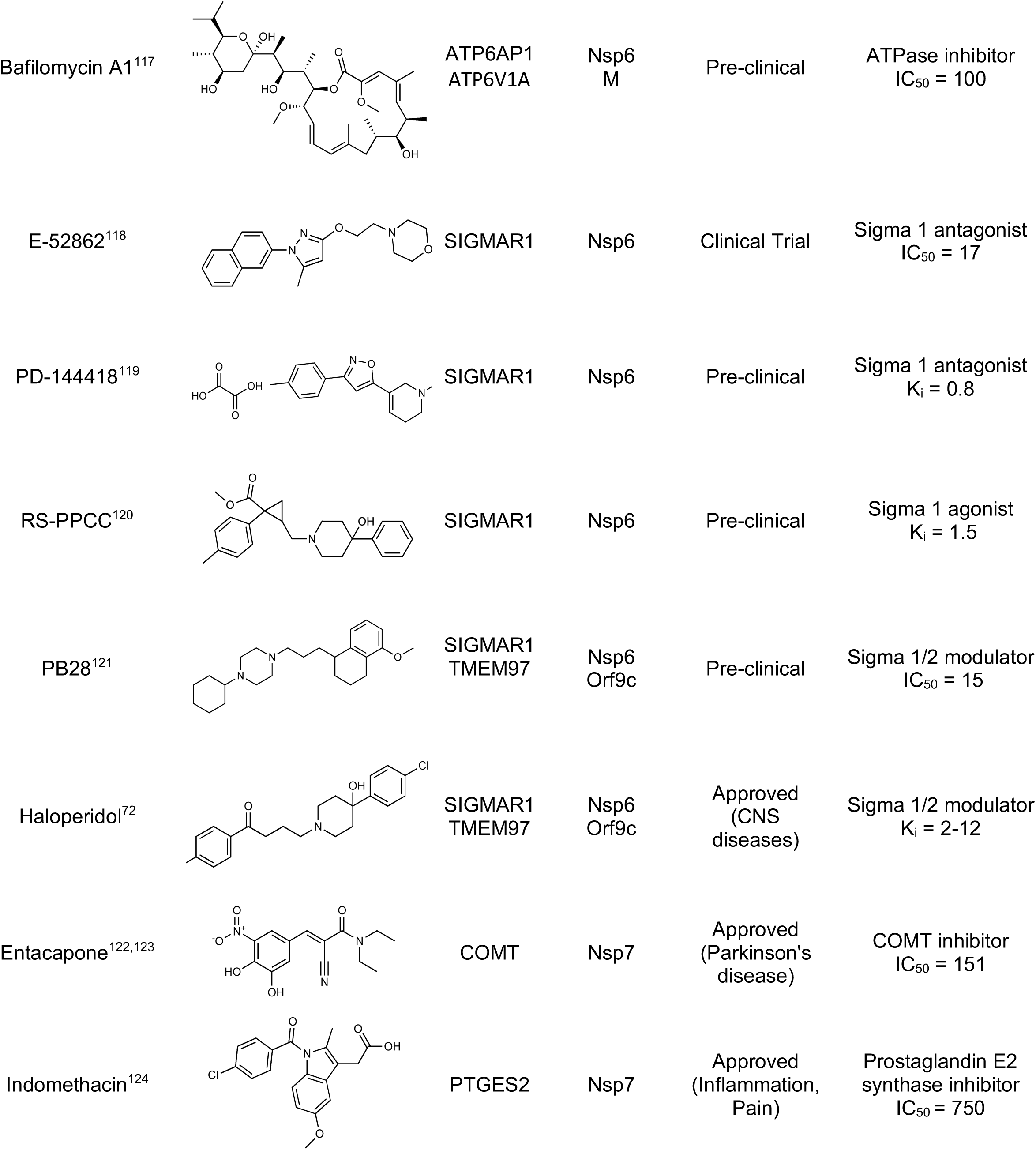
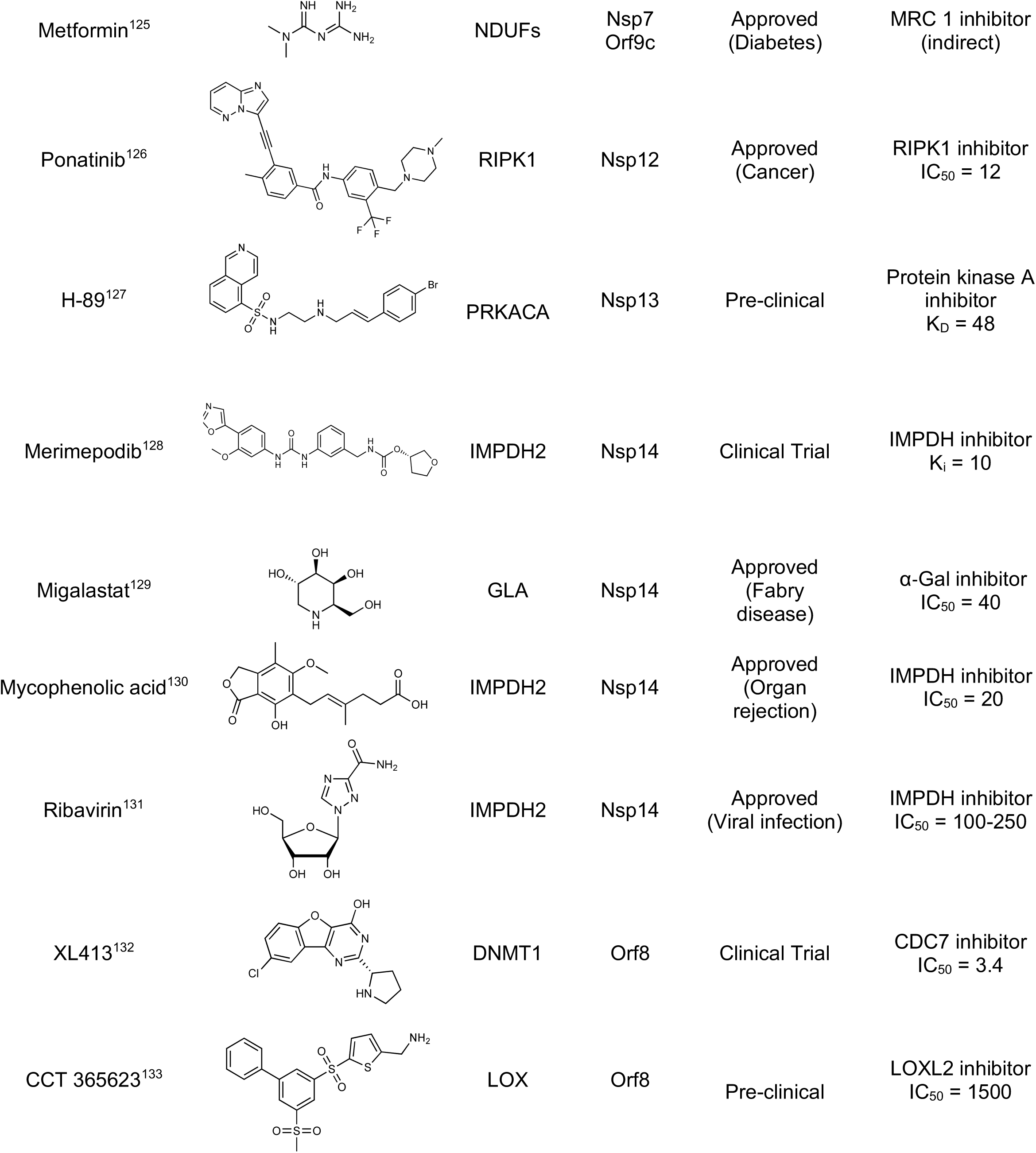
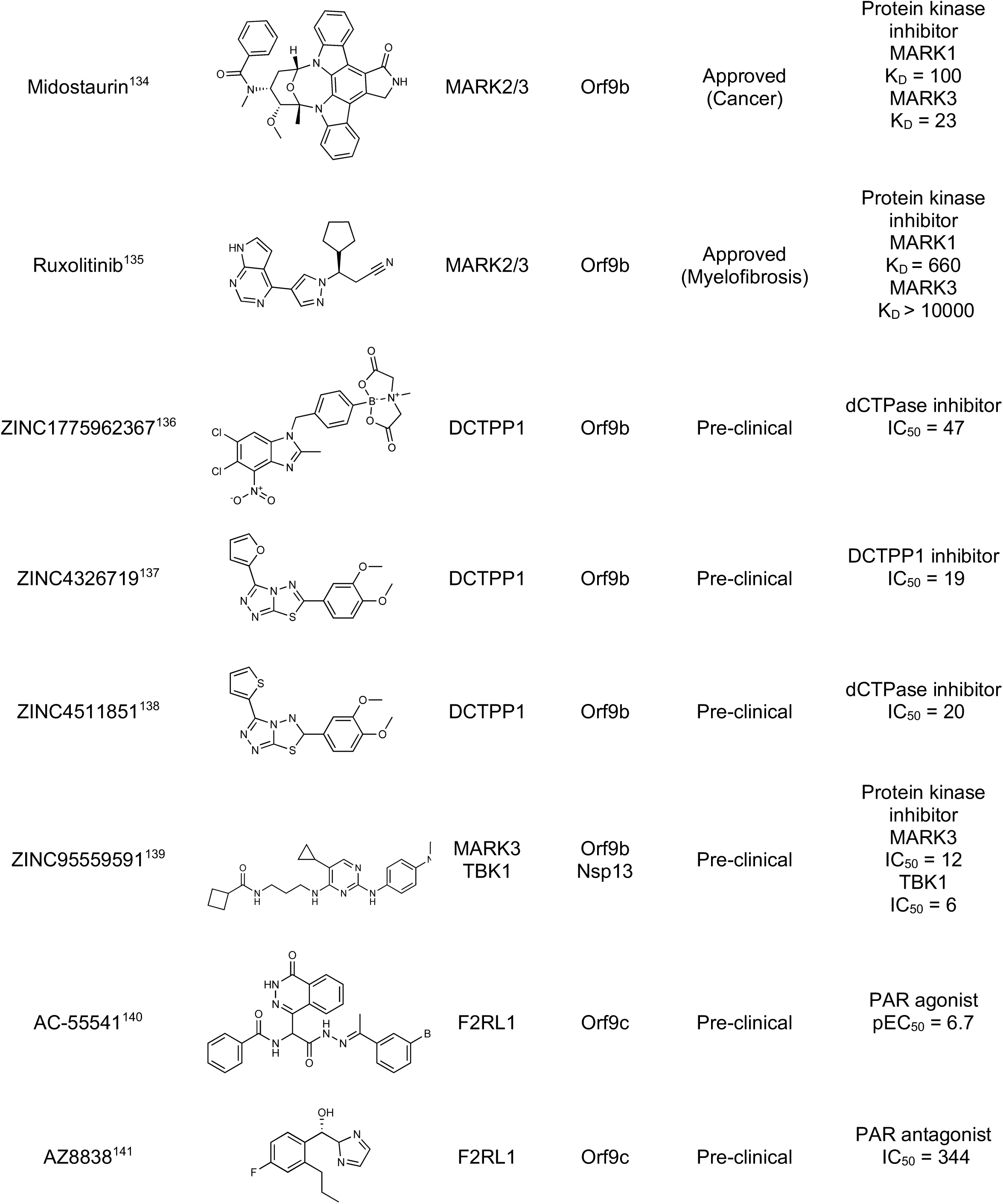
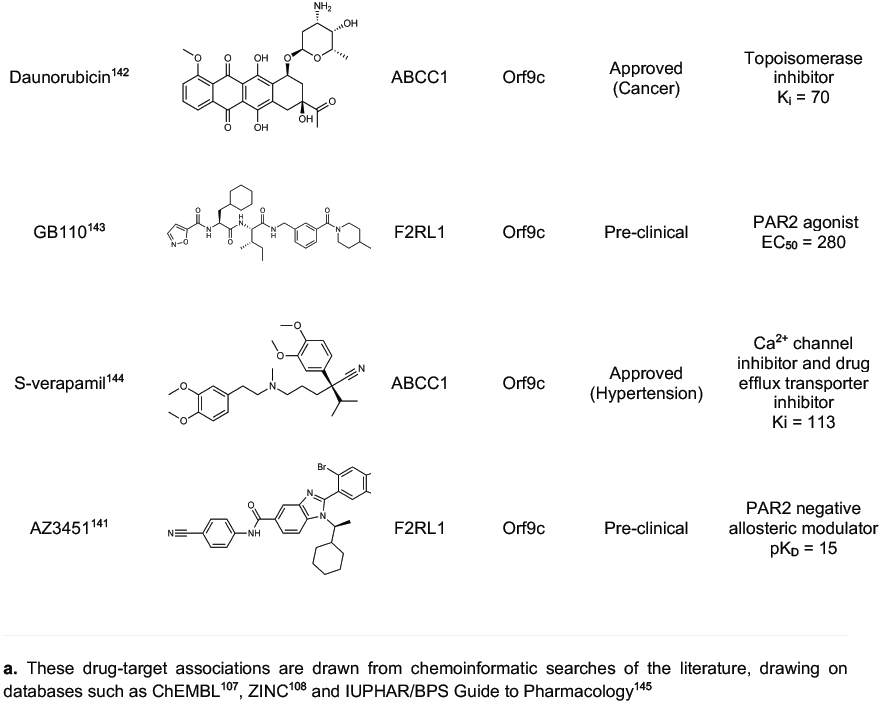
Literature-derived^a^ drugs and reagents that modulate SARS-Cov-2 interactors.

| Compound Name | Compound Structure | Human Gene | Viral Bait | Drug Status | Activity (nM) |
| --- | --- | --- | --- | --- | --- |
| JQ1<sup>109</sup> |  | BRD2/4 | E | Pre-clinical | BRD inhibitor<br>IC<sub>50</sub> = 40-120 |
| RVX-208<sup>109</sup> |  | BRD2/4 | E | Clinical Trial | BRD inhibitor<br>IC<sub>50</sub> = 50-1800 |
| Silmitasertib<sup>110,111</sup> |  | CSNK2A2 | N | Approved (Cancer) | CK2 inhibitor<br>IC<sub>50</sub> = 1 |
| TMCB<sup>112</sup> |  | CSNK2A2 | N | Pre-clinical | Multi-targeted protein kinase inhibitor<br>K<sub>i</sub> = 21 |
| Apicidin<sup>113</sup> |  | HDAC2 | Nsp5 | Pre-clinical | HDAC inhibitor<br>IC<sub>50</sub> = 120 |
| Valproic Acid<sup>114-116</sup> |  | HDAC2 | Nsp5 | Approved (CNS diseases, Cancer) | HDAC2 inhibitor<br>IC<sub>50</sub> = 62,000 |
| Bafilomycin A1<sup>117</sup> |  | ATP6AP1<br>ATP6V1A | Nsp6<br>M | Pre-clinical | ATPase inhibitor<br>IC<sub>50</sub> = 100 |
| E-52862<sup>118</sup> |  | SIGMAR1 | Nsp6 | Clinical Trial | Sigma 1 antagonist<br>IC<sub>50</sub> = 17 |
| PD-144418<sup>119</sup> |  | SIGMAR1 | Nsp6 | Pre-clinical | Sigma 1 antagonist<br>K<sub>i</sub> = 0.8 |
| RS-PPCC<sup>120</sup> |  | SIGMAR1 | Nsp6 | Pre-clinical | Sigma 1 agonist<br>K<sub>i</sub> = 1.5 |
| PB28<sup>121</sup> |  | SIGMAR1<br>TMEM97 | Nsp6<br>Orf9c | Pre-clinical | Sigma 1/2 modulator<br>IC<sub>50</sub> = 15 |
| Haloperidol<sup>72</sup> |  | SIGMAR1<br>TMEM97 | Nsp6<br>Orf9c | Approved<br>(CNS diseases) | Sigma 1/2 modulator<br>K<sub>i</sub> = 2-12 |
| Entacapone<sup>122,123</sup> |  | COMT | Nsp7 | Approved<br>(Parkinson's disease) | COMT inhibitor<br>IC<sub>50</sub> = 151 |
| Indomethacin<sup>124</sup> |  | PTGES2 | Nsp7 | Approved<br>(Inflammation, Pain) | Prostaglandin E2 synthase inhibitor<br>IC<sub>50</sub> = 750 |
| Metformin<sup>125</sup> |  | NDUFs | Nsp7<br>Orf9c | Approved<br>(Diabetes) | MRC 1 inhibitor<br>(indirect) |
| Ponatinib<sup>126</sup> |  | RIPK1 | Nsp12 | Approved<br>(Cancer) | RIPK1 inhibitor<br>IC<sub>50</sub> = 12 |
| H-89<sup>127</sup> |  | PRKACA | Nsp13 | Pre-clinical | Protein kinase A<br>inhibitor<br>K<sub>D</sub> = 48 |
| Merimepodib<sup>128</sup> |  | IMPDH2 | Nsp14 | Clinical Trial | IMPDH inhibitor<br>K<sub>i</sub> = 10 |
| Migalastat<sup>129</sup> |  | GLA | Nsp14 | Approved<br>(Fabry disease) | α-Gal inhibitor<br>IC<sub>50</sub> = 40 |
| Mycophenolic acid<sup>130</sup> |  | IMPDH2 | Nsp14 | Approved<br>(Organ rejection) | IMPDH inhibitor<br>IC<sub>50</sub> = 20 |
| Ribavirin<sup>131</sup> |  | IMPDH2 | Nsp14 | Approved<br>(Viral infection) | IMPDH inhibitor<br>IC<sub>50</sub> = 100-250 |
| XL413<sup>132</sup> |  | DNMT1 | Orf8 | Clinical Trial | CDC7 inhibitor<br>IC<sub>50</sub> = 3.4 |
| CCT 365623<sup>133</sup> |  | LOX | Orf8 | Pre-clinical | LOXL2 inhibitor<br>IC<sub>50</sub> = 1500 |
| Midostaurin<sup>134</sup> | | MARK2/3 | Orf9b | Approved<br>(Cancer) | Protein kinase<br>inhibitor<br>MARK1<br>$K_D = 100$<br>MARK3<br>$K_D = 23$ |
| Ruxolitinib<sup>135</sup> | | MARK2/3 | Orf9b | Approved<br>(Myelofibrosis) | Protein kinase<br>inhibitor<br>MARK1<br>$K_D = 660$<br>MARK3<br>$K_D > 10000$ |
| ZINC1775962367<sup>136</sup> | | DCTPP1 | Orf9b | Pre-clinical | dCTPase inhibitor<br>$IC_{50} = 47$ |
| ZINC4326719<sup>137</sup> | | DCTPP1 | Orf9b | Pre-clinical | DCTPP1 inhibitor<br>$IC_{50} = 19$ |
| ZINC4511851<sup>138</sup> | | DCTPP1 | Orf9b | Pre-clinical | dCTPase inhibitor<br>$IC_{50} = 20$ |
| ZINC95559591<sup>139</sup> | | MARK3<br>TBK1 | Orf9b<br>Nsp13 | Pre-clinical | Protein kinase<br>inhibitor<br>MARK3<br>$IC_{50} = 12$<br>TBK1<br>$IC_{50} = 6$ |
| AC-55541<sup>140</sup> | | F2RL1 | Orf9c | Pre-clinical | PAR agonist<br>$pEC_{50} = 6.7$ |
| AZ8838<sup>141</sup> | | F2RL1 | Orf9c | Pre-clinical | PAR antagonist<br>$IC_{50} = 344$ |
| Daunorubicin<sup>142</sup> | | ABCC1 | Orf9c | Approved<br>(Cancer) | Topoisomerase<br>inhibitor<br>$K_i = 70$ |
| GB110<sup>143</sup> | | F2RL1 | Orf9c | Pre-clinical | PAR2 agonist<br>$EC_{50} = 280$ |
| S-verapamil<sup>144</sup> | | ABCC1 | Orf9c | Approved<br>(Hypertension) | $Ca^{2+}$ channel<br>inhibitor and drug<br>efflux transporter<br>inhibitor<br>$K_i = 113$ |
| AZ3451<sup>141</sup> | | F2RL1 | Orf9c | Pre-clinical | PAR2 negative<br>allosteric modulator<br>$pK_D = 15$ |

a. These drug-target associations are drawn from chemoinformatic searches of the literature, drawing on databases such as ChEMBL<sup>107</sup>, ZINC<sup>108</sup> and IUPHAR/BPS Guide to Pharmacology<sup>145</sup>

**Table 1b.**
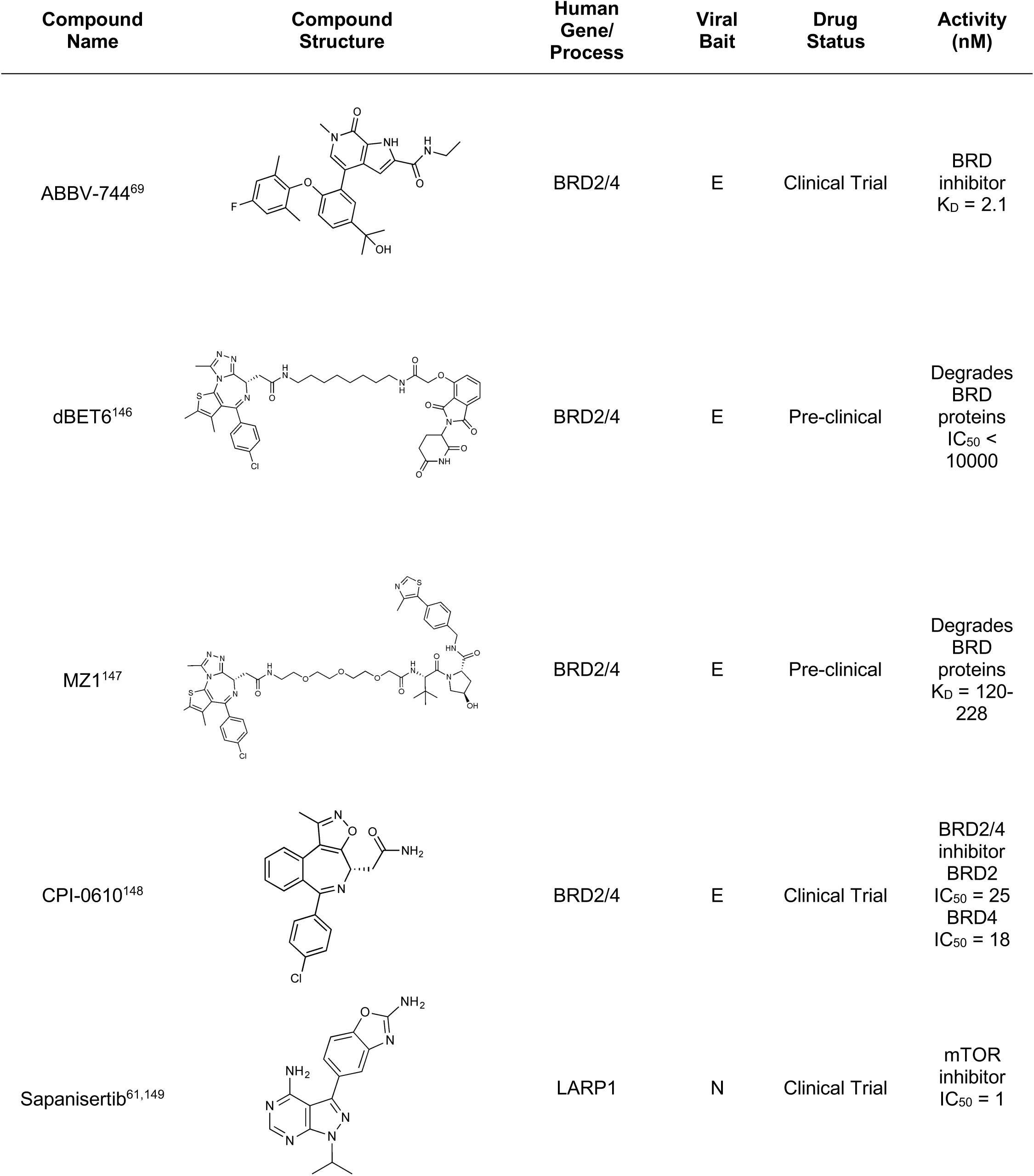
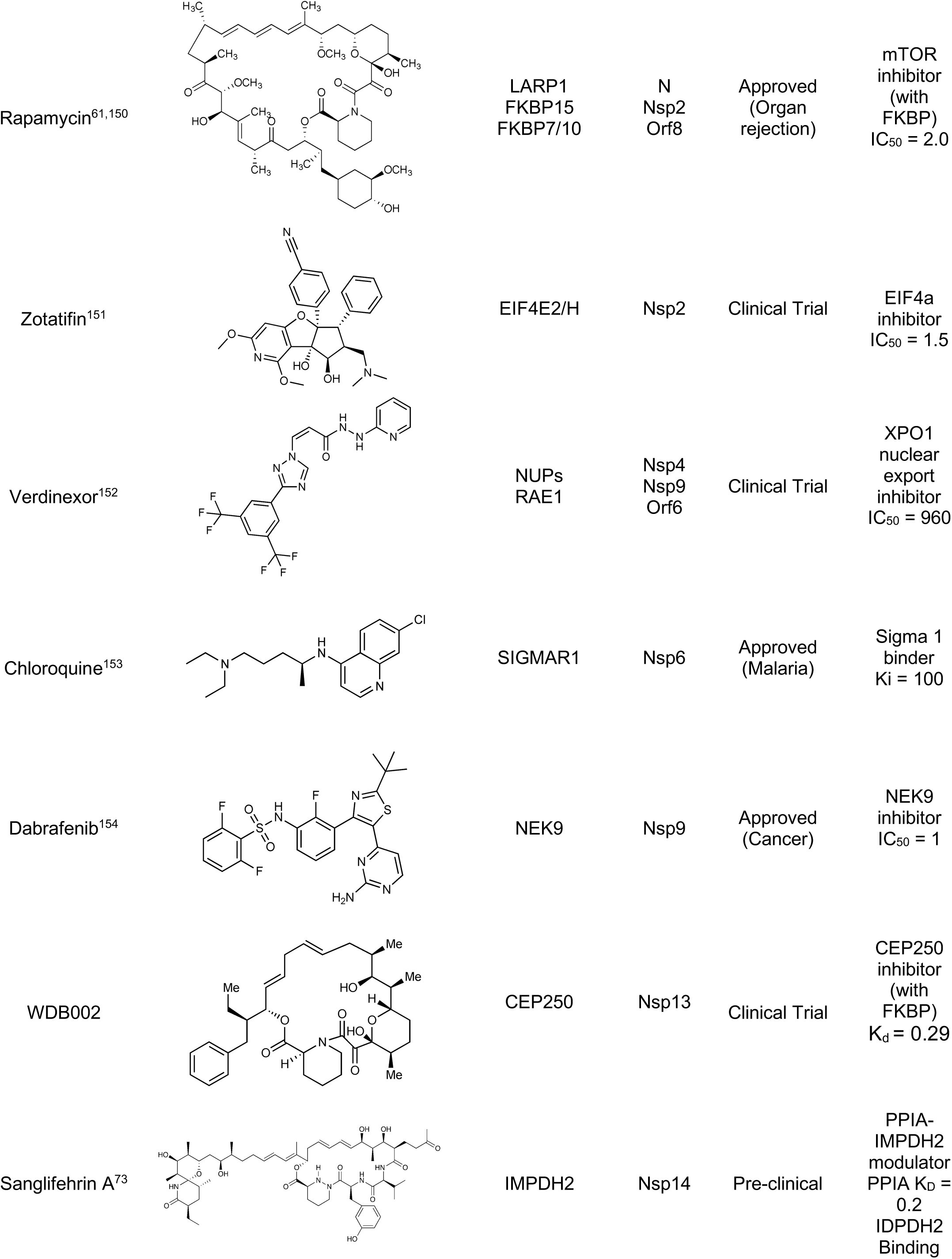
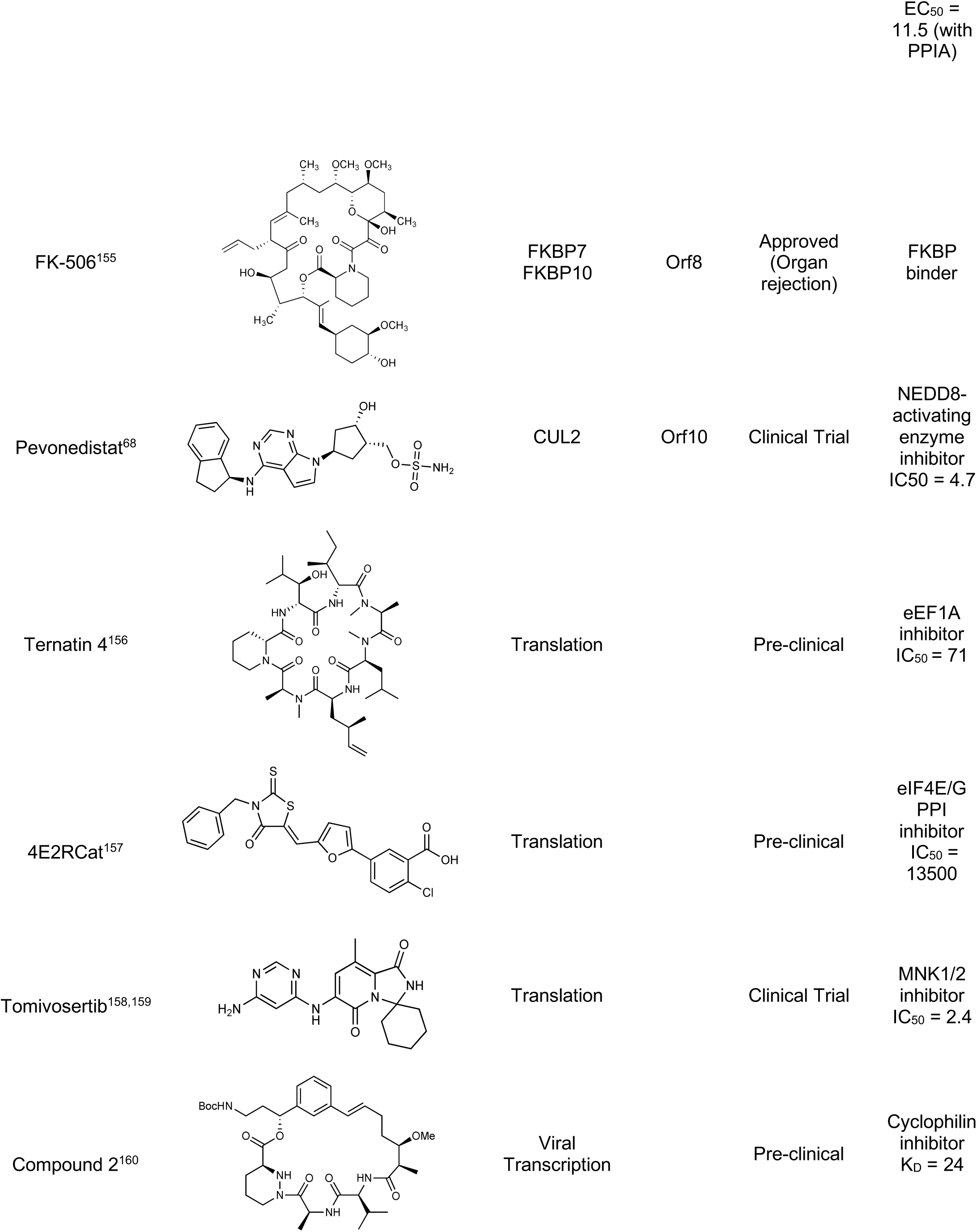
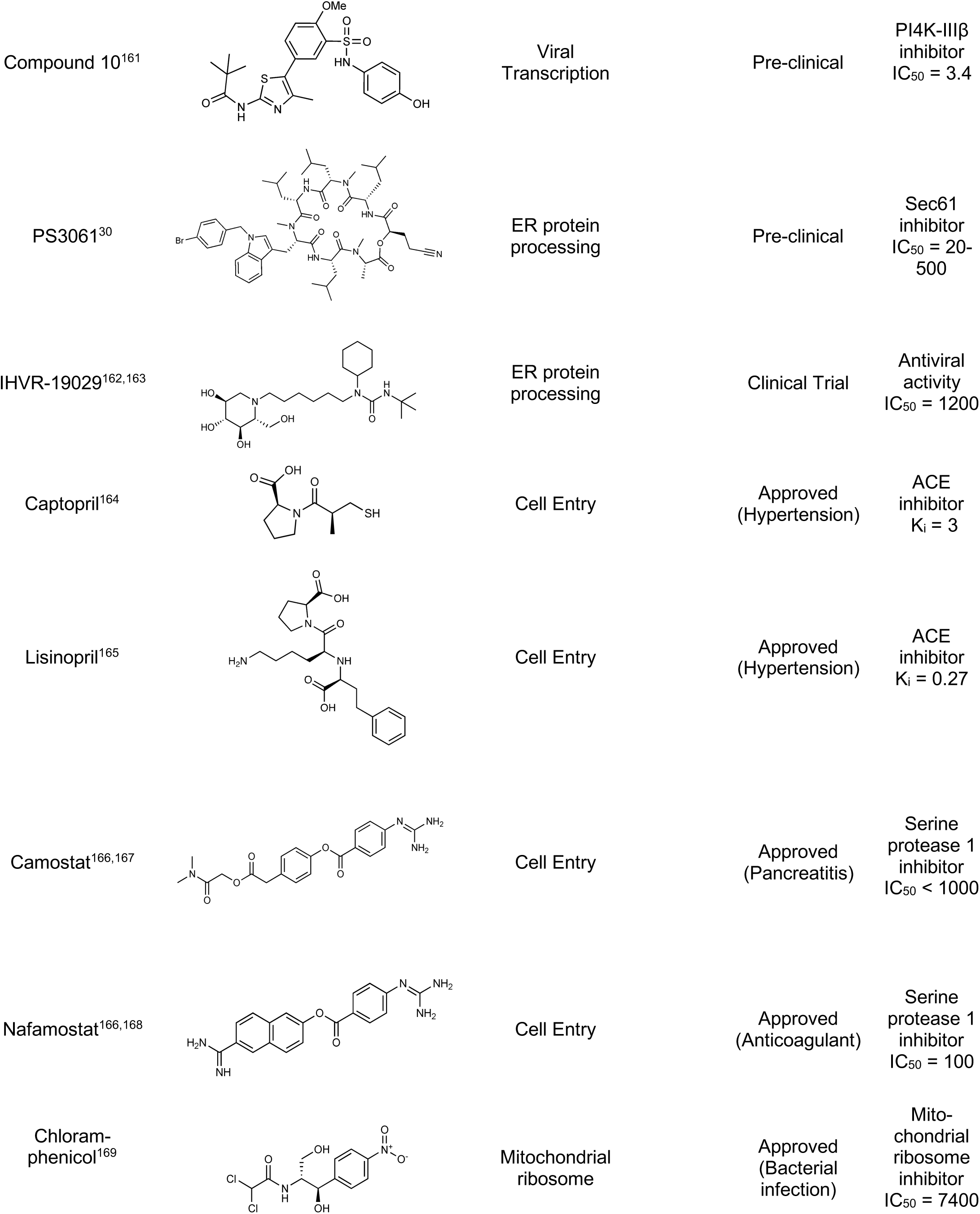
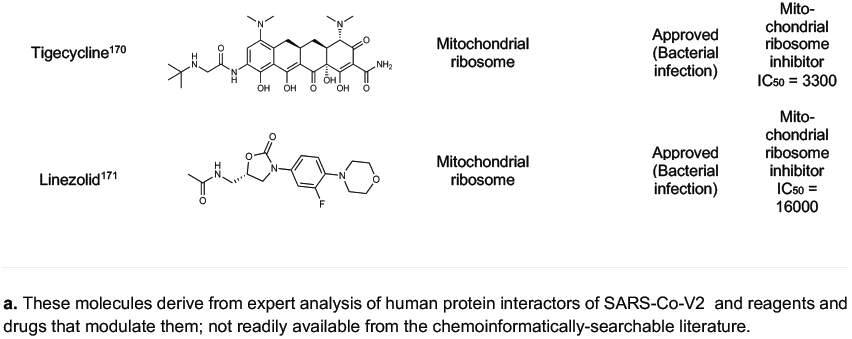
Expert-identified^a^ drugs and reagents that modulate SARS-CoV-2 interactors.

**Figure 5:**
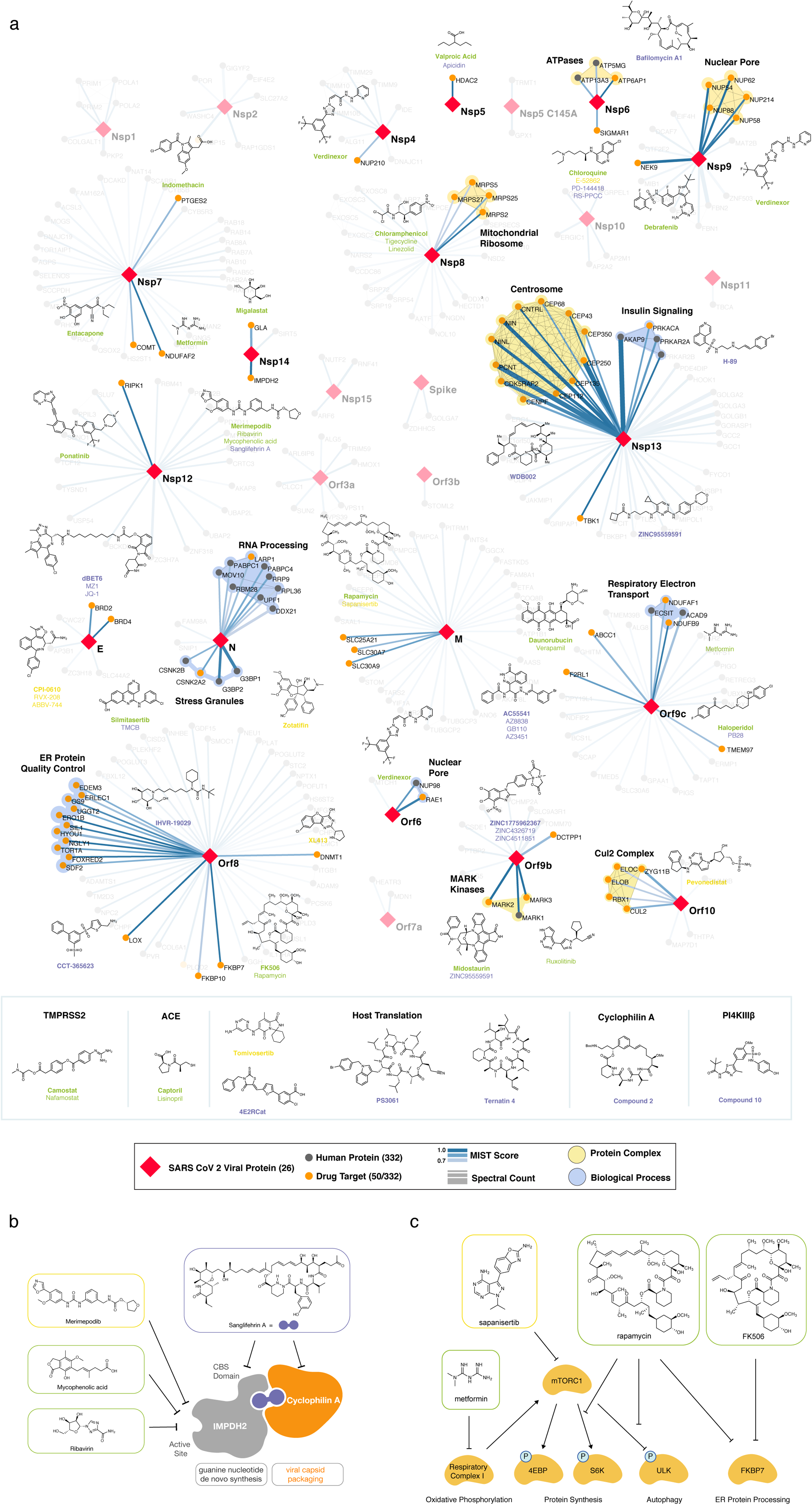
Drug-human target network. **(a)** Significant interactions identified by AP-MS between SARS-CoV-2 baits (red diamonds) and human prey proteins (orange circles) are shown as in **Fig 3**. Chemoinformatic and expert analysis identified FDA approved drugs (green), clinical candidates (yellow), and preclinical candidates (purple) with experimental activities against the host proteins and processes, with representative chemicals shown. (**b**) Inosine Monophosphate Dehydrogenase 2 (IMPDH2) regulates de novo nucleic acid biosynthesis. It is a target for proliferative diseases including cancer^81^ and autoimmune disorders, for instance by the approved drug mycophenolic acid^82^, and as a broad spectrum antiviral by Ribavirin^83^. While Ribavirin has activity against SARS in vitro^84^, it has low tolerability, something that might be addressed by the more selective Merimepodib, which is in phase II clinical trials^85^. (**c**) The mammalian target of Rapamycin (mTOR) pathway is a master regulator of cell proliferation and autophagy, which viruses including Influenza A are known to modulate^86,87^. Several proteins that interact with SARS-CoV-2 baits, including components of the Respiratory complex 1 by Nsp7, Nsp12, and Orf9c, the leucine importer B(0)AT2 (SLC6A15)^88,89^ by Nsp6 and LARP1 by N (not shown). In addition to Rapamycin, the mTOR pathway can be indirectly modulated by metformin, a widely prescribed diabetes drug, and by Sapanisertib, a drug in clinical trials for solid tumors^61^.

There are several mechanistically interesting, and potentially disease-relevant drug-target interactions revealed in the chemoinformatic network (Fig. 5a). Among them, the well-known chemical probe, Bafilomycin A1, is a potent inhibitor of the V1-ATPase, subunits of which interact with Nsp6 and M. Bafilomycin’s inhibition of this cotransporter acts to prevent the acidification of the lysosome, inhibiting autophagy and endosome trafficking pathways, which may impact the viral life-cycle. Similarly, drugs exist to target several well-known epigenetic regulators prominent among the human interactors, including HDAC2, BRD2 and BRD4, which interact with viral proteins nsp5 and E, respectively (Figs. 3 and 5a). The approved drug Valproic acid (an anticonvulsant) and the pre-clinical candidate Apicidin inhibit HDAC2 with affinities of 62 μM and 120 nM, respectively. Clinical compounds ABBV-744 and CPI-0610 act on BRD2/4, with an affinity of 2 nM or 39 nM, respectively -- several preclinical compounds also target bromodomain-containing proteins (Table 1a,b). As a final example, we were intrigued to observe that the SARS-CoV-2 Nsp6 protein interacts with the Sigma receptor, which is thought to regulate ER stress response^71^. Similarly, the Sigma2 receptor interacted with the vial protein orf9. Both Sigma1 and Sigma2 are promiscuous receptors that interact with many non-polar, cationic drugs. We prioritized several of these drugs based on potency or potential disease relevance, including the antipsychotic Haloperidol, which binds in the low nM range to both receptors^72^, and Chloroquine, which is currently in clinical trials for COVID-19 and has mid-nM activity vs the Sigma1 receptor, and low μM activity against the Sigma2 receptor. Because many patients are already treated with drugs that have off-target impact on Sigma receptors, associating clinical outcomes accompanying treatment with these drugs may merit investigation, a point to which we return. Finally, in addition to the druggable host factors, a few of which we have highlighted here, the SARS-CoV-2-human interactome reveals many traditionally “undruggable” targets. Among these, for instance, are components of the centriole such as CEP250, which interacts with the viral Nsp13. Intriguingly, a very recent patent disclosure revealed a natural product, WDB002, that directly and specifically targets CEP250. As a natural product, WDB002 would likely be harder to source than the molecules on which we have focused on here, but may well merit investigation. Similarly, other “undruggable” targets may be revealed to have compounds that could usefully perturb the viral-human interaction network, and act as leads to therapeutics.

Beyond direct interactions, several drug-pathway interactions seemed noteworthy. The human purine biosynthesis enzyme Inosine-5′-monophosphate dehydrogenase (IMPDH2) interacts with the viral protein nsp14. Several chemically diverse compounds inhibit IMPDH2, including the clinically approved mycophenolic acid (20 nM), the approved antiviral drug ribavirin (200 nM), and the investigational new drug Merimepodib (10 nM) (Table 1a). Intriguingly, the preclinical molecule Sanglifehrin A (Table 1b) is known to act as a molecular glue linking IMPDH with cyclophilin A (Fig. 5b)^73^, which itself is implicated in viral capsid packaging, even though it itself is not a human “prey” in the viral-human protein interactome. Similarly, direct viral-human interactions with proteins regulated by the mTORC1 pathway, such as LARP1, and FKBP7, which interact with the viral N and Orf8 proteins, led us to inhibitors of mTORC1, even though that kinase itself is not found to directly interact with a viral protein (Fig. 5c). Sapanisertib and rapamycin are low nM inhibitors of mTORC1, while metformin is an indirect modulator of this protein complex.

## Discussion

We have used affinity purification-mass spectrometry to identify 332 high-confidence SARS-CoV-2-human PPIs. We find the viral proteins connected to a wide array of biological processes, including protein trafficking, translation, transcription and ubiquitination regulation. Using a combination of a systematic chemoinformatic drug search with a pathway centric analysis, we uncovered close to 70 different drugs and compounds, including FDA approved drugs, compounds in clinical trials as well as preclinical compounds, targeting parts of the resulting network. We are currently testing these compounds for antiviral activity and encourage others to do the same as well as extract insights from the map that could have therapeutic value.

More generally, this proteomic/chemoinformatic analysis is not only identifying drug and clinical molecules that might perturb the viral-human interactome, it gives these potentially therapeutic perturbations a mechanistic context. Among those that may be infection relevant are the inhibition of lysosomal acidification and trafficking with Bafilomycin A1, via inhibition of V-ATPase^74^, and modulation of the ER stress and the protein unfolding response pathway by targeting the Sigma1 and Sigma2 receptor by drugs like haloperidol (Fig. 5a, Tables 1a,b). Indeed, several of the human proteins in the interactome are targeted by drugs that have emerged phenotypically as candidate therapeutics for treating COVID-19, such as chloroquine^75,76^. While we do not pretend to have identified the molecular basis of chloroquine’s putative activity, we do note that this drug targets the Sigma1 and Sigma2 receptors at mid-nM and low μM concentrations, respectively. Similarly, antibiotics like azithromycin have also been mooted as treatments for COVID-19. While this too remains to be demonstrated, we note that Azithromycin has off-target activity against human mitochondrial ribosomes, components of which interact with the SARS-CoV-2 Nsp8 protein (MRPS5, MRPS27, MRPS2, and MRPS25). Other antibiotics that also have an off-target effect on mitochondrial ribosomes, such as chloramphenicol, tigecycline, and Linezolid^77,78^ may also merit study for efficacy. Indeed, this logic may be extended. Many COVID-19 patients will be on the drugs identified here, treating pre-existing conditions. It may be useful to correlate clinical outcomes with the taking of these drugs, cross-referencing with the networks described here. In some senses, this is already occurring phenomenologically, leading to concerns about ACE inhibitors such as captopril and enalapril, and for NSAIDs. What this study provides is a systematic schema for clinical/drug associations going forward, giving them a mechanistic context that allows investigators to seek them directly.

Systematic validation using genetic-based approaches^79,80^ will be key to determine the functional relevance of these interactions and if the human proteins are being used by the virus or are fighting off infection, information that would inform future pharmacological studies. *It is important to note that pharmacological intervention with the agents we identified in this study could be either detrimental or beneficial for infection.* For instance, the HDAC2 inhibitors may compound the potential action of the Nsp5 protease to hydrolyze this human protein. Future work will involve generation of protein-protein interaction maps in different human cell types, as well as bat cells, and the study of related coronaviruses including SARS-CoV, MERS-CoV and the less virulent OC43^5^, data that will allow for valuable cross-species and viral evolution studies. Targeted biochemical and structural studies will also be crucial for a deeper understanding of the viral-host complexes, which will inform more targeted drug design.

Along with SARS-CoV-2, we have previously utilized global affinity purification-mass spectrometry (AP-MS) analysis to map the host-pathogen interfaces of a number of human pathogens including Ebola^22^, Dengue^30^, Zika^30^, Herpesvirus^29^, Hepatitis C^28^, Tuberculosis^27^, Chlamydia^26^, Enteroviruses^25^, HIV^19^, HPV^24^, and West Nile Fever^23^. Excitingly, we have uncovered both shared and unique mechanisms in which these pathogens co-opt the host machinery during the course of infection. Although host-directed therapy is often not explored for combatting pathogenic infections, it would be interesting to use this information to identify host factors that could serve as targets that would harbor pan-pathogenic activity so that when the next virus undergoes zoonosis, we will have treatment options available.

## Supporting information

Supplementary Information Discussion

DNA Sequences

## Abbreviations

HC-PPIs: High confidence protein-protein interactions
PPIs: protein-protein interaction
AP-MS: affinity purification-mass spectrometry
COVID-19: Coronavirus Disease-2019
ACE2: angiotensin converting enzyme 2
Orf: open reading frame
Nsp3: papain-like protease
Nsp5: main protease
Nsp: nonstructural protein
TPM: transcripts per million

Supplementary Information is available for this paper.

Reprints and permissions information is available at www.nature.com/reprints

The authors have not filed for patent protection on the SARS-CoV-2 host interactions or the use of predicted drugs for treating COVID-19 to ensure all the information is freely available to accelerate the discovery of a treatment.

## ACKNOWLEDGEMENTS

We thank Joshua Sarlo for his artistic contributions to the manuscript.

This research was funded by grants from the National Institutes of Health (P50AI150476, U19AI135990, U19AI135972, R01AI143292, R01AI120694, P01A1063302, and R01AI122747 to N.J.K.; R35GM122481 to B.K.S.; 1R01CA221969 and 1R01CA244550 to K.M.S.; K08HL124068 to J.S.C.; 1F32CA236347-01 to J.E.M.); funding from F. Hoffmann-La Roche and Vir Biotechnology and gifts from The Ron Conway Family. K.M.S is an investigator of the Howard Hughes Medical Institute.

## MATERIALS AND METHODS

### Genome annotation

The genbank sequence for SARS-CoV-2 isolate 2019-nCoV/USA-WA1/2020, accession MN985325, was downloaded on January 24, 2020. In total, we identified 29 open reading frames and proteolytically mature proteins encoded by SARS-CoV-2^1,16^. Proteolytic products resulting from Nsp3 and Nsp5-mediated cleavage of the Orf1a / Orf1ab polyprotein were predicted based on the protease specificity of SARS-CoV proteases^90^, and 16 predicted nonstructural proteins (Nsps) were subsequently cloned (Nsp1-Nsp16). For the proteases Nsp3 (papain-like / Plpro) and Nsp5 (3Clike / 3CLpro), we also designed catalytic dyad/triad mutants: Nsp3 C857A^91^ and Nsp5 C145A ^92,93^. Open reading frames at the 3’ end of the viral genome annotated in the original genbank file included 4 Structural proteins: S, E, M, N, and the additional open reading frames Orf3a, Orf6, Orf7a, Orf8, and Orf10. Based on analysis of open reading frames in the genome and comparisons with other annotated SARS-CoV open reading frames, we annotated a further four open reading frames: Orf3b, Orf7b, Orf9b, and Orf9c.

### Cell culture

HEK293T cells were cultured in Dulbecco’s Modified Eagle’s Medium (Corning) supplemented with 10% Fetal Bovine Serum (Gibco, Life Technologies) and 1% Penicillin-Streptomycin (Corning) and maintained at 37°C in a humidified atmosphere of 5% CO_2_.

### Transfection

For each affinity purification, ten million HEK293T cells were plated per 15-cm dish and transfected with up to 15 μg of individual Strep-tagged expression constructs after 20-24 hours. Total plasmid was normalized to 15 μg with empty vector and complexed with PolyJet Transfection Reagent (SignaGen Laboratories) at a 1:3 μg:μl ratio of plasmid to transfection reagent based on manufacturer’s recommendations. After more than 38 hours, cells were dissociated at room temperature using 10 ml Dulbecco’s Phosphate Buffered Saline without calcium and magnesium (D-PBS) supplemented with 10 mM EDTA for at least 5 minutes and subsequently washed with 10 ml D-PBS. Each step was followed by centrifugation at 200 xg, 4°C for 5 minutes. Cell pellets were frozen on dry ice and stored at - 80°C. At least three biological replicates were independently prepared for affinity purification.

### Affinity purification

Frozen cell pellets were thawed on ice for 15-20 minutes and suspended in 1 ml Lysis Buffer [IP Buffer (50 mM Tris-HCl, pH 7.4 at 4°C, 150 mM NaCl, 1 mM EDTA) supplemented with 0.5% Nonidet P 40 Substitute (NP40; Fluka Analytical) and cOmplete mini EDTA-free protease and PhosSTOP phosphatase inhibitor cocktails (Roche)]. Samples were then frozen on dry ice for 10-20 minutes and partially thawed at 37°C before incubation on a tube rotator for 30 minutes at 4°C and centrifugation at 13,000 xg, 4°C for 15 minutes to pellet debris. After reserving 50 μl lysate, up to 48 samples were arrayed into a 96-well Deepwell plate for affinity purification on the KingFisher Flex Purification System (Thermo Scientific) as follows: MagStrep “type3” beads (30 μl; IBA Lifesciences) were equilibrated twice with 1 ml Wash Buffer (IP Buffer supplemented with 0.05% NP40) and incubated with 0.95 ml lysate for 2 hours. Beads were washed three times with 1 ml Wash Buffer and then once with 1 ml IP Buffer. To directly digest bead-bound proteins as well as elute proteins with biotin, beads were manually suspended in IP Buffer and divided in half before transferring to 50 μl Denaturation-Reduction Buffer (2 M urea, 50 mM Tris-HCl pH 8.0, 1 mM DTT) and 50 μl 1x Buffer BXT (IBA Lifesciences) dispensed into a single 96-well KF microtiter plate, respectively. Purified proteins were first eluted at room temperature for 30 minutes with constant shaking at 1,100 rpm on a ThermoMixer C incubator. After removing eluates, on-bead digestion proceeded (below). Strep-tagged protein expression in lysates and enrichment in eluates were assessed by western blot and silver stain, respectively. The KingFisher Flex Purification System was placed in the cold room and allowed to equilibrate to 4°C overnight before use. All automated protocol steps were performed using the slow mix speed and the following mix times: 30 seconds for equilibration/wash steps, 2 hours for binding, and 1 minute for final bead release. Three 10 second bead collection times were used between all steps.

### On-bead digestion

Bead-bound proteins were denatured and reduced at 37°C for 30 minutes and after bringing to room temperature, alkylated in the dark with 3 mM iodoacetamide for 45 minutes and quenched with 3 mM DTT for 10 minutes. Proteins were then incubated at 37°C, initially for 4 hours with 1.5 μl trypsin (0.5 μg/μl; Promega) and then another 1-2 hours with 0.5 μl additional trypsin. To offset evaporation, 15 μl 50 mM Tris-HCl, pH 8.0 were added before trypsin digestion. All steps were performed with constant shaking at 1,100 rpm on a ThermoMixer C incubator. Resulting peptides were combined with 50 μl 50 mM Tris-HCl, pH 8.0 used to rinse beads and acidified with trifluoroacetic acid (0.5% final, pH < 2.0). Acidified peptides were desalted for MS analysis using a BioPureSPE Mini 96-Well Plate (20mg PROTO 300 C18; The Nest Group, Inc.) according to standard protocols.

### Mass spectrometry data acquisition and analysis

Samples were re-suspended in 4% formic acid, 2% acetonitrile solution, and separated by a reversed-phase gradient over a nanoflow C18 column (Dr. Maisch). Each sample was analyzed on two different mass spectrometers. First, a 75 min acquisition, in which peptides were directly injected via a Easy-nLC 1200 (Thermo) into a Q-Exactive Plus mass spectrometer (Thermo), with all MS1 and MS2 spectra collected in the orbitrap. For all acquisitions, QCloud was used to control instrument longitudinal performance during the project^94^. All proteomic data was searched against the human proteome (uniprot reviewed sequences downloaded February 28th, 2020), EGFP sequence, and the SARS-CoV-2 protein sequences using the default settings for MaxQuant^95,96^. Detected peptides and proteins were filtered to 1% false discovery rate in MaxQuant, and identified proteins were then subjected to protein-protein interaction scoring with both SAINTexpress^20^ and MiST^19,97^. We applied a two step filtering strategy to determine the final list of reported interactors which relied on two different scoring stringency cutoffs. In the first step, we chose all protein interactions that possess a MiST score ≥ 0.7, a SAINTexpress BFDR ≤ 0.05 and an average spectral count ≥ 2. For all proteins that fulfilled these criteria we extracted information about stable protein complexes they participate in from the CORUM^98^ database of known protein complexes. In the second step we then relaxed the stringency and recovered additional interactors that (1) form complexes with interactors determined in filtering step 1 and (2) fulfill the following criteria: MiST score ≥ 0.6, SAINTexpress BFDR ≤ 0.05 and average spectral counts ≥ 2. Proteins that fulfilled filtering criteria in either step 1 or step 2 were considered to be HC-PPIs and visualized with cytoscape^99^. Using this filtering criteria, nearly all of our baits recovered a number of HC-PPIs in close alignment with previous datasets reporting an average of ∼6 PPIs per bait^100^. However, for a subset of baits (Orf8, Nsp8, Nsp13, and orf9c) we observed a much higher number of PPIs passing these filtering criteria. For these four baits, the MiST scoring was instead performed using an larger in-house database of 87 baits that were prepared and processed in an analogous manner to this SARS-CoV-2 dataset. This was done to provide a more comprehensive collection of baits for comparison, to minimize the classification of non-specifically binding background proteins as HC-PPIs. All mass spectrometry raw data and search results files have been deposited to the ProteomeXchange Consortium via the PRIDE partner repository with the dataset identifier PXD018117^101,102^. PPI networks have also been uploaded to NDEx.

### Gene Ontology Over-representation Analysis

The targets of each bait were tested for enrichment of Gene Ontology (GO Biological Process) terms. The over-representation analysis (ORA) was performed using the enricher function of clusterProfiler package in R with default parameters. The gene ontology terms were obtained from the c5 category of Molecular Signature Database (MSigDBv6.1). Significant GO terms (1% FDR) were identified and further refined to select non-redundant terms. In order to select non-redundant gene sets, we first constructed a GO term tree based on distances (1-Jaccard Similarity Coefficients of shared genes) between the significant terms. The GO term tree was cut at a specific level (h=0.99) to identify clusters of non-redundant gene sets. For a bait with multiple significant terms belonging to the same cluster, we selected the broadest term i.e. largest gene set size.

### Virus Interactome Similarity Analysis

Interactome similarity was assessed by comparing the number of shared human interacting proteins between pathogen pairs, using a hypergeometric test to calculate significance. The background gene set for the test consisted of all unique proteins detected by mass spectrometry across all pathogens (N=10,181 genes).

### Orf6 peptide modeling

The proposed interaction between Orf6 and the NUP98-RAE1 complex was modeled in PyRosetta 4^103^ (release v2020.02-dev61090) using the crystal structure of Vesicular stomatitis virus matrix (M) protein bound to NUP98-RAE1 as a template^66^ (PDB 4OWR downloaded from the PDB-REDO server^104^). The M protein chain (C) was truncated after residue 54 to restrict the model to the putative interaction motif in Orf6 (M protein residues 49-54, sequence DEMDTH). These residues were mutated to the Orf6 sequence, QPMEID, using the *mutate_residue* function in the module *pyrosetta.toolbox*, without repacking at this initial step. After all six residues were mutated, the full model was relaxed to a low energy conformation using the *FastRelax* protocol in the module *pyrosetta.rosetta.protocols.relax. FastRelax* was run with constraints to starting coordinates and scored with the ref2015 score function. The resulting model was inspected for any large energetic penalties associated with the modeled peptide residues or those NUP98 and RAE1 residues interacting with the peptide, and was found to have none. The model was visualized in PyMOL (The PyMOL Molecular Graphics System, Version 2.3.4 Schrödinger, LLC.).

### CUL2^ZYG11B^ homology model generation

The CRL2^ZYG11B^ homology model was built with Swissmodel^105^ and Modeller^106^ by using the homology template of each domain from PDB database (PDB codes: 4b8o, 5jh5,1g03, and 6r7n). The ZYG11B model has two structured domains: a leucine rich repeat (LRR) and Armadillo Repeat (ARM) at the N and C-terminus respectively. The linker between each domain was not modelled due to high flexibility between residues 32 to 49 and residues 304 to 322. Putative protein interaction surfaces on ZYG11B were modelled based on contiguous surface exposed residues that are conserved in ZYG11B orthologues from *C. elegans to H. sapiens* (ZY11B_HUMAN; ZY11B_MOUSE; F1M8P2_RAT; ZYG11_XENLA; ZYG11_DANRE; ZYG11_CAEEL) and located at typical substrate binding sites in the homologous structures of LRR and ARM domain co-complexes.

### Alignment of Protein E and Histone H2A

In order to align protein E and histone H2A, the structure of the protein E SARS-CoV homolog (PDB ID: 2MM4) was compared to the human nucleosome structure (PDB ID: 6K1K). Protein E was structurally aligned to the histone subunits using Pymol’s “*align”* function (https://pymolwiki.org/index.php/Align). *Align* performs a sequence alignment followed by a structural superposition, and then carries out zero or more cycles of refinement in order to reject structural outliers found during the fit. The best superposition was obtained for H2A residues 49-60 & 63-70 and Protein E residues 25-44 at an RMSD of 2.8Å, as reported in Figure 4d.

### Chemoinformatic Analysis of SARS-CoV2 Interacting Partners

To identify drugs and reagents that modulate the 332 host factors interacting with SARS-CoV-2-HEK293T (MiST >= 0.70), we used two approaches: 1) a chemoinformatic analysis of open-source chemical databases and 2) a target- and pathway-specific literature search, drawing on specialist knowledge within our group. Chemoinformatically, we retrieved 2,472 molecules from the IUPHAR/BPS Guide to Pharmacology (2020-3-12) that interacted with 30 human “prey” proteins (38 approved, 71 in clinical trials), and found 10,883 molecules (95 approved, 369 in clinical trials) from the ChEMBL25 database^107^ (Supplementary Tables 5, 6). For both approaches, molecules were prioritized on their FDA approval status, activity at the target of interest better than 1 μM, and commercial availability, drawing on the ZINC database^108^. FDA approved molecules were prioritized except when clinical candidates or preclinical research molecules had substantially better selectivity or potency on-target. In some cases, we considered molecules with indirect mechanisms of action on the general pathway of interest based solely on literature evidence (e.g., captopril modulates ACE2 indirectly via its direct interaction with Angiotensin Converting Enzyme, ACE). Finally, we predicted 6 additional molecules (2 approved, 1 in clinical trials) for proteins with MIST scores between 0.7-0.6 to viral baits (Supplemental Tables 3 and 4). Complete methods can be found here (www.github.com/momeara/BioChemPantry/vignette/COVID19).

## SUPPLEMENTARY INFORMATION

Supplementary table 1: Scoring results for all baits and all proteins

Supplementary table 2: SARS-CoV 2 high confidence interactors

Supplementary table 3: Literature-derived drugs and reagents that modulate SARS-Cov-2 interactors. Drug-target associations drawn from chemoinformatic searches of the literature, including information about purchasability

Supplementary table 4: Expert-identified drugs and reagents that modulate SARS-CoV-2 interactors. Drug-target associations drawn from expert knowledge of human protein interactors of SARS-Co-V2 and reagents and drugs that modulate them; not readily available from the chemoinformatically-searchable literature

Supplementary table 5: Raw chemical associations to prey proteins IUPHAR/BPS Guide to Pharmacology (2020-3-12)

Supplementary table 6: Raw chemical associations to prey proteins ChEMBL25

Supplementary Methods: Computational methods used to propagate tables and supplemental figures

Supplementary Discussion: In depth look at the SARS-CoV-2 individual bait subnetworks

