## Extended Data Figures for "A SARS-CoV-2-Human Protein-Protein Interaction Map Reveals Drug Targets and Potential Drug-Repurposing"

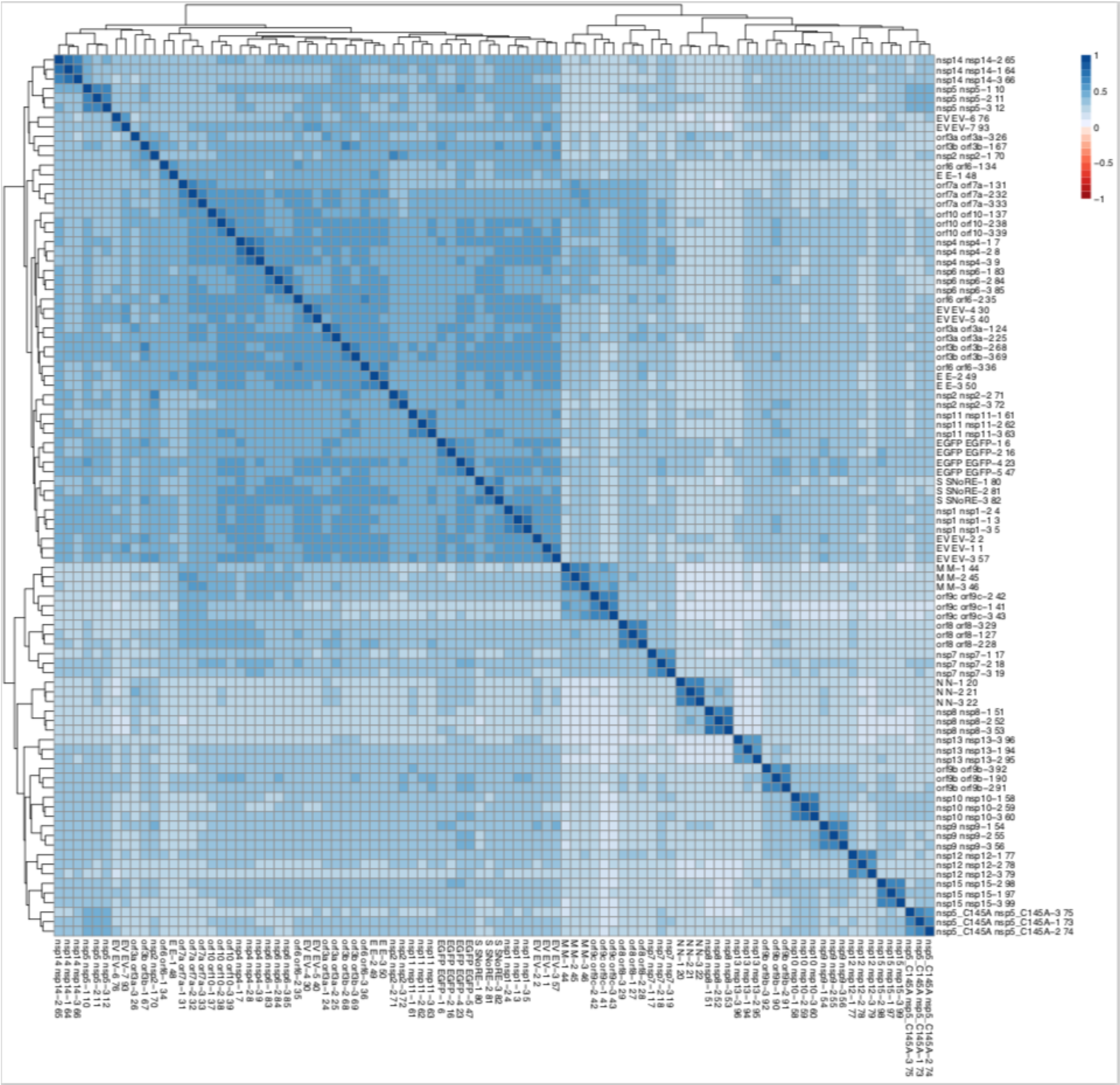


**Extended Data Figure 1. Clustering analysis of AP-MS dataset reveals biological replicates of individual baits are well correlated.** All MS runs were compared and clustered using artMS (David Jimenez-Morales, Alexandre Rosa Campos and John Von Dollen. (2019). artMS: Analytical R tools for Mass Spectrometry. R package version 1.3.9. https://github.com/biodavidjm/artMS). This figure depicts all Pearson's pairwise correlations between MS runs, and is clustered according to similar correlation patterns.

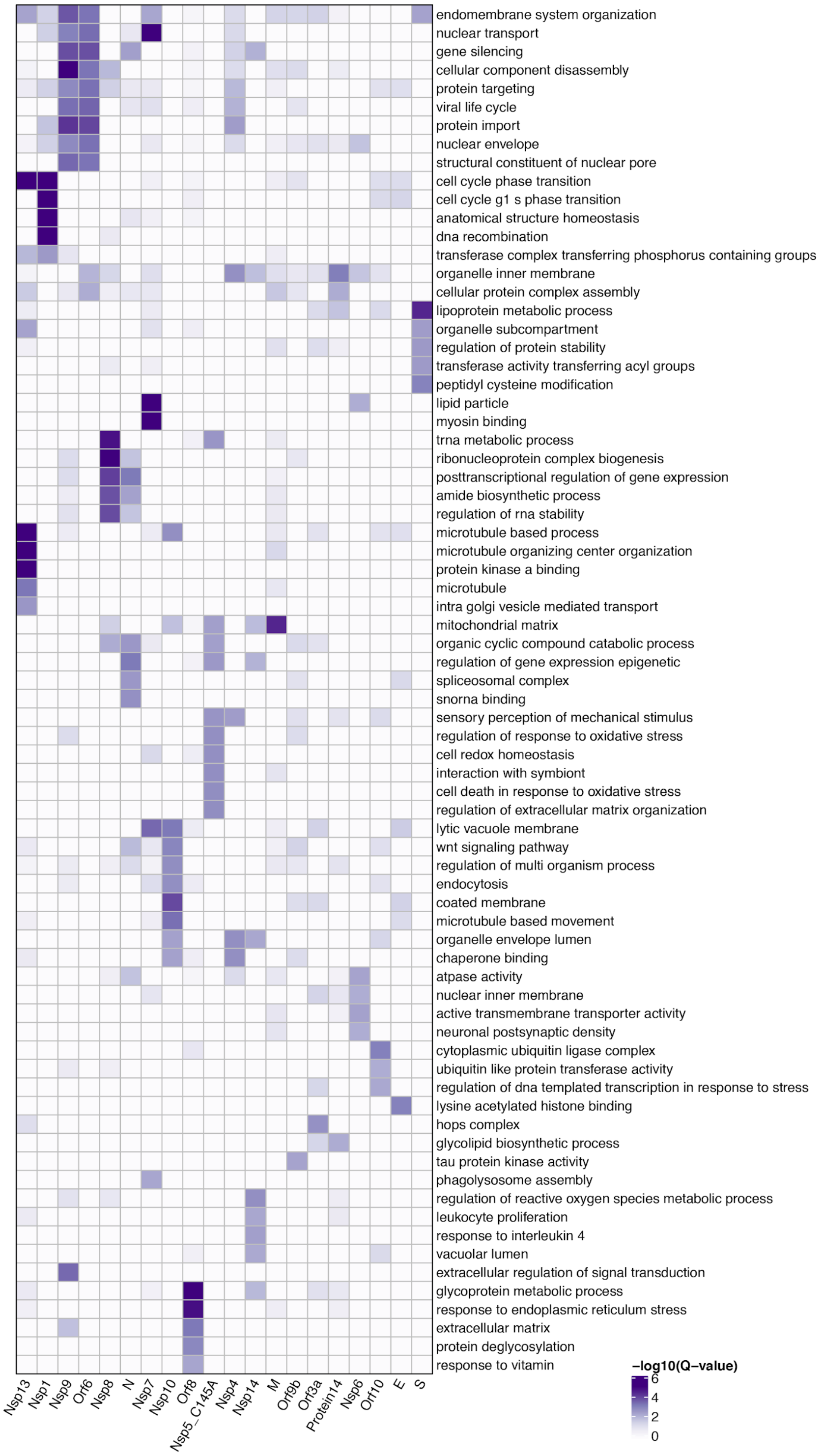


**Extended Data Figure 2. Gene Ontology Biological Process Enrichments for SARS-CoV-2 Host Factors.** We performed GO biological process enrichments (Methods) for the host factors identified as binding to each SARS-CoV-2 viral protein and represent here the top 5 most significant terms for each viral protein.

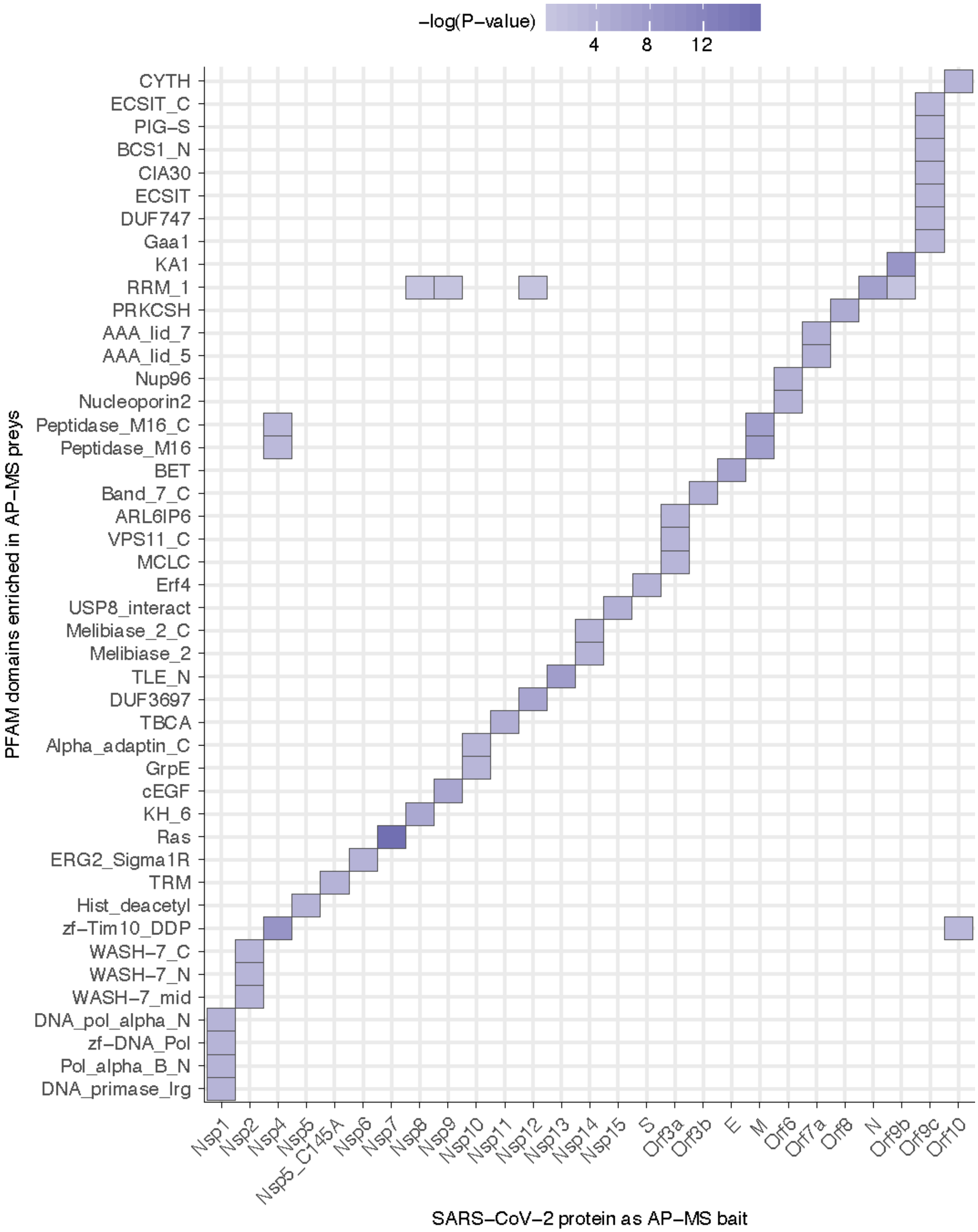


**Extended Data Figure 3. Pfam Protein Families Enrichments for SARS-CoV-2 Host Factors.** The enrichment of individual PFAM domains was calculated using a hypergeometric test where success is defined as the number of domains, and the number of trials is the number of individual preys pulled-down with each viral bait. The population values were the numbers of individual PFAM domains in the human proteome. To make sure that the p-values that signify enrichment were meaningful, we only considered PFAM domains that have been pulled-down at least three times with any SARS-CoV-2 protein, and which occur in the human proteome at least five times. Here, we show PFAM domains with the lowest p-value for a given viral bait protein.

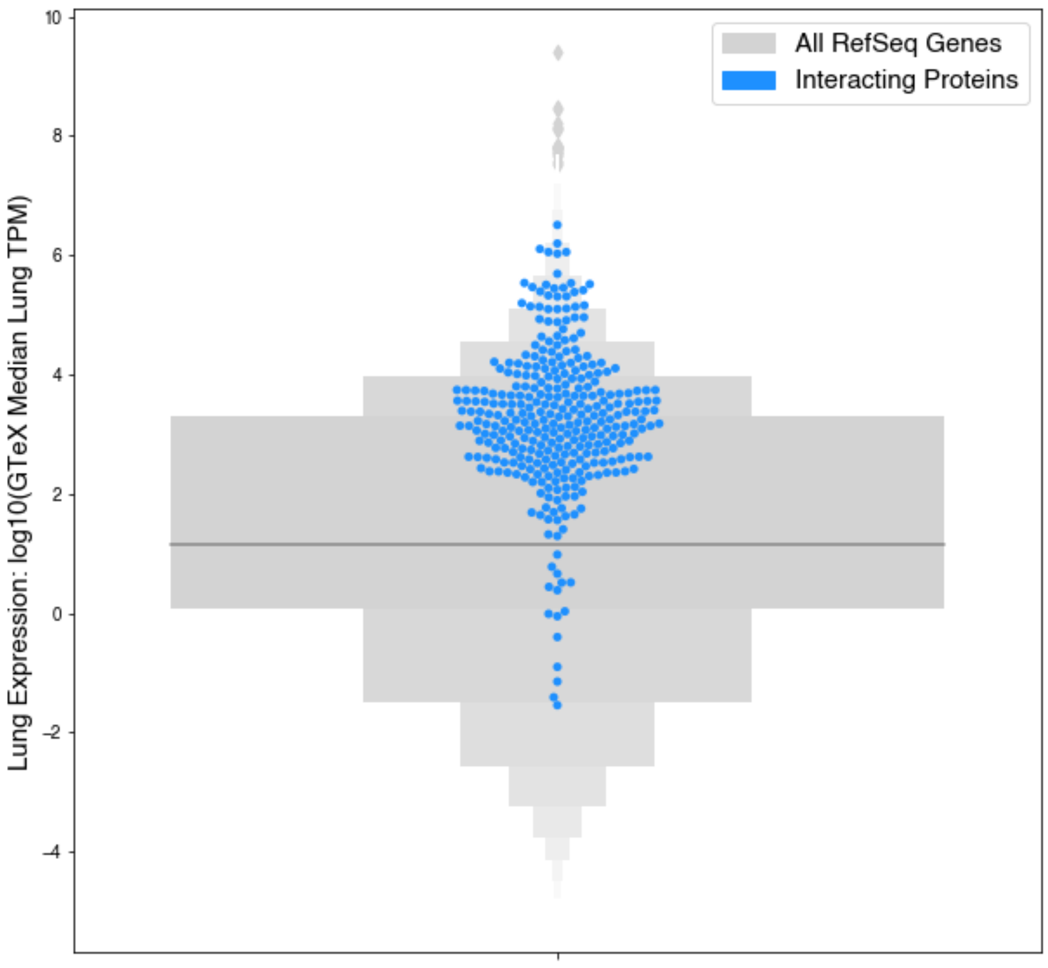


**Extended Data Figure 4. Lung mRNA expression of the interacting human proteins relative to other proteins.** The gene expression in the lung of the interacting proteins was observed to be higher that all other proteins (blue interacting proteins; median=25.52 TPM, grey all other proteins; median=3.198 TPM), p=0.0007).

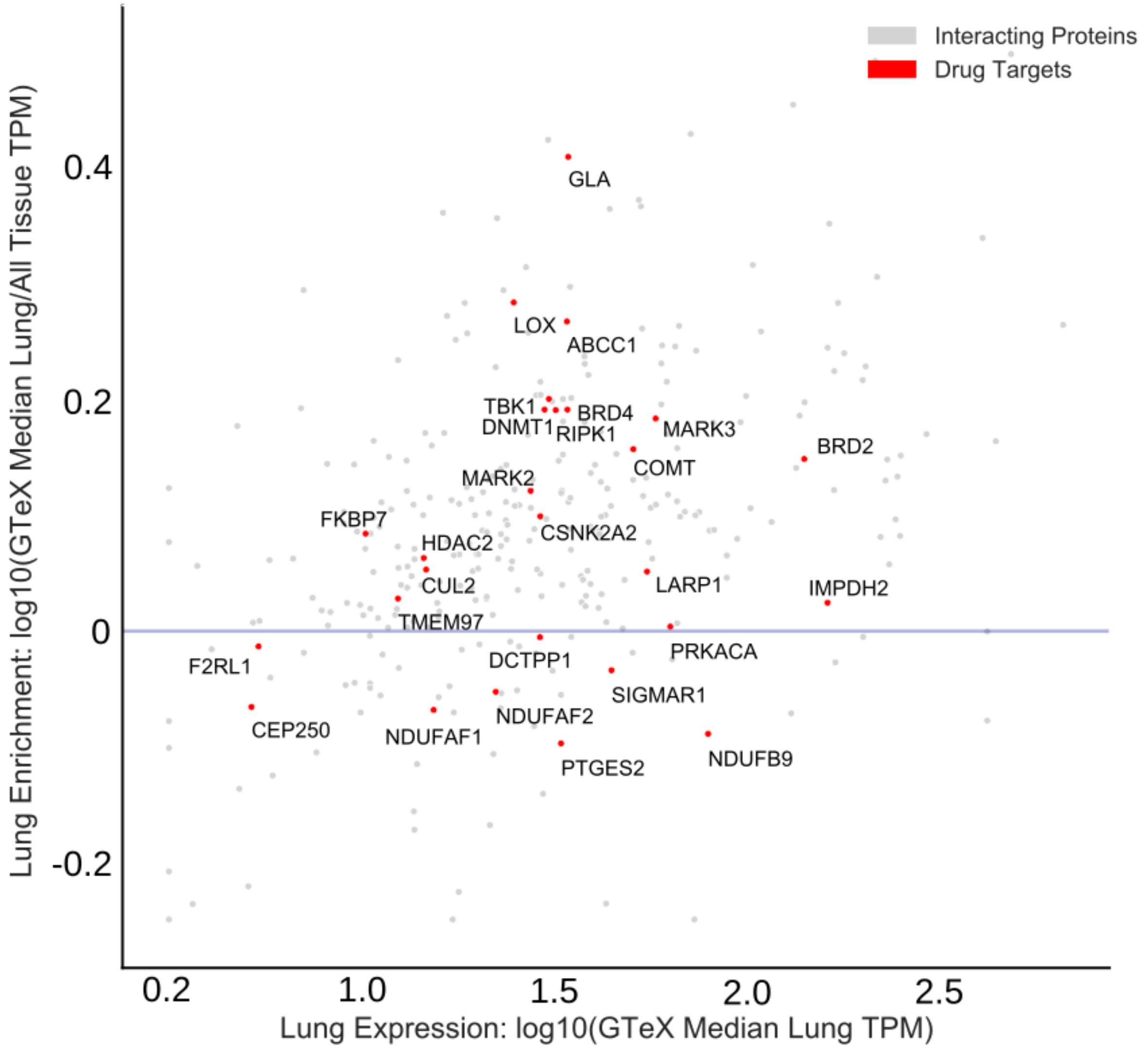


**Extended Data Figure 5. Gene level expression values and specificity for the lung.** Scatterplot of the lung mRNA expression (TPM) versus enrichment of lung mRNA expression (lung TPM/median all tissue TPM) for human interacting proteins. Red points denote drug targets that are labelled with their gene names. Points above the horizontal blue line represent interacting proteins that are enriched in lung expression and show how most human interacting proteins tend to be enriched in the lung.

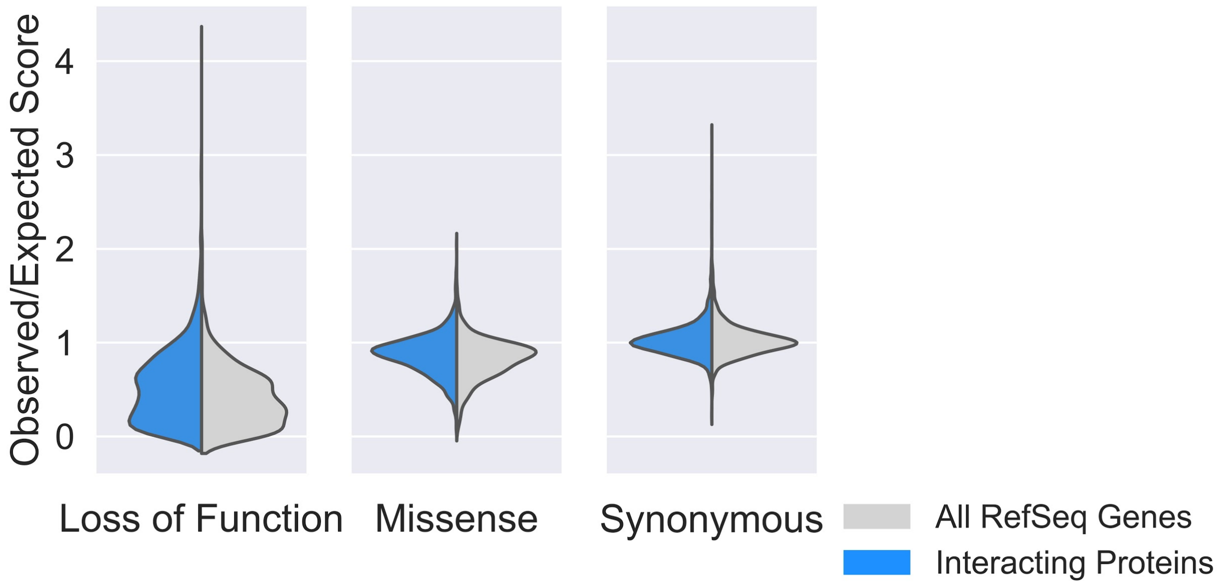


**Extended Data Figure 6. Evolutionary Conservation of SARS-CoV-2 Interacting Proteins.** We calculated the observed/expected ratio from gnomAD[1] for loss of function, missense, and synonymous mutations across all RefSeq genes. Compared to other genes (RefSeq Genes, grey), interacting proteins (blue) had lower ratios (indicating stronger intolerance of mutations) of observed/expected missense (median refseq score: 0.49, median interacting score: 0.36, p=2.46*10-11; t-test), and loss of function (median refseq score: 0.89, median interacting score: 0.85 p=9.12*10-7; t-test) but no difference in observed/expected synonymous mutations (median refseq score: 1.009 median interacting score: 1.004, p=0.48; t-test). Collectively, these results indicate that the interacting proteins have reduced genetic variation in human populations.

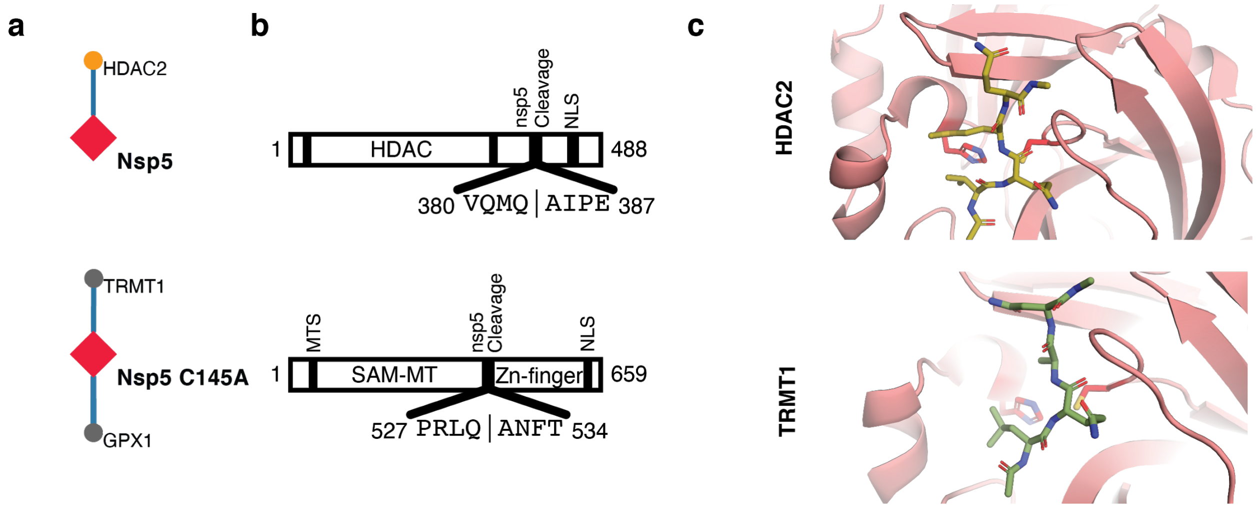


**Extended Data Figure 7. Candidate targets for the viral Nsp5 protease.** (**a**) nsp5 WT and nsp5 C145A (catalytic dead mutant) interactome. (**b**) Domain maps of HDAC2 and TRMT1 illustrating predicted cleavage sites (using Coronanet 1.0). HDAC: Histone Deacetylase Domain, NLS: Nuclear Localization Sequence, MTS: Mitochondrial Targeting Sequence, SAM-MT: S-adenosylmethionine-Dependent Methyltransferase Domain. (**c**) Peptide docking of predicted cleavage peptides into the crystal structure of SARS-CoV nsp5.

**Consensus analysis of SARS-CoV-2 Orf6 homologs**

```
Reference sequence (1): ref|YP_009724394.1
Identities normalised by aligned length.
Colored by: identity

                            cov    pid   1 [         .         .         .         .         :         .] 61
 1 ref|YP_009724394.1 100.0% 100.0%      MFHLVDFQVTIAEILLIIMRTFKVSIWNLDYIINLIIKNLSKSLTENKYSQLDEEQPMEID
 2 gb|QIG55989.1      100.0%  98.4%      MFHLVDFQVTIAEILLIIMRTFKVSIWNLDYIINLIIKNLSKSLTVNKYSQLDEEQPMEID
 3 gb|AVP78035.1      100.0%  93.4%      MFHLVDFQVTIAEILLIIMRTFKVSIWNLDYIINLIIKNLSKPPTENNCSQLDEEQPMEID
 4 gb|AGZ48837.1      100.0%  73.8%      MFHLVDFQVTIAEILIIIMRTFRIAIWNLDMIISSIVRQLFKPLTKNKYSELDDEEPMEID
 5 gb|ALK02462.1      100.0%  72.1%      MFHLVNFQVTIAEILIIIMRTFRIAIWNLDMIISSIVRQLFKPLTKNKYSELDDEEPMEID
 6 gb|ACU31045.1      100.0%  70.5%      MFHLVDFQVTIAEILIIIMRTFRIAIWNLDVIISSIVRQLFKPLTKKKYSELDDEEPMELD
 7 gb|AGZ48811.1      100.0%  70.5%      MFHLVDFQVTIAEILIIIMRTFRIAIWNLDMIISSIVRQLFKPLTKKKYSELDDEEPMELD
 8 gb|ABD75318.1      100.0%  68.9%      MFHLVDFQVTIAEILIIIMRTFRIAIWNLDVLISSIVRQLFKPLTKKKYPQLDDEEPMELD
 9 gb|AS066813.1      100.0%  67.2%      MFHLVDFQVTIAEMLIIIMRTFRIAIWNLDVLISSIVRQLFKPLTKKKYPQLDDEEPMELD
10 sp|Q0Q471.1        100.0%  68.9%      MFHLVDFQVTIAEILIIIMKTFRVAIWNLDILISSIVRQLFKPLTKKKYSELDDEEPMELD
11 gb|AHX37562.1      100.0%  70.5%      MFHLVDFQVTIAEILIIIMRTFKIAIWNLDVIISSIVRQLFKPLTKKNYSELDDEEPMEIE
12 ref|NP_828856.1    100.0%  68.9%      MFHLVDFQVTIAEILIIIMRTFRIAIWNLDVIISSIVRQLFKPLTKKNYSELDDEEPMELD
13 gb|AT098150.1      100.0%  68.9%      MFHLVDFQVTIAEILIIIMRTFRIAIWNLDMIISSIVRQLFKPLTKKNYSELDDEEPMELD
14 gb|AEA11018.1      100.0%  68.9%      MFHLVDFQVTIAEILIIIMRTFRIAIWNLDVIISSIVRQLLKPLTKKNYSELDDEEPMELD
15 gb|AT098137.1      100.0%  68.9%      MFHLVDFQVTIAEILIIIMRTFRITIWNLDMIISSIVRQLFKPLTKKNYSELDDEEPMELD
16 gb|AT098186.1      100.0%  68.9%      MFHLVDFQVTIAEILIIIMRTFRIAIWNLDIIISSIVRQLFKPLTKKNYSELDDEEPMELD
17 gb|AAS01069.1      100.0%  67.2%      MFHLVDFQVTVAEILIIIMRTFRIAIWNLDVIISSIVRQLFKPLTKKNYSELDDEEPMELD
18 gb|ACZ71831.1      100.0%  67.2%      MFHLVDFQVTIAEILIIIMRTFRIAIWNLDVVISSIVRQLFKPLTKKNYSELDDEEPMELD
19 gb|AAP72979.1      100.0%  67.2%      MFHLVDFQVTIAEILIIIMRTFRIAIWNLDVIISSIVRQLFKPLTKKNYSELDDEEPMELB
20 gb|AR076386.1      100.0%  67.2%      MFHLVDFQVTIAEILIIIMRTFRIAIWNLDVLISSIVRQLFKPLTKKNYSELDDEEPMELD
21 gb|AKZ19091.1      100.0%  70.5%      MFHLVDFQVTIAEILVIIMRTFRIAIWNLDMITSSIVTQLFKPLTKKKYSELDDEVPMEID
22 gb|AT098162.1      100.0%  67.2%      MFHLVDFQVTIAEILIIIMRTFRIAIWNLDMIISSIVRQLFKPLTKKNYPELDDEEPMELD
23 gb|ACZ72113.1      100.0%  67.2%      MFHLVDFQVTIAEILIIIMRTFRIAIWNLDVIISSIVRQLFKPLTKKNYSELDDEEPMKLD
24 sp|Q3LZX8.1        100.0%  67.2%      MFHLVDFQVTIAEILIIIMKTFRVAIWNLDILISSIVRQLFKPLTKKNYSELDDEEPMELD
25 gb|AKZ19080.1      100.0%  70.5%      MFHLVDFQVTIAEILVIIMRTFRIAIWNLDMITSSIVTQLFKPLTKKKYSELDDEVPMEID
26 gb|AGT21083.1      100.0%  67.2%      MFHLVDFQVTIAEISIIIMRTFRIAIWNLDVIISSIVRQLLKPLTKKNYSELDDEEPMELD
27 gb|ANA96032.1      100.0%  65.6%      MFHLVDFQVTIAEMLIIIMRTFRIAILNLDVLISSIVRQLFKPLTKKKYPQLDDEEPMELD
28 gb|AAP30035.1      100.0%  67.2%      MFHLVDFQVTIAEILIIIMRTFRIAIWNLDVIISSIVRQLFKPLTKKNYSELDDEELMELD
29 gb|AIA62282.1      100.0%  62.3%      MFHLVDFQVTIAEMLIIIMRTFRIAILDLDVLISSIVRQSFKPLTKKKYPQLDDEEPMELD
30 gb|AIA62334.1       82.0%  68.0%      MFHPVDFQVTIAEILIIIMRTFRIAIWNLDVIISSIVRQLFKPLTKKNYS-----------
31 gb|AP040583.1      100.0%  50.8%      MFSLVEFQVTIAELLIIIMRSLGIGLVQFQIRMIALLKIISKHLDRNQHSKLDEEVPMEID
32 dbj|BAC81367.1      49.2%  86.7%      MFHLVDFQVTIAEILIIIMRTFRIAIWNLD-------------------------------
33 ref|YP_003858588.1  98.4%  47.5%      MFSLVAFQVTVAELLILIMKSFGLALTHIQIGIVSLLKILTNRL-DRRYSKLDEEEPMEID
34 gb|AAS44598.1       65.6%  55.0%      ---------------------FRIAIWNLDVIISSIVRQLFKPLTKKNYSELDDEEPMELD
35 gb|AAS44653.1       65.6%  55.0%      ---------------------FRIAIWNLDVIISSIVRQLFKPLTKKNYSKLDDEEPMELD
   consensus/100%                        .....................htlslhphp...............................
   consensus/90%                         MFpLVsFQVTlAEhLlIIM+oF+luIhNLDhlhs.Il+pL.KsLTcppYspLD-E.PMEl-
   consensus/80%                         MFHLVDFQVTIAEILIIIMRTFRIAIWNLDhlISSIV+QLhKPLTK+pYSpLDDEpPMElD
   consensus/70%                         MFHLVDFQVTIAEILIIIMRTFRIAIWNLDhlISSIVRQLFKPLTKKpYScLDDEEPMElD
```

**Extended Data Figure 8. Consensus analysis of SARS-CoV-2 Orf6 homologs.** Multiple sequence alignment of 34 homologs of SARS-CoV-2 Orf6. Sequence coverage (*cov)* and percent identity (*pid*) shown for each homologous sequence. Amino acids are colored according to identity with SARS-CoV-2 Orf6 (sequence *1 ref|YP_009724394.1*). Colors indicate chemical properties of amino acids: small and/or hydrophobic (bright green, A, V, L, I, M, P, G), bulky (dark green, F, Y, W, H), basic (red, R, K), acidic (blue, D, E), neutral (purple, Q, N), polar (light blue, S, T), cysteine (C, yellow). The key methionine M58, and the acidic residues E55, D59, D61 of the putative NUP98-RAE1 binding motif are shown to be highly conserved. Multiple sequence alignment was visualized using the MView web server[2]: https://www.ebi.ac.uk/Tools/msa/mview/.

**Experimental and predicted structural features of BRD2 and Protein E**

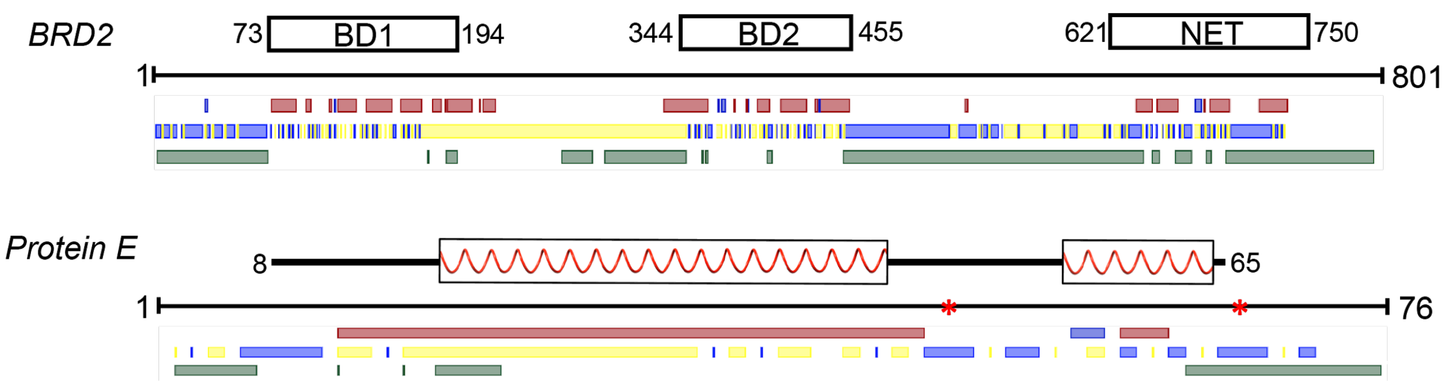


**Extended Data Figure 9. Experimental and predicted structural features of BRD2 and Protein E.** Domain names are indicated for BRD2 along with residue numbers. Areas of protein E where its SARS-CoV homolog has been described by high-resolution structural models (PDB ID: 2MM4) are shown in white blocks and thick black lines above the sequence line. The ribbons indicate alpha helical sections. Protein E is predicted to have two N-linked glycosylation sites[3] (red stars on sequence line). Structural predictions, as generated using PredictProtein[4] are found below the sequence line. The first row indicates secondary structure predictions for alpha helices (red) and beta sheets (blue). The second row beneath the sequence line indicates areas expected to be buried (yellow) and exposed (blue). At the bottom, green bars indicate areas predicted to be disordered.

**Extended Data References**


1. Karczewski, K. J. *et al.* Variation across 141,456 human exomes and genomes reveals the spectrum of loss-of-function intolerance across human protein-coding genes. *bioRxiv* 531210 (2019) doi:10.1101/531210.
2. Brown, N. P., Leroy, C. & Sander, C. MView: a web-compatible database search or multiple alignment viewer. *Bioinformatics* **14**, 380–381 (1998).
3. Fung, T. S. & Liu, D. X. Post-translational modifications of coronavirus proteins: roles and function. *Future Virol.* **13**, 405–430 (2018).
4. Yachdav, G. *et al.* PredictProtein--an open resource for online prediction of protein structural and functional features. *Nucleic Acids Res.* **42**, W337–43 (2014).
