## Supplementary Information Discussion for "A SARS-CoV-2-Human Protein-Protein Interaction Map Reveals Drug Targets and Potential Drug-Repurposing"

#### Supplementary Discussion

All SARS-CoV-2 protein and gene functions described in the subnetwork appendices, including the text below and the text found in the individual bait subnetworks, are based on the functions of homologous genes from other coronavirus species. These are mainly from SARS-CoV and MERS-CoV, but when available and applicable other related viruses were used to provide insight into function. The SARS-CoV-2 proteins and genes listed here were designed and researched based on the gene alignments provided by Chan et. al. 2020[1]. Though we are reasonably sure the genes here are well annotated, we want to note that not every protein has been verified to be expressed or functional during SARS-CoV-2 infections, either *in vitro* or *in vivo*. In an effort to be as comprehensive and transparent as possible, we are reporting the sub-networks of these functionally unverified proteins along with the other SARS-CoV-2 proteins. In such cases, we have made notes within the text below, and on the corresponding subnetwork figures, and would advise that more caution be taken when examining these proteins and their molecular interactions. Due to practical limits in our sample preparation and data collection process, we were unable to generate data for proteins corresponding to Nsp3, Orf7b, and Nsp16. Therefore these three genes have been left out of the following literature review of the SARS-CoV-2 proteins and the protein-protein interactions (PPIs) identified in this study. Below we provide what we hope is a thorough, well sourced, review of the principal interactions for each bait. Given the urgent and unprecedented nature of the current crisis and global pandemic, we hope that this will be of significant use to both the scientific and global communities.

### STRUCTURAL PROTEINS (S, E, M, N)

## S

**Function*:** Spike (S) surface glycoprotein is responsible for binding and fusion with the host membrane.

- Classified as a class I fusion protein.

- Has 2 subunits that need to be processed by cellular protease TMPRSS2. S1 mediates receptor (ACE2) binding whereas S2 mediates fusion.

Similarity to SARS

| | |
|---|---|
| **Identity:** 76.3% | **Similarity:** 87.0% |

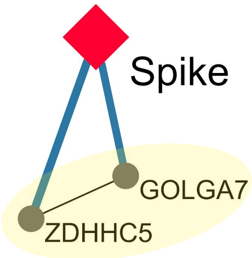


**Protein Palmitoylation:** GOLGA7-ZDHHC5 is a protein acyl-transferase (PAT) complex that may play a role in Spike palmitoylation.

## E

**Function*:** Envelope (E) protein plays a central role in virus morphogenesis and assembly.

- Acts as a viroporin, assembling in host membranes and forming pentameric protein-lipid pores that allow ion transport.

- Binds to protein M. Co-expression of M and E is sufficient for VLP formation and release. Lack of E reduces viral titers about 20-fold.

Similarity to SARS

| | |
|---|---|
| **Identity:** 94.7% | **Similarity:** 96.1% |

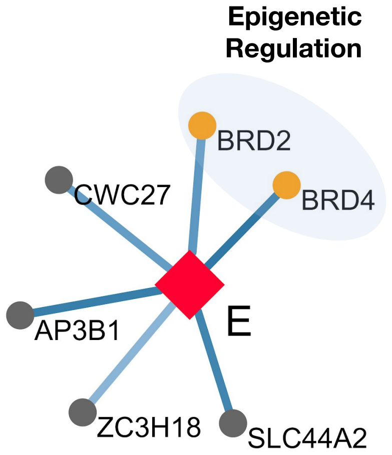


**BRD2/BRD4**: Bromodomain extra terminal (BET) proteins are implicated as epigenetic factors that regulate genes crucial for cell cycle progression, inflammation and immune response.

## M

**Function*:** Membrane (M) protein is the major driver for virus assembly and budding.

- Exists as a dimer in two major conformations, long and compact, which together determine the membrane curvature and spike density.

- M-M, M-S and M-N protein interactions contribute to virus assembly.

Similarity to SARS

| | |
|---|---|
| **Identity:** 90.5% | **Similarity:** 96.4% |

**REEP6:** Involved in IL-8 secretion, which plays a role in the recruitment of neutrophils to the lungs in patients with sepsis-induced acute respiratory distress syndrome.

**PITRM1:** Differentially expressed in RSV-infected human small airway epithelial cells.

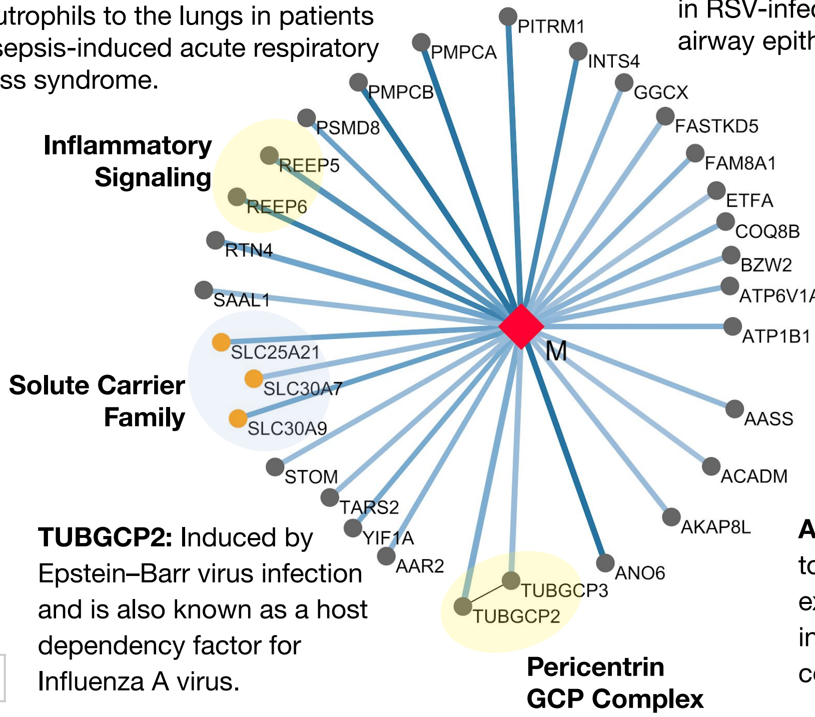


**BZW2:** A known host restriction factor for Dengue virus infection.

**ATP6V1A:** Affects Dengue, West Nile and Influenza A H1N1 virus infections.

**TUBGCP2:** Induced by Epstein–Barr virus infection and is also known as a host dependency factor for Influenza A virus.

**ANO6:** Deficiency leads to severe T cell exhaustion and the inability of the host to control viral burden.

1.0 0.7 MIST Score Spectral Counts Bait Prey Drug Target *Function based on SARS

# N

**Function*:** Nucleocapsid (N) protein binds to the RNA genome.

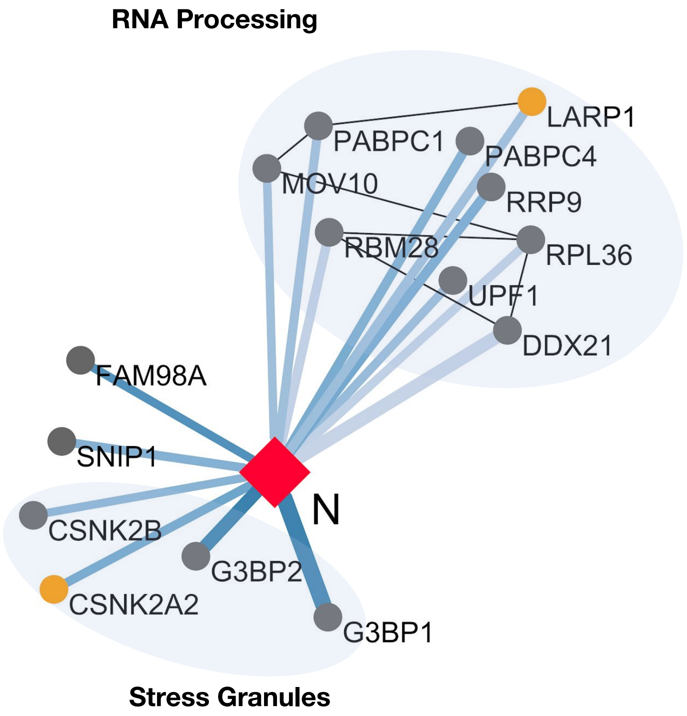


**G3BP1 and G3BP2:** G3BP1 and G3BP2 are core structural components of stress granules (SGs) which are broadly refractory to replication of viruses. Viruses have evolved diverse mechanisms such as direct cleavage of G3BPs (poliovirus, FMDV) or sequestration away from other granule components (SmFV, VV, TMCV, SAFV-2, Mengovirus, DENV, JEV, Ebola virus) to prevent granule formation. Coronaviridae like MERS and IBV also possess specific mechanisms to abrogate SG assembly. Certain viruses utilize G3BPs to promote their replication cycle, a function that is almost exclusively extra-granular.

**CK2 (CSNK2A2 and CSNK2B):** CSNK2A2 and CSNK2B are subunits of the tetrameric Casein Kinase 2. CK2 phosphorylates G3BPs and disassembles and/or inhibits the formation of stress granules. The activity of CK2 is thus presumptively proviral. CK2 is inhibited at sub-nanomolar concentrations by an orally bioavailable molecule, Silmitasertib.

**LARP1:** LARP1 is a major effector of the mTOR pathway, suppressing translation of terminal oligopyrimidine mRNAs. LARP1 binds the N protein in a variety of viruses (e.g. IBV, IAV). LARP1 knockdown decreases DENV viral titers, while inhibition of mTOR (e.g. with rapamycin) impairs MERS-CoV replication and exerts immunosuppressant functions.

**MOV10:** MOV10 is a 5' to 3' RNA helicase that interacts with UPF1 and binds 3'UTRs. Its antiviral functions are independent of the helicase activity, and often through IFN stimulation. MOV10 exhibits P-body dependent antiviral activity by binding the N protein and preventing its nuclear localization.

**PABPC1/4:** Poly-A binding proteins are involved in many steps of mRNA processing. Viral factors cause PABPC1/4 to shuttle into the nucleus, causing mRNA hyperadenylation and nuclear retention. In a complex with LARP1 and RyDEN, PABPC1/4 can promote virulence.

**UPF1:** UPF1 is a RNA helicase that functions in the nonsense-mediated mRNA decay (NMD) pathway. Murine Hepatitis Virus (MHV) mRNAs are subjected to NMD, while the MHV N protein has been shown to have NMD inhibitory functions.

Similarity to SARS

| | |
|---|---|
| **Identity:** 90.5% | **Similarity:** 94.3% |

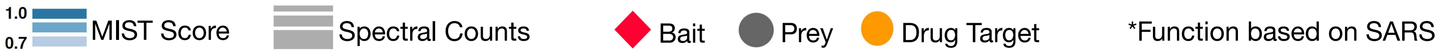

### NON-STRUCTURAL PROTEINS

#### Nsp1

**Function*:** Nsp1 antagonizes interferon induction to suppress host antiviral response.

- Overexpression of Nsp1 in A549 cells increases production of pro-inflammatory chemokines CCL5, CXCL10, and CCL3.

- Can also inhibit host gene expression by binding to ribosomes and modifying host mRNAs.

Similarity to SARS

| | |
|---|---|
| **Identity:** 84.4% | **Similarity:** 91.1% |

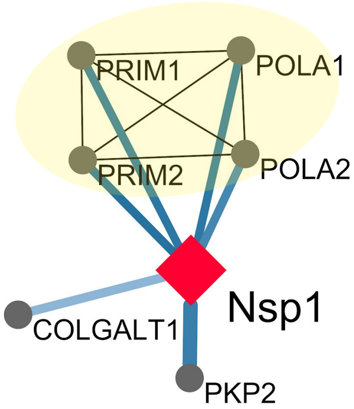


**DNA Polymerase Alpha Complex:** Regulates the activation of type I interferons through cytosolic RNA-DNA synthesis and primes DNA replication in the nucleus.

**PKP2 (Plakophilin):** Binds cadherins and intermediate fibers, crucial for desmosome formation.

**COLGALT1 (Collagen Beta (1-O) Galactosyltrasferase 1):** Required for galactosylation of collagen IV and VI to form the collagen triple helix.

#### Nsp2

**Function*:** Nsp2 is translated as part of a single protein along with Nsp3 and may serve as an adaptor for Nsp3.

-While not essential for viral replication, deletion of Nsp2 diminishes viral growth and RNA synthesis

Similarity to SARS

| | |
|---|---|
| **Identity:** 68.3% | **Similarity:** 82.9% |

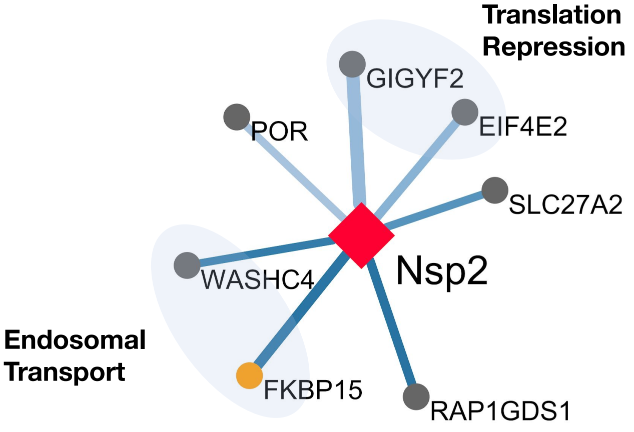


**SLC27A2**: Previously described as a Dengue virus restriction factor.

#### Nsp4

**Function*:** Nsp4 forms a complex with Nsp3 and Nsp6.

- Together, these proteins are predicted to nucleate and anchor viral replication complexes on double-membrane vesicles in the cytoplasm.

Similarity to SARS

| | |
|---|---|
| **Identity:** 80.0% | **Similarity:** 90.8% |

**TIM Complex:** Involved in the import and insertion of hydrophobic membrane proteins into the mitochondrial inner membrane. Regulates import of transmembrane proteins into the inner mitochondrial membrane.

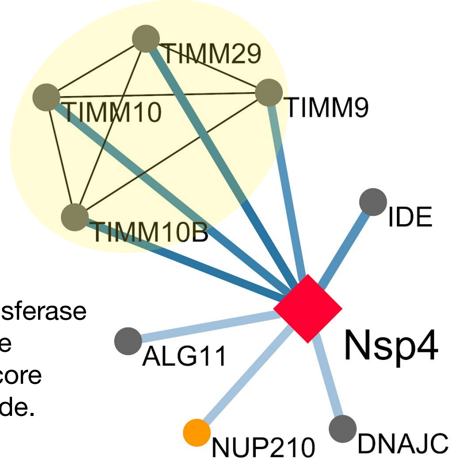


**IDE**: Involved in antigen presentation via MHC class I.

**ALG11**: Mannosyltransferase involved in the synthesis of core oligosaccharide.

1.0 / 0.7 MIST Score | Spectral Counts | Bait | Prey | Drug Target | *Function based on SARS

### Nsp5

**Function*:** Nsp5 is the 3C-like protease.

- Cleaves the viral polyprotein.

Similarity to SARS

| **Identity:** 96.1% | **Similarity:** 98.7% |
|---|---|

**HDAC2:** Deacetylates lysines at the N-terminus of histones. Reduced levels of HDAC2 leads to increased transcription of inflammatory genes. Low HDAC2 expression contributes to disease severity in COPD patients.

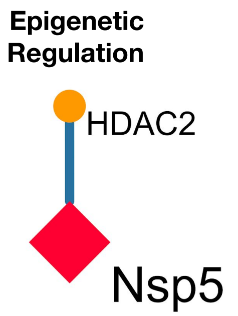


### Nsp5_C145A

**Function*:** Nsp5_C145A is a catalytically dead mutant of the Nsp5 3C-like protease.

- The catalytic residues of SARS-CoV Nsp5 align to H41 and C145 of SARS-CoV-2 Nsp5.

Similarity to SARS

| **Identity:** 96.1% | **Similarity:** 98.7% |
|---|---|

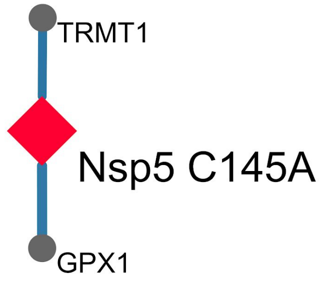


### Nsp6

**Function*:** Nsp6 limits autophagosome expansion.

- Nsp6 may favor SARS-CoV infection by compromising the ability of autophagosomes to deliver viral components to lysosomes for degradation.

- Complexes with Nsp3 and Nsp4 to form double-membrane vesicles that anchor viral replication complexes.

Similarity to SARS

| **Identity:** 87.2% | **Similarity:** 94.8% |
|---|---|

**ATP Synthase in the Mitochondria:** Components of the Mitochondrial Complex V co-purify with Nsp6. This complex regenerates ATP from ADP.

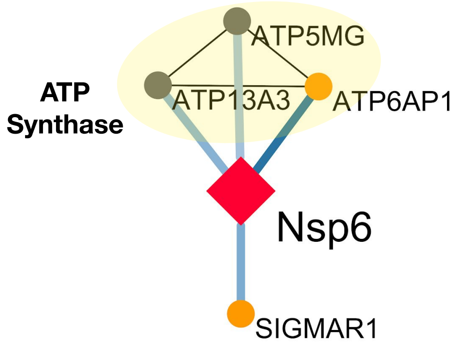


**ATP6AP1:** Subunit of the vacuolar ATP synthase protein pump. Dysregulation of ATP6AP1 results in impaired vesicle acidification and intracytoplasmic granules, resulting in a range of pathologies including an immunodeficiency syndrome and granular cell tumors. Identified as a potential host factor for IAV, WNV, and DENV. Is similar to the known IAV host factor ATP6V1A.

1.0 0.7 MIST Score    Spectral Counts    Bait    Prey    Drug Target    *Function based on SARS

### Nsp11

**Function*:** It is unclear if Nsp11 encodes a functional viral protein.

- Short peptide (13 amino acids) at the end of Orf1a.

Similarity to SARS

| | |
|---|---|
| **Identity:** 84.6% | **Similarity:** 92.3% |

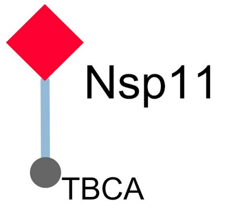


**TBCA:** Involved in the early step of the tubulin folding pathway.

1.0 0.7 MIST Score | Spectral Counts | Bait | Prey | Drug Target | *Function based on SARS

### REPLICATION COMPLEX (Nsp7, Nsp8 and Nsp12)

#### Nsp7

**Function*:** The Nsp7-Nsp8 complex is part of a unique multimeric RNA-dependent RNA replicase capable of both *de novo* initiation and primer extension.

- Forms the primase in complex with Nsp8.

**ACSL3:** Induction results in increased acyl-CoA synthesis that is essential for providing prostaglandin.

**MOGS:** Cleaves the distal alpha 1,2-linked glucose residue from the N-linked oligosaccharide precursor in a highly specific manner.

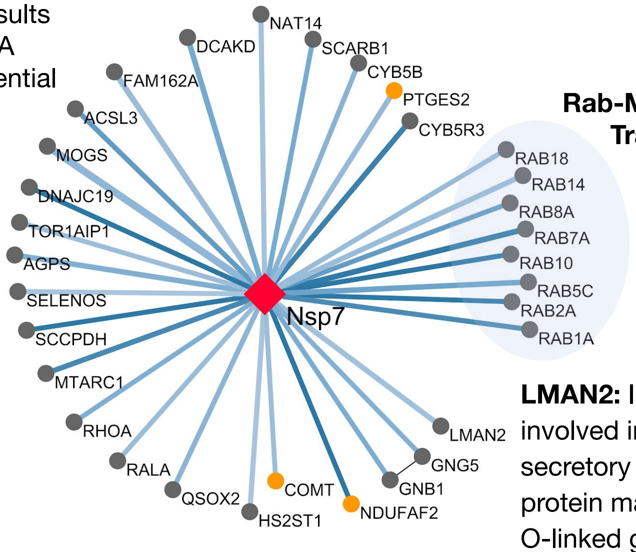


**Rab-Mediated Trafficking**

**LMAN2:** Intracellular lectin involved in the early secretory pathway. The protein may interact with O-linked glycans and N-acetyl-D-galactosamine.

Similarity to SARS

| | |
|---|---|
| **Identity:** 98.8% | **Similarity:** 100.0% |

#### Nsp8

**Function*:** Nsp8 forms a primase in complex with Nsp7.

-Eight Nsp8 proteins complex with eight Nsp7 proteins to form a hexadecameric structure surrounding dsRNA.

**Exosome Complex:** Degrades ssRNA in a 3' to 5' direction and is involved in homeostatic degradation of host RNA as well as antiviral immunity.

**7SK snRNP:** Regulates activity of the positive Transcription Elongation Factor b (p-TEFb).

**Mitochondrial Ribosome**

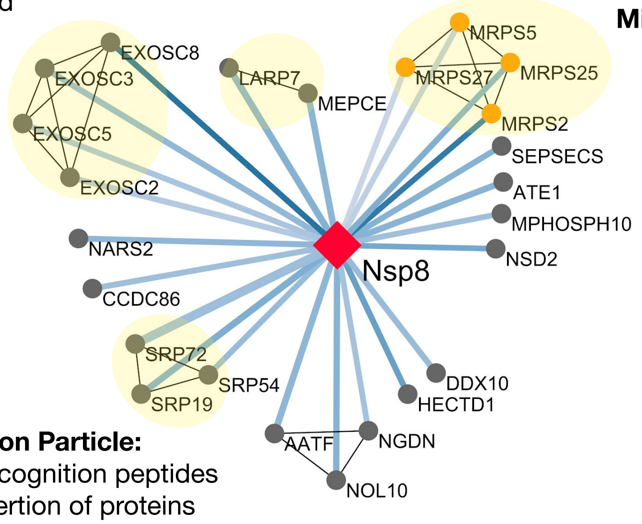


**Signal Recognition Particle:** Binds to signal recognition peptides and mediates insertion of proteins into the endoplasmic reticulum.

Similarity to SARS

| | |
|---|---|
| **Identity:** 97.5% | **Similarity:** 99.0% |

#### Nsp12

**Function*:** Nsp12 is the RNA-dependent RNA polymerase (RdRp).

- Nsp12 contains a large two-domain N-terminus of little known function and a canonical RdRp domain in the C-terminus.

- Nsp7-Nsp8 heterodimer binds the RdRp domain of Nsp12.

**Spliceosome:** Removes introns from pre-mRNA. SLU7 is essential for the second catalytic step of pre-mRNA splicing.

**RIPK1:** Triggers cell death by apoptosis or necrosis as an active regulatory kinase. Regulates inflammatory signaling and inhibits cell death as a scaffold protein.

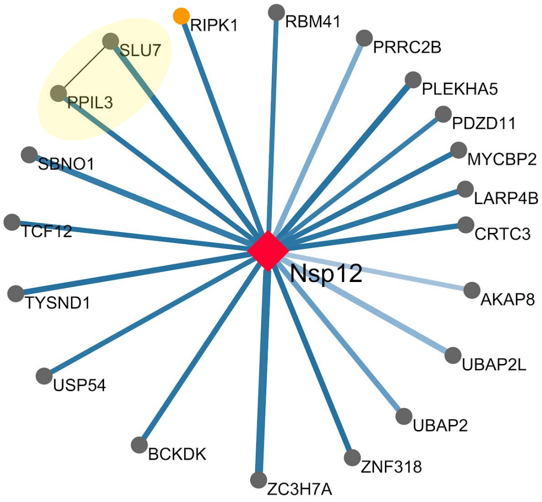


Similarity to SARS

| | |
|---|---|
| **Identity:** 96.4% | **Similarity:** 98.3% |

1.0 0.7 MIST Score — Spectral Counts — Bait — Prey — Drug Target — *Function based on SARS

### Nsp15

**Function*:** Nsp15 has uridine-specific endoribonuclease (endoU) activity and is essential for viral RNA synthesis.

- The endoUs were shown to: (i) have endonucleolytic activity; (ii) cleave 3' of pyrimidines, preferring uridine over cytidine; and (iii) release reaction products with 2'-3'-cyclic phosphate and 5'-OH ends.

- Shown to form homohexamers composed of a dimer of trimers.

Similarity to SARS

| | |
|---|---|
| **Identity:** 88.7% | **Similarity:** 95.7% |

**NUTF2:** Mediates the import of GDP-bound RAN from the cytoplasm into the nucleus, thus indirectly plays a more general role in cargo receptor-mediated nucleocytoplasmic transport.

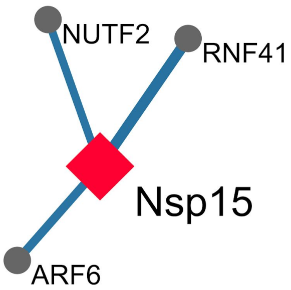


**RNF41:** Acts as a E3 ubiquitin-protein ligase. Promotes TRIF-dependent production of type I interferon and inhibits infection with vesicular stomatitis virus.

**ARF6:** GTP-binding protein involved in protein trafficking that regulates endocytic recycling and cytoskeleton remodeling as well as the activation of cholera toxin.

### Nsp9

**Function*:** Nsp9 is an essential single-stranded RNA binding protein.

- Shown to interact with the replication complex (Nsp7, Nsp8, and Nsp12).

- Binds to both DNA and RNA but preferentially binds single-stranded RNA.

Similarity to SARS

| | |
|---|---|
| **Identity:** 97.3% | **Similarity:** 98.2% |

**Nuclear Pore Complex (NPC):** Several components of the NPC are targeted by viruses in an effort to block or promote nuclear transport beneficial to virus replication.

**NEK9:** Serine/threonine kinase that regulates mitotic progression and has been shown as an adenovirus dependency factor.

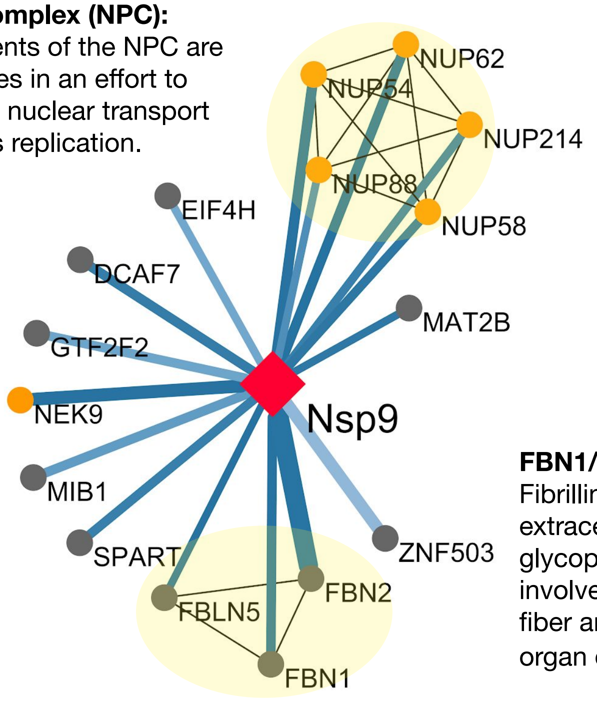


**FBN1/FBN2:** Fibrillin-1 and -2 are extracellular matrix glycoproteins involved in elastin fiber and respiratory organ development.

### Nsp10

**Function*:** Nsp10 is a zinc-finger protein essential for replication.

- Has been implicated in negative-strand RNA synthesis.

- Acts as a stimulatory factor for Nsp16 to execute its methyltransferase activity.

Similarity to SARS

| | |
|---|---|
| **Identity:** 97.1% | **Similarity:** 99.3% |

**GFER:** FAD-dependent sulfhydryl oxidase that regenerates the redox-active disulfide bonds in CHCHD4/MIA40.

**GRPEL1:** Essential component of the PAM complex, required for the translocation of transit peptide-containing proteins from the inner membrane into the mitochondrial matrix in an ATP-dependent manner.

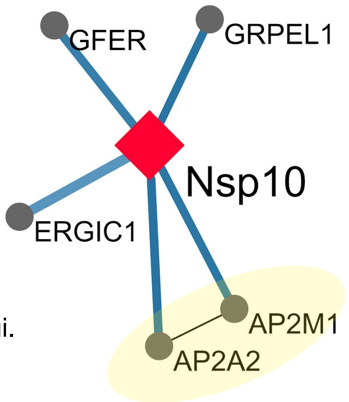


**ERGIC1:** Plays a possible role in transport between the endoplasmic reticulum and Golgi.

**AP2 Clathrin:** Nsp10 may utilize this interaction to hijack the clathrin machinery and endocytose host proteins.

1.0 0.7 MIST Score | Spectral Counts | Bait | Prey | Drug Target | *Function based on SARS

### CAPPING ENZYMES (Nsp13, Nsp14)

#### Nsp13

**Function*:** Nsp13 is a helicase and triphosphatase that initiates the first step in viral mRNA capping.

- Nsp13, along with Nsp14 and Nsp16, installs the cap structure onto viral mRNA in the cytoplasm instead of the nucleus where host mRNA is capped.

**Protein Kinase A Signaling Pathway:** PKA signalling coordinates multiple steps of membrane transport and may be hijacked for viral benefit. AKAP9 is a large scaffolding protein that localizes to the Golgi apparatus and centrosomes and binds multiple protein complexes including PKA.

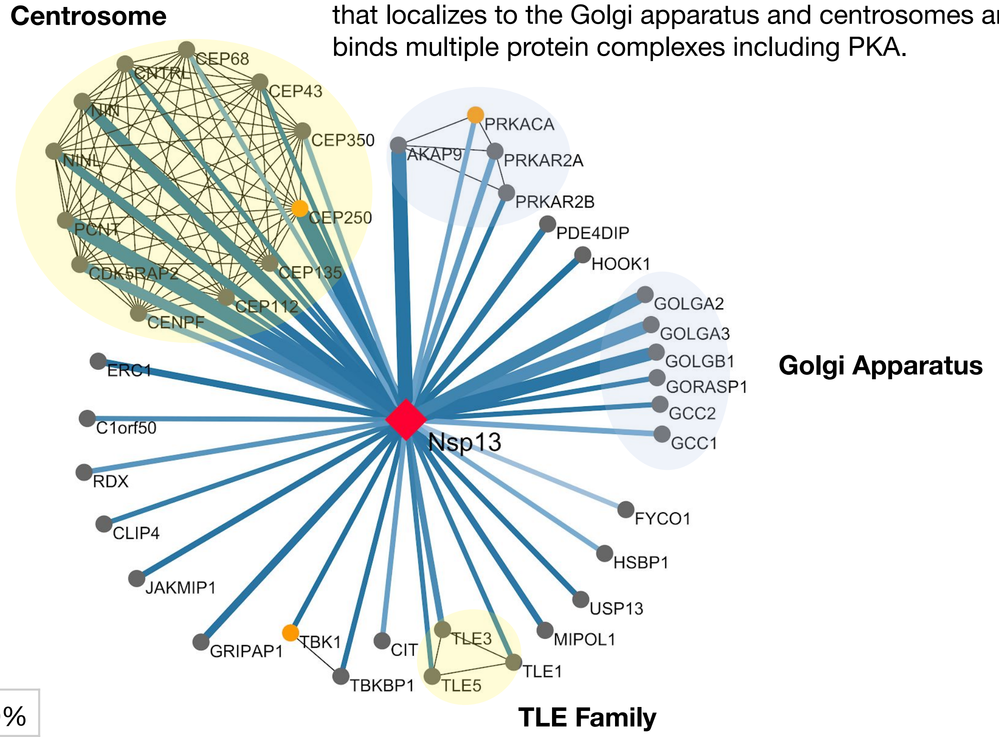


Similarity to SARS

| **Identity:** 99.8% | **Similarity:** 100.0% |
|---|---|

#### Nsp14

**Function*:** Nsp14 is a bifunctional enzyme encoding both an exonuclease domain and a separate domain that acts as a SAM dependent methyltransferase.

- The exonuclease domain corrects mutations that arise during genome replication.

- The SAM dependent methyltransferase domain facilitates capping of viral mRNA.

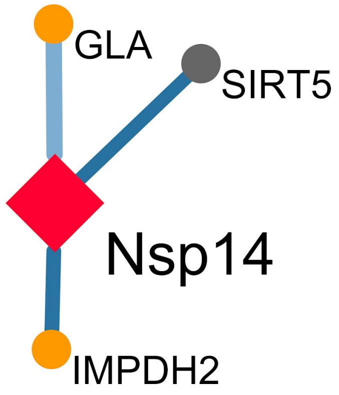


**IMPDH2:** Catalyzes the conversion of isosine 5’ phosphate (IMP) ultimately to guanine 5’ monophosphate for *de novo* synthesis of guanine nucleotides. Nsp14’s interaction with IMPDH2 may reflect an interplay with purine nucleotide metabolism.

Similarity to SARS

| **Identity:** 95.1% | **Similarity:** 98.7% |
|---|---|

1.0 0.7 MIST Score   Spectral Counts   Bait   Prey   Drug Target   *Function based on SARS

### OPEN READING FRAMES

#### Orf3a

**Function*:** Orf3a is not essential for replication but contributes to pathogenesis. It is packaged into virions.

- Mediates trafficking of Spike (S protein) by providing an ER/Golgi retention signal.

- Induces elevation of IL-1β secretion and activates NF-κB and the NLRP3 inflammasome by promoting TRAF3-dependent ubiquitination of p105 and ASC.

- Expression of Orf3a induces apoptosis in viral infection and cell line models.

Similarity to SARS

| | |
|---|---|
| **Identity:** 72.4% | **Similarity:** 85.1% |

**ALG5**: Involved in N-linked protein glycosylation. Identified as a host factor for IAV replication.

**HMOX1:** Key enzyme in heme catabolism. Has shown a cytoprotective and anti-inflammatory effect both in pulmonary pathologies and viral pathogenesis.

**CLCC1**: An intracellular chloride channel implicated in ER stress. Interacts physically with RSV and druggable protein SLC5A13.

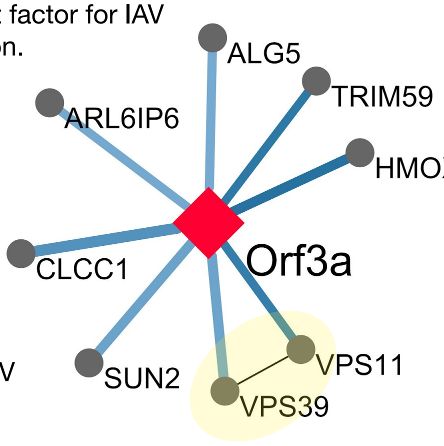


**Lysosomal Fusion Complexes:** VPS11 and VPS39 are members of the HOPS and CORVET complexes, respectively, which coordinate fusion of the lysosome with the endosome and autophagosome.

#### Orf3b

**Function*:** Orf3b is shown to be an interferon antagonist and is involved in pathogenesis.

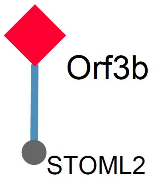


Similarity to SARS

| | |
|---|---|
| **Identity:** 7.1% | **Similarity:** 9.5% |

#### Orf6

**Function*:** Orf6 is a type 1 interferon antagonist. Expression of Orf6 suppresses the induction of interferon and interferon signaling pathways.

- C-terminal region of SARS-CoV Orf6 interacts with the nuclear import protein, karyopherin alpha-2, sequestering it in the cytoplasm. This prevents import of STAT1, activator of interferon response genes, into the nucleus.

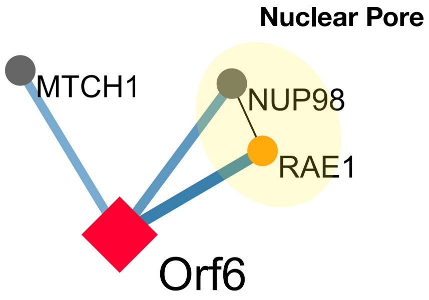


Similarity to SARS

| | |
|---|---|
| **Identity:** 66.7% | **Similarity:** 85.7% |

1.0 0.7 MIST Score   Spectral Counts   Bait   Prey   Drug Target   *Function based on SARS

### Orf7a

**Function*:** Orf7a may play a role in pathogenesis via its role in virus-induced apoptosis.

- ΔOrf7a SARS-CoV is still able to release virions at similar levels as wild-type virus.

Similarity to SARS

| | |
|---|---|
| **Identity:** 85.2% | **Similarity:** 90.2% |

**HEATR3**: Involved in ribosomal protein transport. Implicated in mediating NF-kB signaling.

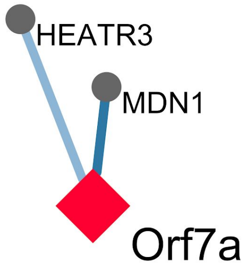


**MDN1**: Nuclear chaperone that is critical for the export of pre-60s ribosome subunits.

### Orf8

**Function***: Orf8 is an accessory protein not essential for virus replication *in vitro* and *in vivo*.

- Previously shown to be a recombination hotspot, one of the most rapidly evolving regions among SARS-CoV genomes.

- SARS-CoV Orf8b was shown to induce ER stress and activate NLRP3 inflammasomes.

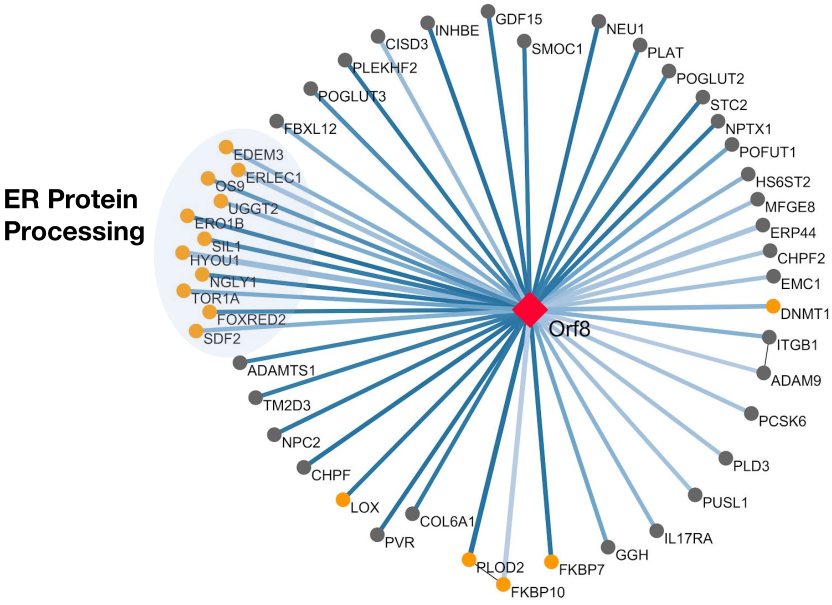


Similarity to SARS

| | |
|---|---|
| **Identity:** 28.5% | **Similarity:** 45.3% |

### Orf9b

**Function***: Orf9b is an accessory protein synthesized from an alternative complete reading frame within the viral N gene.

- Targets the mitochondrial-associated adaptor molecule MAVS signalosome by utilizing PCBP2 and E3 ligase AIP4, resulting in the degradation of MAVS and therefore limiting host cell interferon responses.

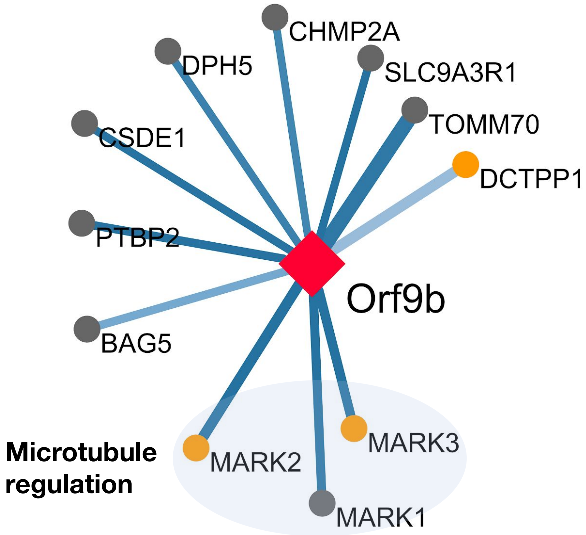


**TOMM70 (Translocase Of Outer Mitochondrial Membrane 70):** Receptor that accelerates the import of all mitochondrial precursor proteins. TOMM70 interacts with MAVS protein upon virus infection.

Similarity to SARS

| | |
|---|---|
| **Identity:** 72.4% | **Similarity:** 84.7% |

1.0 – 0.7 MIST Score | Spectral Counts | Bait | Prey | Drug Target | *Function based on SARS

### Orf9c

**Function*:** Orf9c is a short polypeptide (70 amino acids) dispensable for viral replication. There is no data yet providing evidence that the protein is expressed during SARS-CoV-2 infection.

**ABCC1:** ABCC1 (also MRP1) is a multifunctional ATP-binding cassette protein involved in controlling the efflux of drugs in cells. MRP1 is a well-known viral host factor and physical interactor of both IAV and WNV proteins. Further, it has been implicated in disease progression of pneumonia and COPD, as well as drug resistance in the lung.

**F2LR1:** The protein product of F2LR1, PAR2, is a protease-activated receptor that has a cytoprotective and inflammatory role in the pathogenesis of viral infection and progression of pulmonary disease.

**BCS1L:** BCS1L is a mitochondrial chaperone located in the inner mitochondrial membrane. Loss of BCS1L is associated with severe clinical disorders such as GRACILE and Bjornstad Syndrome.

**ALG8:** This enzyme adds the second glucose residue to the lipid-linked N-linked glycan.

**Respiratory Electron Transport:** Orf9c interacts with four proteins in the mitochondrial respiratory electron transport chain complex I, which has demonstrated roles in TLR/IL-1 signaling and mediating inflammation.

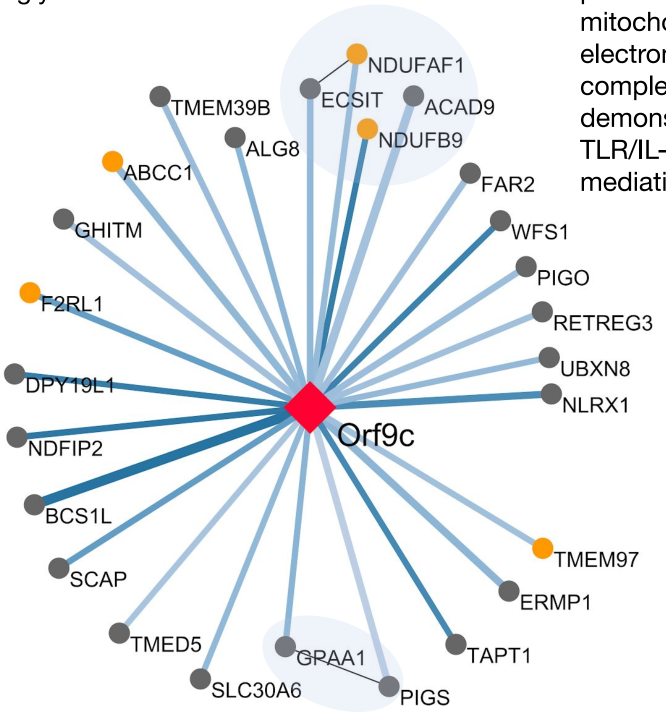


Similarity to SARS

| | |
|---|---|
| **Identity:** 74.0% | **Similarity:** 78.1% |

**GPI-Anchor Biosynthesis:** GPAA1 is essential for GPI-anchoring of precursor proteins, while PIGS and PIGO are involved in GPI-synthesis.

### Orf10

**Function*:** Orf10 codes for a peptide only 38 amino acids long. There is no data yet providing evidence that the protein is expressed during SARS-CoV-2 infection.

- Does not have a homolog in SARS-CoV.

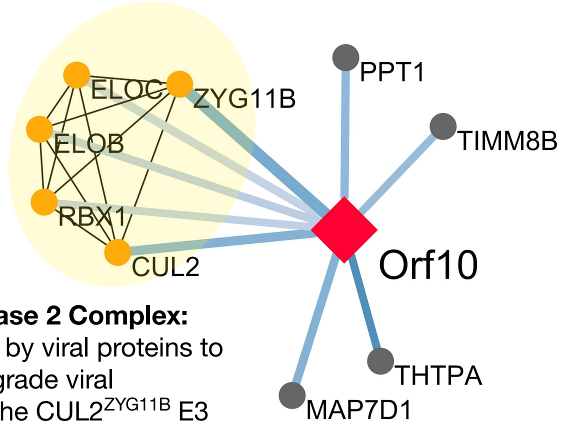


**Cullin RING E3 Ligase 2 Complex:** Commonly hijacked by viral proteins to ubiquitinate and degrade viral restriction factors. The CUL2$^{ZYG11B}$ E3 ligase targets substrates with exposed N-terminal glycines for degradation.

1.0 0.7 MIST Score   Spectral Counts   Bait   Prey   Drug Target   *Function based on SARS

### SARS-CoV Structural Proteins

#### SARS-CoV-2 Spike/S Protein

Spike (S protein) is the viral surface protein that mediates viral entry through its interaction with cellular host factor angiotensin-converting enzyme 2 (ACE2)[2]. The full-length S protein is 141 kDa (1273 amino acids (aa)) and encodes a N-terminal signal peptide, a transmembrane region (aa 1214-1237), and a patch of cysteine residues (aa 1235-1254) which are predicted to be palmitoylated (protein S-acylation of cysteine residues)[3,4]. In previous studies, mutation of these cytosolic cysteine residues inhibited fusogenicity of S, and suggest that targeting the s-acylation modifications via acyl-transferase inhibitors could potentially be a therapeutic strategy for inhibiting coronavirus infection[5].

In our interactome, we demonstrate high-confidence binding of SARS-CoV-2 Spike to the GOLGA7-ZDHHC5 complex, a protein acyl-transferase (PAT) complex. GOLGA7 characteristically localizes to the Golgi apparatus[6] but the GOLGA7-ZDHHC5 complex has been found at the plasma membrane[7]. While this interaction is exciting and could potentially point to a targetable enzymatic activity responsible for Spike palmitoylation, further experiments need to be done to fully clarify the functional role of this complex and the consequence of its inhibition. ZDHHC5 is highly expressed in neuronal tissue and is reported to have a role in hippocampal function[8]. In addition, ZDHHC5 is shown to palmitoylate G protein-coupled receptors[9] and activate nucleotide oligomerization domain (NOD)-like receptors 1 and 2 (NOD1/2)[10] which recognize pathogenic peptidoglycans and activate immune signaling[10]. GOLGA7 is also a general adaptor for ZDHHC proteins and, in complex with these acyl-transferase proteins, can regulate additional processes including the palmitoylation of Ras[11]. It should also be noted that the ZDHHC5-GOLGA7 complex is required for the non-apoptotic cell death phenotype attributed to the synthetic oxime CIL56, indicating a role for this complex in the cell death pathway[7]. Thus, we would caution against long-term inhibition of this enzyme as a general treatment strategy.

Though identified just below our cutoffs, one of the lower confidence hits for Spike was ATP1A1, a protein at the plasma membrane that promotes viral entry and replication of related coronaviruses (M-CoV, FCoV, MERS-CoV), and a number of other viruses including Ebola, Lassa, live-attenuated Junín virus, and respiratory syncytial virus (RSV)[12–15]. Interestingly, this protein is from the same ATPase family as ATP1B1, a high-confidence interactor of M protein.

#### SARS-CoV-2 E Protein

The SARS-CoV Envelope (E) protein, a small integral membrane protein, is important for virus production, assembly, intracellular trafficking, morphogenesis and virulence[16,17]. The transmembrane domain of E has amphipathic properties and can oligomerize to form cation-permeant ion channels[18]. The C-terminal cytoplasmic domain contains a targeting signal for protein localization to the Golgi complex[19,20] and a PDZ binding motif important for virulence[21]. Interestingly, E protein is expressed at very high levels inside the infected cells, but only a small portion is incorporated in virions and the rest is largely distributed in intracellular membranes between ER and Golgi compartments indicating an important role of E protein in manipulating the cellular environment[22]. Increased cellular stress, unfolded protein response, apoptosis, and heightened host immune response were all observed in infected cells lacking E protein[17]. In addition, E has also been implicated in inflammasome activation and consequential inflammation in the lung parenchyma[17].

In our study, we identify six high-confidence host protein interactions with E protein: AP31B, BRD2, BRD4, CWC27, SLC44A2, and ZC3H18. BRD2 and BRD4 are bromodomain extra terminal (BET) family proteins that bind acetylated chromatin and activate transcription[23]. BRD4 is a co-activator of NF-kB and facilitates the transcription of NF-kappaB-dependent inflammatory response[24]. Sequestration of BRD4 by E protein may represent a means for SARS-CoV-2 to protect against the host immune response. Previously it's been shown that inhibition of BRD4 activity in primary lung epithelial cells results in diminished innate immune response after challenge with poly(I:C), a viral pattern that simulates acute RNA virus infections[25]. In addition, the interaction of BRD4 with Bovine Papillomavirus E2 Protein tethers viral DNA to host mitotic chromosomes ensuring viral persistence in infected cells[26]. BRD2 is a transcriptional regulator that binds hyperacetylated histones and is implicated in a variety of cellular processes [27,28]. Though functionally similar, BRD2 and BRD4 are shown to regulate different transcriptional programs [29]. Interestingly, the α3 helix at the N-terminus of histone 2A shares local sequence similarity (~15 non-contiguous aa residues) with an α-helix of E protein (see main text **Fig. 4d**). This aligned sequence spans many acetylated lysine residues shown to bind BRD2[30]. This information suggests a potential role for E protein to act as a sort of molecular mimic, disrupting the H2A-BRD2 interaction in order to disrupt BRD2 regulated transcription programs in a manner beneficial to the virus. Given that bromodomain proteins and their ability to regulate transcription are implicated in the life cycle of several viruses[26,27,31,32], the observed interactions of E protein with BRD2 and BRD4 present exciting avenues to pursue. However, future research is still needed to fully establish the structural basis of these interactions and the resulting functional implication for SARS-CoV-2 infection.

E protein also interacts with host factors involved in protein and mRNA trafficking, important for potentially carrying out many of E's functions. AP3B1 encodes the beta-1 subunit of adaptor protein complex 3 (AP-3), that is involved in signal-mediated protein sorting from Golgi membranes to endosomal-lysosomal organelles[33,34]. Mutations in AP3B1 affect protease activity in endosomes and produce pathology-associated defects with aberrant transmembrane lysosomal protein trafficking[35]. AP-3 depletion resulted in decreased localization of the HIV-1 major structural protein Gag from plasma membrane and late endosome, suggesting that intact AP-3 is required for HIV-1 particle production and release[34]. Also, we have previously shown that AP-3 is targeted by the NS5 protein of West Nile virus[36]. SARS-CoV-2 may also utilize AP-3 for particle assembly, transport, and release.

Other notable high-confidence host protein interactions with E protein include CWC27 and SLC44A2. Both SLC44A2 and CWC27 are proteins implicated in other diseases, but have yet to be characterized in relation to viral infections.

**SARS-CoV-2 M Protein**

Membrane (M) protein, the most abundant protein in coronavirus particles, is a type III transmembrane glycoprotein located in the virus envelope. M protein manipulates cellular membranes to bring viral and host factors together and therefore is the main driver for virus assembly and budding processes. M-M, M-S and M-N protein interactions all contribute to this assembly process[37]. In the virion, M exists as a dimer in two major conformations (long and compact), which together determine the membrane curvature and spike density[38,39]. SARS-CoV M protein is known to affect various host processes in a manner beneficial to viral replication and infectivity[40]. M suppresses key inflammatory molecules (e.g. NF-κB and Cox-2)[41] and counteracts host viral defenses by inhibiting type I interferon production[42]. SARS-CoV M protein modulates apoptosis by interfering with PDK1–PKB/Akt signaling. This interaction between M protein and PDK1 could be targeted as a plausible therapeutic approach to modulate the pro-apoptotic properties attributed to coronavirus infection[43–45].

In this study, we identified 30 high-confidence physical interactors of SARS-CoV-2 M protein. Transmembrane domain-mediated oligomerization of M protein at ERGIC membranes drives the assembly of coronavirus which buds into the lumen of the ERGIC[46]. Therefore, it was not surprising to find many SARS-CoV-2 M protein interactors were transmembrane proteins including mitochondrial proteases (PITRM1, PMPCA, PMPCB), ATPases (ATP6V1A, ATP1B1) and kinases (AKAP8L, COQ8B, FASTKD5). In addition, we identify known host dependency (ATP6V1A, ATP1B1, TUBGCP2, RTN4) and restriction (BZW2) factors of other viruses that bind to SARS-CoV M protein. ATP6V1A is a dependency factor for Dengue virus[47], West Nile virus[47] and Influenza A H1N1 virus[48], ATP1B1 is involved in viral autophagy[49] and has a role in Sindbis virus (SINV) infection[50], and TUBGCP2 is a host dependency factor for Influenza A virus[51,52]. RTN4 is a reticulon protein involved in the formation and stabilization of endoplasmic reticulum (ER) tubules[53,54]. Reticulon proteins play crucial roles in viral RNA replication, compartment formation, and function[55] and are known to protect ER membrane integrity during polyomavirus SV40 infection[56]. RTN4 also is involved in the formation of the *Legionella pneumophila* containing vacuoles[57]. In a functional RNAi screen, BWZ2 was identified as a host restriction factor for DENV infection[58]. BZW2 is a translation repressor known to regulate the PI3K/AKT/mTOR signaling pathway[59–61]. Recently, BZW2 was identified as an oncogene implicated in a number of cancers including lung adenocarcinoma, where overexpression was negatively correlated with overall disease free survival[60–64]. Studies suggest that BWZ2 overexpression could also contribute to drug resistance in cancer cells, including resistance to rapamycin[61].

The mechanism behind acute respiratory distress in severe SARS-CoV-2 infections is not understood but the host immune response is likely to determine the pathogenesis to some extent. We found that two related host factors REEP5 and REEP6 copurify with SARS-CoV-2 M protein. REEP5 and REEP6 can refine CXCR1-mediated cellular responses and lung cancer progression[65]. REEP6 is also involved in IL-8 secretion[66]. IL-8 is the major chemoattractant for neutrophils and is implicated to have a major role in the recruitment of neutrophils to the lungs in patients with sepsis-induced acute respiratory distress syndrome[67]. Another high-confidence M protein interactor ANO6 is also very exciting from the perspective of viral pathogenesis. ANO6 is required to curb excessive T cell responses in chronic viral infections. ANO6-deficient T cells are hyperactivated during the early phase of infection, exhibiting increased proliferation and cytokine production. This overactivation ultimately leads to severe T cell exhaustion and the inability of the host to control viral burden[68]. Interestingly, ANO6 is localized in late endosomes and SARS-CoV is also targeted to endosomes[69]. Mice deficient for a related protein ANO1 suffer from tracheomalacia and die shortly after birth because of respiratory failure; ANO6 is highly expressed in the respiratory system and its involvement in respiratory functions is likely[70] which makes it an interesting candidate for future mechanistic studies in the context of SARS-CoV-2 infection.

**SARS-CoV-2 N Protein**

SARS-CoV nucleocapsid (N) protein is an essential structural protein that binds to viral genomic RNA (gRNA) in virions. N protein dimerization and its association with viral gRNA are crucial for viral assembly[71,72]. It has been shown that the N protein binds to both intracellular gRNA and subgenomic RNA (sgRNA), suggesting functions in viral transcription and translation[73]. N protein also has implications in a variety of cellular responses, including stress granule formation and host translation shutoff[74], inhibition of the nonsense mediated mRNA decay (NMD) pathway[75] and cell cycle regulation[76].

The SARS-CoV-2 N interactome includes 15 high-confidence host protein interactions, many of which are host mRNA binding proteins, including stress granule (SG)-related factors (G3BP1/2, MOV10, CK2 subunits, PABP), mRNA decay factors (MOV10 and UPF1), mTOR translational repressors (LARP1), and protein kinases (CK2). Several N protein interactors including G3BP1, G3BP2, LARP1, MOV10, PABPC1 and PABPC4 have also been detected in a mouse hepatitis virus interactome[77] and in two previous nucleocapsid protein interactomes, namely infectious bronchitis virus[78] and Influenza A virus[79]. Numerous viruses have evolved diverse mechanisms to abrogate SG assembly. Poliovirus specifically cleaves G3BP to facilitate translation of viral mRNAs[80]. Foot and Mouth Disease virus cleaves G3BP and inhibits SG formation[81]. Several other viruses including Semliki Forest virus, vaccinia virus, cardioviruses, dengue virus, japanese encephalitis virus, Ebola virus sequester G3BP to inhibit SG formation and to promote virus replication[82,83,84,85,86,87,88]. Amongst *Coronaviridae*, MERS virus employs its dsRNA-binding 4a protein to inhibit SG formation and promote viral replication[89] whereas IBV infection prevents the formation of SGs via yet-unknown mechanisms[90].

N protein also interacts with subunits of the broad-spectrum Casein Kinase 2 (CK2) complex (CSNK2A2 and CSNK2B). The N protein of other *Coronaviridae* has been shown to be phosphorylated at sites predicted to be substrates for CK2[91], and there is a NetPhos predicted CK2 phosphorylation site in N. CK2 has also previously been shown to inhibit granule formation or promote granule disassembly in a G3BP phosphorylation-dependent manner[92]. CK2 inhibition sequesters N protein in the nucleus, away from the G3BP subunits of SGs[76]. It is possible that the SARS-CoV-2 N protein inhibits the formation of cellular SGs, potentially by scaffolding the phosphorylation of G3BP by CK2. siRNA-mediated knockdown of either CSKN2A2 or CSNK2B reduced viral replication of SARS-CoV, indicating that CK2 may have a proviral function[93]. These findings suggest that the CK2 inhibitor Silmitasertib (also known as CX-4945) could be effective in slowing SARS-CoV-2 replication. Silmitasertib is a sub-nanomolar inhibitor of CK2 currently in phase II trials for indications such as multiple myeloma or metastatic basal cell carcinoma[94]. Potential antiviral effects of Silmitasertib via inhibition of CK2 and subsequent increase in the formation of antiviral SGs merits further investigation.

UPF1 is an RNA helicase functioning in the nonsense-mediated mRNA decay (NMD) pathway. In common with other virus species in the order *Nidovirales*, SARS-CoV-2 produces a nested set of sgRNAs sharing the same 3'UTR but are different in number of ORFs contained[95]. In principle, only the first ORF in the 5' end is translated. Thus, many of the subgenomic mRNAs have long 3' UTRs and are potentially targeted for NMD. A previous study showed that Murine Hepatitis Virus (MHV) mRNAs were subjected to NMD[75]. Transfection of plasmids containing MHV N protein had an NMD inhibitory function and prevented MHV mRNA from rapid decay. Based on the observation that SARS-CoV-2 shares a similar feature of nested sgRNA with MHV, we expect it is also subjected to NMD, and its N protein may have similar NMD inhibitory function. It is also possible that UPF1 is involved in the programmed ribosomal frameshifting during SARS-CoV-2 translation given its discovered function in suppressing nonsense mutations[96].

LARP1 is an RNA binding protein, which is known to regulate protein synthesis as well as modulate mTOR pathway[97]. LARP interacts with actively translating ribosomes via another hit from our list, PABPC1, also a regulator of mTOR pathway [98]. Activation of the mTOR pathway is advantageous for a broad spectrum of virus species, as it counteracts the host cell response, by inhibiting autophagy and apoptosis[99], therefore inhibition of mTOR activity has been proven useful to counteract viral infection and replication. For instance, Sirolimus (rapamycin) has been used to reduce MERS-CoV infection[100,101] and alleviate H1N1 pneumonia and acute respiratory failure[102], therefore mTOR pathway is a potential therapeutics target for SARS-CoV-2 as well. Another interesting aspect about PABPC1/4 is that they are known to shuttle into the nucleus on interaction with Nsp1 proteins from different viruses (including SARS-CoV), down-regulating gene expression via hyperadenylation and nuclear retention of mRNAs[103]. Also, in a complex with LARP1 and RyDEN, PABPC1/4 can promote DENV replication[104]. PABPC4 has been described as a potential biomarker for primary lung adenocarcinoma[105].

Moloney Leukemia Virus 10 Protein (MOV10) is a host cytoplasmic 5' to 3' RNA helicase that interacts with UPF1 and binds 3'UTRs[106]. It exhibits antiviral functions independent of its helicase activity towards PRRSV, DENV and Influenza A viruses, through IFN stimulation[107,108]. Additionally, MOV10 exhibits P-body dependent antiviral activity by binding the N protein and preventing its nuclear localization[109,110].

#### SARS-CoV Non-Structural Proteins

**SARS-CoV-2 Nsp1**

In SARS-CoV, Nsp1 is likely dispensable for CoV RNA synthesis[111], but may play specific roles in the interaction of the virus with the innate immune response via directly antagonizing IFN induction[112]. In addition, overexpression of Nsp1 in the lung epithelial cell line A549 increases the production of the chemokines CCL5, CXCL10 and CCL3 30-200-fold compared with mock-transfected cells or cells expressing Nsp5, suggesting that Nsp1 may contribute to the inflammatory phenotype of SARS-CoV and SARS-CoV-2 pathology[113].

In our interactome, SARS-CoV-2 Nsp1 interacts with six host proteins. Four of these host proteins form the DNA polymerase alpha complex (POLA1, POLA2, PRIM1, and PRIM2). The DNA polymerase alpha complex was recently shown to modulate the type I interferons through cytosolic RNA:DNA synthesis[114], raising a possibility that SARS-CoV-2 Nsp1 may bind to the DNA polymerase alpha complex in the cytosol and modulate its activity to antagonize the innate immune response. Alternatively but not exclusively, Nsp1 may also interfere with the canonical DNA replication function of the complex, causing DNA replication stress and ATR activation[115]. Along this line, treatment with ATR inhibitors significantly reduced viral RNA replication, but it remains elusive how ATR activation promotes viral replication. Other proteins interacting with Nsp1 are PKP2, which interacts with Influenza A virus PB1 protein and restricts Influenza A virus replication[116]; and therefore may act as a restriction factor for SARS-CoV-2.

**SARS-CoV-2 Nsp2**

Nsp2 is highly variable among *Coronaviridae*, and while not essential for viral replication, deletion of Nsp2 in SARS-CoV diminishes viral growth and RNA synthesis[117,118]. Nsp2 is translated as part of a single protein along with Nsp3 and may serve as an adaptor for Nsp3[118].

In this study, we identified seven high-confidence host protein interactions of Nsp2. Among these are endosomal proteins FKBP15 and WASHC4 that regulate endosome transport[119–121]. Because the related virus SARS-CoV is translocated to endosomes after cell entry[2], interaction with endosomal transport proteins may reflect a mechanism by which SARS-CoV-2 gains cell entry. Nsp2 also interacts with translational repressors EIF4E2 and GIGYF2, and disruption of the EIF4E2-GIGYF2 complex leads to increased translation[122]. Nsp2 may bind to EIF4E2 and GIGYF2 to modulate translation of host and viral mRNAs. Nsp2 was also shown to interact with the acyl-CoA synthetase SLC27A2, a previously described Dengue virus restriction factor[123].

SARS-CoV-2 Nsp4

Nsp4 is likely essential, as loss of the Nsp3-Nsp4 interaction eliminated viral replication using an infectious cDNA clone and replicon system of SARS-CoV[124]. Nsp4 is a non-structural, transmembrane protein that complexes with Nsp3 and Nsp6, and is involved in double membrane vesicle (DMV) formation[125]. Together, Nsp3, Nsp4, and Nsp6 are predicted to function to nucleate and anchor viral replication complexes on DMVs in the cytoplasm[126].

We identified eight high-confidence host protein interactions with Nsp4. Nsp4 interacts with IDE, which is involved in intercellular signaling through the cellular breakdown of diverse signaling peptides[127,128]. IDE is also involved in antigen processing through the production of an antigenic peptide that is presented to cytotoxic T lymphocytes by MHC class I[129]. IDE acts as an entry receptor for varicella-zoster virus (VZV), where VZV glycoprotein E interacts with IDE through its extracellular domain[130,131]. Nsp4 interacts with various mitochondrial proteins linked to transport, including TIMM10, TIMM10B, TIMM9 and TIMM29, members of the TIM22 complex. TIMM22 facilitates the import and insertion of multi-pass transmembrane proteins into the mitochondrial inner membrane, with TIMM9 additionally involved in protein homodimerization and chaperone binding[132]. Nsp4 also interacts with DNAJC11, which is required for mitochondrial inner membrane assembly and functions through involvement with the MICOS complex and the MOM sorting assembly machinery complex[133]. Other interactions include ALG11, a mannosyltransferase catalyzing oligosaccharide linkage[134,135] and NUP210, a nuclear pore membrane glycoprotein involved in nuclear pore assembly and fusion, nuclear pore spacing, and structural integrity[136,137].

**SARS-CoV-2 Nsp5/Nsp5_C145**

Nsp5 encodes the coronavirus main protease ($M^{pro}$) responsible for cleaving itself and the other subunits from the polyproteins Orf1/1ab[138]. As these proteins include the replicase machinery, Nsp5 $M^{pro}$ is essential for all coronaviruses, and indeed is functionally and structurally conserved throughout order *Nidovirales*[139]*.* The catalytic residues of SARS-CoV align to H41 and C145 in our SARS-CoV-2 construct. We have used as baits both catalytically active Nsp5 as well as a catalytically dead C145A mutant.

Notably, we see the catalytically active Nsp5, but not the C145A mutant, interact with Histone Deacetylase 2 (HDAC2). HDAC2 is known to regulate genes of the inflammatory response, especially in the pulmonary context, where low HDAC2 expression contributes to increased disease severity in chronic obstructive pulmonary disease[140].

**SARS-CoV-2 Nsp6**

Nsp6 complexes with two other transmembrane proteins, Nsp3 and Nsp4, to form double membrane vesicles (DMV), anchoring viral replication complexes inside[141–143]. Chemical inhibition and targeted mutagenesis studies have shown that these DMVs are crucial to viral replication and are formed early after viral entry into the cell[144–147]. The most well-characterized function of the Nsp6 protein is limiting autophagosome expansion, which likely benefits the virus by preventing its components from being sent to the lysosome for degradation[148]. Further, in the context of IBV, it was shown that the generation of autophagosomes by Nsp6 can be induced by chemical inhibition of the mTOR pathway[148]. While this behavior has been shown for SARS-CoV Nsp6 (showing 95% similarity to SARS-CoV-2 Nsp6) and related viruses such as MSV and IBV, its exact mechanism and whether SARS-CoV-2 Nsp6 will function similarly remains uncertain.

In this study, we identified four high confidence interactors of SARS-CoV-2 Nsp6 (ATP5MG, ATP6AP1, ATP13A3, and SIGMAR1). Notably, three out of four of these prey proteins are subunits of different ATP synthases, suggesting that Nsp6 may be manipulating the metabolic program of the cell through ATP synthases of different organelles. As viral infection requires a new surplus of energy, it is unsurprising that viruses hijack host metabolic machinery to carry out a wide variety of cellular functions[149–152]. Specifically, ATPases are crucial for budding by multiple viruses, including HIV-1, IAV, and Ebola[153–155]. One such target is vacuolar ATPases (vATPases), which are responsible for modulating endo-lysosomal acidification and thus a number of processes like membrane trafficking and protein degradation[156]. In the context of IAV infection, vATPase ATP6V1A was identified as a key host dependency factor for viral replication across multiple RNAi knockdown screens[48,157]. Influenza A virus is believed to leverage this interaction to lower the pH of the lysosomal environment to accelerate the process of viral uncoating in the early stages of viral infection[158,159]. ATP6AP1, a subunit of the vATPase protein pump and high-confidence physical interactor of Nps6 in this study, has also been described as a host factor for a number of viruses, including IAV, WNV, and DENV[47,48,157,160,161]. Further, dysregulation of AT6AP1 leads to impaired vesicle acidification and intracytoplasmic granules, resulting in multiple clinical pathologies including an immunodeficiency syndrome and granular cell tumors[162,163]. Given the known role of ATP6V1A as an IAV host factor, the mechanism of SARS-CoV-2 Nsp6 could leverage its interaction with ATP6AP1 similarly to speed replication in DMV complexes.

ATP13A3 is a poorly understood cation-transporting P-type subunit of the mitochondrial ATPase that has recently been implicated in both host innate immunity to pathogens and pulmonary pathologies. In the case of HSV-1, ATP13A3 was shown to be less abundant in the plasma membrane of infected cells, but its expression could be rescued by the deletion of a specific HSV-1 protein pUL56, suggesting that it may be specifically targeted as a host factor in the context of some viral infections[164]. Further, ATP13A3 was identified in the plasma membrane of primary CD4+ T cells infected with Vpu-deficient and Nef-deficient HIV-1, and was differentially expressed in CD8+ T cells during IAV infection[165]. More broadly, ATP13A3 expression was found to be a significant variable in determining interindividual response to lipopolysaccharide sensing in human dendritic cells[166]. While ATP13A3 is lowly expressed in the lungs (though more highly expressed in the nasopharynx and bronchus), a rare variant of ATP13A3 is known to cause pulmonary arterial hypertension. It was recently reported that loss of function of *ATP13A3* results in disruption of polyamine homeostasis in pulmonary arterial endothelial cells, leading to endothelial dysfunction and ultimately pulmonary arterial hypertension[167,168]. Notably, ATP13A3 is also amenable to direct drug targeting.

#### SARS-CoV RNA polymerases: Nsp7, Nsp8 and Nsp12

SARS-CoV Nsp7, Nsp8 and Nsp12 proteins form the RNA-dependent RNA polymerase (RdRp)[169,170]. Eight Nsp7 proteins complex stoichiometrically with eight Nsp8 proteins to form a hexadecameric barrel-shaped structure thought to surround double-stranded RNA[171]. Mutations in Nsp8 that disrupt interaction with Nsp7, RNA, or Nsp12 are lethal to SARS-CoV, demonstrating the function of Nsp8 is essential. The catalytic activity of the Nsp7-Nsp8 complex is much weaker than that of Nsp12 though it is primer-independent, and RNA products of Nsp7-Nsp8 complex are typically $\leq$6 bases. Thus it is believed that the Nsp7-Nsp8 complex functions as a primase, generating short RNA primers for Nsp12's more processive, primer-dependent RNA polymerase activity[172,173].

Nsp12 encodes a canonical, primer-dependent RdRp domain in its C-terminus[172,174], and is essential for viral RNA synthesis. A Nsp7-Nsp8 heterodimer binds to and stabilizes loops in the polymerase domain of Nsp12, and is thought to facilitate interactions between Nsp12 and RNA synthesis/processing machinery[170]. Nsp12 contains an unusually large N-terminal extension with a conserved, essential nucleotidyltransferase domain[175] and protein interface domain that binds a second Nsp8[170]. The full functions of these domains in virus replication or fitness and host biology are not known. Nsp12 may have additional roles outside of the RdRp complex mediated through interactions with the uncharacterized N-terminus[170,172,175] and/or other virus proteins including Nsp5, Nsp8 and Nsp9[176].

The structure of SARS-CoV-2 Nsp7/Nsp8/Nsp12 complex closely mimics that of SARS-CoV[177], and the complex members bear 98.8%, 97.5%, and 96.4% genetic similarity at the codon level respectively, suggesting that function will be conserved.

**SARS-CoV-2 Nsp7**

We identified 32 high-confidence host protein interactions with Nsp7. Groups of Nsp7 interactors include multiple Rab GTPases with various roles in exocytic and endocytic membrane trafficking, mitochondrial proteins such as cytochromes, and several other factors that were previously identified in the host interactomes of other viruses, yet without defined roles in infection.

Rab GTPases function in networks of crosstalk[178]. Among the identified Rab GTPases, Rab14 regulates the membrane recycling pathway from endosomes to the plasma membrane which is required for ADAM protease trafficking and regulation of cell-cell junctions[179]. Rab14 was previously characterized to be required for HIV-1 envelope glycoprotein particle incorporation[180]. Rab1a regulates vesicular protein transport from the ER to the Golgi and was identified to be required for production of extracellular enveloped virions of VACV[181], as well as for CSFV particle assembly[182]. Rab7a and Rab5c are essential for HBV infection[183]. Rab7a is implied to promote virus entry in early stages[183] and restrict exocytic virion release in late stages[184]. Rab8a was shown to be an important regulator of HIV-1 trafficking to the virological synapses[185]. Virological synapses enable delivery of the virus to CD4+ T-cells.

ACSL3-LPIAT1 fusion protein is a known cancer gene and ACLS3 is highly expressed in human lung cancer[186]. ACSL3 induction results in increased acyl-CoA synthesis that is essential for providing the prostaglandin required for the maintenance of non-small cell lung cancer[187] and mutant KRAS lung cancer tumorigenesis in vivo[186]. Besides the function of ACSL3 in lung cancer, it is a host factor that interacts with several viral proteins and is required for poliovirus replication[188–190].

MOGS is another host factor that was identified in the interactomes of various virus proteins, including HBV, HBe, HCV core, HIV VPR and gp120 and Mtb Ppe11[189,191,192]. This gene is well characterized in the context of a congenital disorder of glycosylation. Interestingly, patients with genetic defects in MOGS manifest decreased susceptibility to viral infections[193], providing a potential target for broad spectrum therapy of viral infections.

**SARS-CoV-2 Nsp8**

We find that Nsp8 interacts with 23 host proteins, a number of which are known host-dependency factors of various pathogens. For example, five of the ten members of the exosome complex are identified here as high confidence Nsp8 interactors, with the remaining five just narrowly missing our stringent threshold. LARP7, a member of the 7SK snRNP RNA complex, interacts with WNV, ZIKV capsid, and HIV-1 Tat in such a way as to compete with viral proteins for the transcriptional elongator pTEFb. The interaction of LARP7 and coronavirus primase points to a mechanism by which the virus promotes elongation of its vRNA.

Interestingly, we find that Nsp8 interacts with proteins just under our scoring threshold found to be differentially regulated or expressed during acute lung disease secondary to pathogen infection, including CEBPZ, DAP3, NKRF and ZN512. NFKR for example is upregulated in the circulating monocytes and alveolar macrophages of patients with active pulmonary TB, and NKRF serves as an endogenous repressor for IP-10 and IL-8 synthesis to hinder host from robust response to MTb infection[194,195].

**SARS-CoV-2 Nsp12**

In our study, SARS-CoV-2 Nsp12 interacts with 20 high confidence host proteins. Consistent with Nsp12 RdRp activity, eight host protein interactors are RNA binding factors involved at multiple steps of RNA processing and regulation. These host proteins could facilitate long-range RNA interactions that occur during genome replication and discontinuous transcription[196], or mediate viral protein translation. Five of these host interactions are identified with proteins from other RNA and DNA viruses[36,190]. Of note, Nsp12 interacts with three proteins involved in pre-mRNA splicing: A-kinase anchor protein 8 (AKAP8)[197]; and spliceosome components pre-mRNA-splicing factor SLU7 (SLU7)[198] and peptidyl-prolyl cis-trans isomerase-like 3 (PPIL3). Notably, siRNA knockdown of SLU7 inhibits early stages of HIV-1 replication[199]. Nsp12 also interacts with La-related protein 4B (LARP4B), a cytoplasmic RNA binding protein that promotes mRNA translation and interacts with PABPC1 and RACK1 kinase, potentially connecting 3' mRNA factors with translation machinery[200]. PABP, a 3' poly(A) tail binding protein, is a cis acting element on coronavirus RNA that is essential for bovine coronavirus replication[201,202]. Nsp12 could be targeting LARP4B as a bridge to recruit members of the PABP family for efficient RNA replication and translation. Nsp12 interacts with two RNA binding proteins that localize to and nucleate formation of stress granules: ubiquitin-associated protein 2 (UBAP2) and ubiquitin-associated protein 2-like (UBAP2L)[203]. UBAP2L interaction is identified in our data with SARS-CoV-2 Nsp9, although it falls below our scoring threshold. Coronavirus species mouse hepatitis virus Nsp9 has been shown to interact with Nsp12[176]. This may suggest a potential role for SARS-CoV-2 Nsp12 and Nsp9 in coordinating host and/or viral RNA regulation (see also Nsp9), as stress granules are sites of RNA storage and intermediate stages between translation and mRNA decay[204]. Additionally, since MERS-CoV has been shown to inhibit stress granule formation to promote viral replication[89], SARS-CoV-2 Nsp12 may interact with and sequester UBAP2L as a potential mechanism to inhibit stress granule nucleation and promote virus replication (see also N protein for role of stress granules).

Although coronavirus genome replication and transcription occurs in the cytoplasm, SARS-CoV-2 Nsp12 interacts with five nuclear, DNA-related factors: transcription factors CREB-regulated transcription coactivator 3 (CRTC3), transcription factor 12 (TCF12) and zinc finger protein 318 (ZNF318); chromatin factor SBNO1; and AKAP8 which regulates histone methylation and gene expression[205]. Nsp12 shows DNA-dependent activity and can synthesize nucleotides from a DNA template *in vitro*[172,173]. Nsp12 may have novel roles in chromatin and transcription regulation, although the advantage to the virus is unclear.

Nsp12 also interacts with host proteins that regulate inflammatory signaling and apoptotic pathways. Nsp12 interacts with receptor-interacting serine/threonine-protein kinase 1 (RIPK1) with very high confidence. As an active regulatory kinase, RIPK1 triggers cell death by apoptosis or necroptosis; as a scaffold independent of its kinase activity, RIPK1 regulates inflammatory signaling and inhibits cell death[206–208]. Many diverse viral and bacterial proteins interact with and/or modify RIPK1 to modulate host defense pathways, cytokine signaling and/or host cell death[209–211]. For example, HIV-1 protease cleaves and inactivates RIPK1, which then impairs host defense pathways[211]. Importantly, RIPK1 is a druggable target, and many inhibitors are being tested as anti-inflammatory treatments for neurodegenerative diseases and cancer[212]. However, given the two faces of RIPK1 regulating apoptosis for pathogen clearance[210] or stimulating cytokine production as part of the inflammatory response[208], the usefulness of drugs targeting RIPK1 as a treatment for SARS-CoV-2 will depend on future research identifying which pathway is engaged by Nsp12. In addition, Nsp12 interactor AKAP8 binds and shuttles caspase 3 to the nucleus as part of caspase-mediated proteolysis and apoptosis[213]. These interactions suggest additional novel roles for Nsp12 outside of canonical RdRp activity in regulating host inflammation and cell death, perhaps mediated through the large uncharacterized N-terminal domain of Nsp12.

**SARS-CoV-2 Nsp15**

In SARS-CoV, Nsp15 has uridine-specific endoribonuclease (endoU) activity, and is essential for viral RNA synthesis[214]. The Nsp15-associated endoU domain is one of the most conserved proteins among CoVs and related viruses, suggesting important functions in the viral replicative cycle. The endoUs were shown (i) to have endonucleolytic activity, (ii) to cleave 3' of pyrimidines, preferring uridine over cytidine, and (iii) to release reaction products with 2'-3'-cyclic phosphate and 5'-OH ends. Nsp15s were shown to form homohexamers composed of a dimer of trimers[215]. Deletion of a conserved domain for this enzymatic activity led to loss of viral RNA generation measured by RT-PCR[214]. Nsp15 is also known to be critical for evasion of host dsRNA sensors in macrophages[216]. In our map, Nsp15 interacts with three host proteins. NUFT2 mediates the import of GDP-bound RAN from the cytoplasm into the nucleus, and thus indirectly plays a more general role in cargo receptor-mediated nucleocytoplasmic transport[217,218]. ARF6 is a GTP-binding protein involved in protein trafficking that regulates endocytic recycling and cytoskeleton remodeling[219–223] as well as the activation of cholera toxin[224]. RNF41 acts as E3 ubiquitin-protein ligase and promotes TRIF-dependent production of type I interferon and inhibits infection with vesicular stomatitis virus[225].

**SARS-CoV-2 Nsp9**

In SARS-CoV and related coronaviruses, Nsp9 is an essential non-structural protein that binds to RNA and DNA, with a preference for single-stranded RNA[226–229]. The function of Nsp9 is not well annotated, though in SARS-CoV it is believed to interact with Nsp8 and the viral replication complex (Nsp7, Nsp8, and

Nsp12)[229,230] . In our hands, SARS-CoV-2 Nsp9 interacts with 16 high confidence host proteins, with many having been shown in previous studies to regulate nuclear transport, transcription, and mRNA degradation in response to various viral infections. Unexpectedly, Nsp9 also demonstrates some strong interactions with extracellular matrix proteins involved in elastin formation, lung development, and lung injury and repair.

Nsp9 interacts with Nup62, Nup58, and Nup54, the three components of the Nup62 subcomplex which forms the channel of the nuclear pore complex (NPC)[231–235]. In addition, Nsp9 interacts with nucleoporins on the cytoplasmic side of the nuclear pore complex (i.e. NUP214 and NUP88) but none from the nucleoplasmic side[236] indicating a cytoplasmic role of Nsp9 at the NPC. Many viruses have been shown to exploit cellular nuclear transport machinery to block host-related transport or promote viral transport in a manner beneficial to the virus[237–250]. Nup62 in particular interacts with a number of viral proteins including EBV BGLF4 protein kinase[251], HPV16 and HPV8 E7 proteins[252,253], and HIV-1 IN[254] . Several positive-sense single strand RNA viruses (e.g. poliovirus, EV71, rhinovirus, and cardioviruses) target Nup62 either through viral protease-directed cleavage[240–246] or induced hyperphosphorylation[247–249] in order to limit nuclear transport. Vaccinia virus, a DNA poxvirus, replicates in cytoplasmic virus factories that recruit G3BP1 and Nup62[250]. Nup54 interacts with Influenza A virus polymerase and was shown to be important for virus replication and transcriptional activity[255]. In addition to the nucleoporins, Nsp9 interacts with MIB1, a RING-type E3 ubiquitin ligase shown to act as a dependency factor during adenovirus infection[256]. Adenoviruses are non-enveloped DNA viruses that utilize the NPC to dock and deliver genomic cargo into the nucleus, and in this study, the authors demonstrate that MIB1 mediates the delivery of viral DNA through the NPC[256].

In addition to nuclear transport machinery, Nsp9 interacts with three transcription regulators, Nek9, DCAF7 and eIF4H. Nek9 is a serine/threonine kinase that regulates mitotic progression[257]. It has been shown as an adenovirus dependency factor promoting viral growth, interacting with E1A protein to silence the expression of certain host genes, and was demonstrated to colocalize at adenovirus replication centers[258]. DCAF7 is a potential substrate adaptor for CUL4-DDB1 E3 ligase[259] that has also been shown to interact with adenovirus E1A protein to suppress innate immune response and depresses IFN stimulated genes (ISGs)[260]. In combination with eIF4A, eIF4H was shown to interact with HSV virus host shut off proteins, with eIF4A shown to help degrade mRNA and switch from host to viral gene expression[261–264]. Taken together, these interactors suggest a potential function of Nsp9 in the inhibition of host mRNA expression potentially to limit the express of ISGs.

Somewhat unexpectedly, Nsp9 interacts with several proteins involved in lung development, lung injury and repair, and lung cancers. The most abundant interactor in the Nsp9 interactome is Fibrillin-2 (FBN2), an extracellular matrix glycoprotein protein involved in elastin fiber and respiratory organ development[265]. Additional related interactors were identified, albeit with lower abundance. Fibrillin-1 (FBN1) and Fibulin-5 (FBLN5) are both extracellular matrix proteins, also involved in elastin fibre development. Mutations in FBN1 cause Marfan syndrome (MFS), a connective tissue disease that can result in early morbidity and mortality mainly caused by aortic aneurysm and rupture, though additional clinical manifestations include lung complications[266]. FBLN5 is indicated to serve a role during lung injury and repair, is frequently silenced in lung cancer, and has been shown to suppress cell invasion[267–272].

The Nsp9 interactome reveals a potential function for the protein not only in viral replication, but potentially in regulating nuclear transport, though it is unclear if this regulation would block host cell transport or promote viral transport. In addition, the interaction with transcription regulators indicates a potential role in inhibition of host cell transcription and potentially host shut off. And finally, the unexpected interaction with fibrillins-1 and -2, and fibulin-5 could implicate Nsp9 as a potential complicating factor during disease pathogenesis, and may point to additional molecular reasons underlying SARS-CoV-2 complications in the lungs.

**SARS-CoV-2 Nsp10**

Nsp10 contains two Zn-finger motifs, binds nucleic acids non-specifically, and has been implicated in minus-strand RNA synthesis, thus performing an essential role in viral replication[273,274]. A unique feature for SARS-CoV is that Nsp16 requires Nsp10 as a stimulatory factor to execute its methyltransferase activity[275].

Nsp10 on its own forms a dodecameric homomeric complex[276] which excludes its interfaces with Nsp14 and Nsp16, meaning that Nsp10 could have at least two different functional quaternary structures. In our map, we identify five high-confidence host protein interactions with Nsp10. Nsp10 interacts with two subunits of the clathrin adaptor protein complex 2 (AP-2), AP2A2 and AP2M1. This interaction is reminiscent of the Nef protein from Human and Simian Immunodeficiency Viruses which are shown to bind the AP-2 complex through a canonical AP-2 recognition acidic dileucine motif ([RQED]XXXL[LIV])[277]. HIV and SIV are thought to use the interaction between Nef and AP-2 to hijack the clathrin machinery and endocytose host proteins such as CD4 and MHC-I[278]. Interestingly, coronavirus Nsp10 proteins do not appear to contain the canonical AP-2 binding motifs (the acidic dileucine motif [RQED]XXXL[LIV] for binding AP2A2 nor YXXΦ for binding AP2M1).

Nsp10 also interacts with GFER which was identified as a host dependency factor for West Nile and Dengue virus infections [47]. Two other Nsp10 interactors ERGIC1 and GRPEL do not have a known role in viral pathogenesis. ERGIC1 plays a possible role in transport between endoplasmic reticulum and Golgi[279]; and GRPEL1 participates in the translocation of transit peptide-containing proteins from the inner membrane into the mitochondrial matrix[280].

**SARS-CoV-2 Nsp11**

SARS-CoV-2 Nsp11 is a short peptide (13 aa) at the end of Orf1a, and it is not clear if Nsp11 encodes a functional protein. Nsp11 was only found to interact with two proteins, ERP29 and TBCA. ERP29, a PDI like protein localized to the ER which plays a role in processing of secretory proteins within the endoplasmic reticulum (ER). It has been shown ERP29 triggers a conformational change and exposes the C-terminal arm of polyomavirus VP1 protein, leading to formation of a hydrophobic particle that binds to a lipid bilayer. Therefore, this ER protein mediates the penetration of polyomavirus across the ER membrane[281].

#### SARS-CoV Capping Enzymes: Nsp13, Nsp14 and Nsp16

The m7GpppN (N any nucleotide) cap of mRNA promotes translation. SARS-CoV Nsp13, Nsp14, and Nsp16 encode enzymes that install cap structure onto mRNA[282,283]. The pathway of cap synthesis on nascent viral mRNA is thought to be similar to capping of cellular mRNA, but occurs in the cytoplasm instead of the nucleus[282,284,285].

**SARS-CoV-2 Nsp13**

Nsp13 is essential for SARS-CoV viral RNA synthesis[286], and SARS-CoV-2 Nsp13 shares 100% amino acid similarity with SARS-CoV. Nsp13 is a helicase/triphosphatase, and triphosphate cleavage initiates the first step in mRNA capping[287]. Nsp13 hydrolyses the gamma phosphate of nascent mRNA and the resulting disphophate is then converted to a GpppN RNA by a yet to be identified guanylyl transferase.

SARS-CoV-2 Nsp13 was found to interact with 40 host proteins, from Protein Kinase A (PKA) signalling, to the Golgi apparatus, to multiple members of protein complexes associated with microtubules/centrosomes. SARS-CoV-2 Nsp13 showed a strong interaction with Giantin (GOLGB1), which has previously been shown to interact with herpes simplex virus type 1 (HSV-1) UL37[288] and the Tick-borne encephalitis virus (TBEV) replicon[289]. Interestingly, in macaque experiments expression of the HIV protein Nef resulted in increased GOLGB1 levels and led to Golgi disruption and specific pulmonary vasculopathies[290].

Unexpectedly, we found that SARS-CoV-2 Nsp13 pulled down both the regulatory (PRKAR2A and PRKAR2B) and the catalytic (PRKACA) subunits of PKA as well as the A-kinase anchoring protein AKAP9 and the phosphodiesterase interacting protein PDE4DIP. AKAP9 (also called AKAP450) is a large scaffolding protein that localizes to the Golgi apparatus and centrosomes[291,292] where it assembles multiple signaling proteins (e.g., PKA and PDE4D) that control microtubule organization[293], polarized secretion[294], Golgi morphology[294,295], ciliogenesis[296], directional cell migration[297] and cell cycle progression[292]. Importantly, the activities of PKA signaling complexes have been implicated in multiple membrane transport steps[298–301], suggestive of a role for SARS-CoV-2 Nsp13 in hijacking the host secretory pathway for viral benefit. Also notably, a pool of AKAP9 relocalizes to RNA stress granules upon treatment with arsenite where it forms a complex with G3BP and CCAR1 and regulates stress granule size and composition[302]. PKA, being a kinase, is also targetable by small molecule inhibitors or peptides that target the AKAP-PKA binding interface.

An additional SARS-CoV-2 Nsp13 interaction partner was the endosomal transport protein ERC1. Knockout of ERC1 causes a significant decrease in dengue virus replication, and interestingly a similar phenotype was observed with additional SARS-CoV-2 Nsp13 pulldown target GOLGA2[303]. The NS3 protein of hepatitis C virus binds to ERC1 and may mediate the pathogenesis of HCV[304], and knockdown of ERC1 significantly decreases human cytomegalovirus (HCMV) viral production[305]. Of particular interest, ERC1 has previously been identified as a potential drug target in dengue virus infection[306].

**SARS-CoV-2 Nsp14**

Nsp14 is a bifunctional enzyme. It encodes an exonuclease (exo) domain that corrects mutations that arise during genome replication[307]. In addition, a separate domain of Nsp14 functions as a SAM dependent methyltransferase (MTase) that generates the N-7 Guanosine of the m7GpppN cap on viral mRNA. High-resolution crystal structures of Nsp10/Nsp14 suggest exo and MTase activity is stimulated by Nsp10 through an allosteric mechanism[308]. There are three host factors that copurify with Nsp14. GLA is an alpha galactosidase implicated in Fabry Disease[309]. Migalastat, a pharmacological chaperone, targets GLA through the inhibition of alpha-glucosidase and glycosylation, increasing its lysosomal activity[310]. SIRT5 is a mitochondrial protein linked to metabolism and aging that removes malonyl, succinyl, acetyl, and glutaryl groups on lysines of target proteins[311–314] Numerous compounds target SIRT5, including HDAC inhibitors[315]. IMPDH2 catalyzes the conversion of isosine 5' phosphate (IMP) to xanthine 5'-phosphate (XMP) which is then converted into guanine 5' monophosphate for *de novo* synthesis of guanine nucleotides[316]. It is tempting to speculate that the copurification of Nsp14 and IMPDH2 reflects an interplay between Nsp14 activities and purine nucleotide metabolism. Merimepodib, a nucleoside analog and broad spectrum antiviral, is among the compounds targeting IMPDH2[317].

#### SARS-CoV Open Reading Frames

**SARS-CoV-2 Orf3a**

Orf3a is the largest (274 aa) group-specific Orf in the SARS-CoV-2 genome. It is thought to be non-essential but has multiple key functions in viral pathogenesis, from mediating the trafficking of SARS-CoV Spike (S protein) to inducing apoptosis and inflammation during infection[318–323]. Specifically, Orf3a is a type IIIa integral membrane protein thought to induce pro-IL-1β expression and protein maturation through the TRAF3-dependent ubiquitination of ASC and p105, ultimately activating NF-κB and the NLRP3 inflammasome[319]. Despite its clear functional significance as an accessory protein, Orf3a is not believed to be critical for the formation of viral particles[320], though it has been been shown to upregulate the secretion of fibrinogen in lung epithelial cells, which is responsible for the induction of cytokine storm, particularly in the respiratory tract[324]. Within the cell, Orf3a localizes to the perinuclear region and plasma membrane, forming punctae throughout the cytoplasm as it complexes with cellular factors [318,325,326] . In the rER and Golgi, Orf3a has been shown to cause substantial ER stress during infection[327]. A YxxΦ motif present in the C-terminal cytoplasmic domain of Orf3a is required for delivery to the plasma membrane, from which Orf3a is internalized and traffics through the endocytic pathway to lysosomes[328]. Orf3a can form tetramers that are proposed to act as cation-permeant ion channels[329]. In this study, Orf3a pulled down eight high-confidence host protein interactors (ALG5, ARL6IP6, CLCC1, HMOX1, SUN2, TRIM59, VPS11, VPS39) with functional enrichments in autophagy (HMOX1, VPS11, VPS39) and organelle localization (HMOX1, VPS11, SUN2). VPS11 and VPS39 serve as members of the HOPS and CORVET complexes, respectively, which coordinate fusion of the lysosome with the endosome and autophagosome[330,331].

HMOX1, a key enzyme in heme catabolism, is one of the most promising of these physical interactors. This oxygenase cleaves free heme to produce iron, biliverdin and carbon monoxide, which provides a cytoprotective effect to cells as excess free heme induces apoptosis[332]. In turn, this elicits a cascade of physiological events, most notably the induction of anti-inflammatory cytokines IL-10 and IL-1RA[333,334]. Given these features of HMOX1 and its high expression in the lungs, it has been implicated in a broad range of disease states, including diabetes, heart failure, lung carcinoma and COPD[335,336]. In the context of infection, upregulation of HMOX1 has shown to have a protective effect against the oxidative stress of a whole host of pathogen infections, including viruses like HIV, DENV, HCV, and IAV, as well as parasites and *Mycobacterium* species[337,338]. Importantly, this protein is biochemically tractable, making it amenable to direct targeting with known pharmacologic agents. As severe inflammatory response in the respiratory tract is a key clinical feature of SARS-CoV-2 infection, further exploration of HMOX1 could elucidate mechanisms of SARS-CoV-2 pathogenesis and a path forward for treatments.

Another exciting protein candidate identified in this study, CLCC1, is an intracellular chloride channel that is highly expressed in the lung, heart, and a number of other tissues. We identified a very strong, high-confidence physical interaction between CLCC1 and Orf3a. Both CLCC1 and SARS-CoV Orf3a are localized to the ER membrane, where loss of *CLCC1* results in ER stress and disruption of the protein folding capacity of the ER, leading to misfolded protein accumulation and well-characterized retinal cell dysfunction in the clinic[339,340]. In a similar vein, expression of *CLCC1* has been correlated with volume of adipose tissue in HIV-infected men[341]. Given the similar roles in solute transport, similar localization in the ER, and evidence of CLCC1 interaction with NS1 of Human Respiratory Syncytial Virus (RSV), there is a high potential for not only physical but also functional interaction of CLCC1 and Orf3a in the context of SARS-CoV-2 infection[342]. Interestingly, CLCC1 also interacts directly with SLC15A3, a druggable peptide/histidine transporter and known interferon-stimulated gene that is regulated by TLR-activation and contributes to TLR4-mediated inflammation in macrophages and epithelial cells[343,344]. While SLC15A3 was not identified as a direct interactor of Orf3a, we identified a number of high-confidence interactions between SARS-CoV-2 proteins and members of the solute carrier superfamily such as SLC25A17 and SLC6A15, suggesting that Orf3a may play a role in modulating host innate immunity via an indirect interaction with CLCC1 or as part of a larger protein complex or pathway.

In addition to HMOX1 and CLCC1, Orf3a also interacts with known viral host factors and interactors. ALG5, an ER-localized glycosyltransferase involved in N-glycan biosynthesis, has been characterized as a cellular host factor for Influenza A virus replication in multiple genome-wide CRISPR-Cas9 knockout screening efforts, along with other ALG family proteins[51,52]. Taken together, our data and prior studies suggest a potential role of these ER-localized glucosyltransferases in modulating viral replication and pathogenesis, potentially through manipulation of viral particle formation. Similarly, ARL6IP6, a transmembrane protein responsible for ADP ribosylation, is known to physically interact with both HPV and WNV, and was differentially expressed in CD8+ T cells from HIV+ progressors on HAART[36,345,346].

**SARS-CoV-2 Orf3b**

SARS-CoV-2 Orf3b encodes a 168 aa protein that is not well conserved (9.5% codon similarity as compared to SARS-CoV Orf3b). While Orf3b is thought to be non-essential[1,320], it has been shown to be an IFN antagonist in both SARS-CoV and SL-CoV (bat)[347,348] and to subsequently be involved in pathogenesis[1]. The only confident interaction partner detected for Orf3b was the mitochondrial protein STML2/STOML2. STOML2 stimulates cardiolipin biosynthesis and recruits and stabilizes prohibitin. Both STOML2 and its interactor prohibitin have been shown to be host dependency factors for Enterovirus 71 neuropathogenesis[349]. STOML2 forms large complexes with the i-AAA protease YME1L and the rhomboid protease PARL at the inner mitochondrial membrane, which regulate key proteins of the mitochondrial stress response such as PGAM5 and PINK1[350]. STOML2 may directly be involved in regulating T-cell-mediated immune responses by modulating T-cell receptor activation[351].

**SARS-CoV-2 Orf6**

Orf6 is not essential for the replication of SARS-CoV, but affects viral production[352]. Orf6 may function as a type I IFN antagonist, suppressing IFN induction and IFN signalling pathways by binding to and sequestering karyopherin α2, which normally facilitates the nuclear import of the interferon signalling responsive transcription factor STAT1[348,353]. In SARS-CoV infected cells, Orf6 complexes with Orf9b[354]. SARS-CoV-2 Orf6 was found to interact with three host proteins: NUP98, RAE1, and MTCH1. As described in the main text, NUP98-RAE1 is an interferon-inducible mRNA nuclear export complex that is targeted by multiple viruses including VSV, IAV, KSHV, and Polio [355,356].

**SARS-CoV-2 Orf7a**

SARS-CoV Orf7 is divided into two open reading frames, designated Orf7a and Orf7b. Orf7a (also known as U122) encodes a 122-amino-acid protein and contains a compact seven-stranded β-stack similar in structure to members of the immunoglobulin superfamily[357]. Orf7a has a well-characterized localization to different locations throughout the cell, from the perinuclear region to the cytoplasm, and even contains a type-1 transmembrane domain anchoring it in the plasma membrane[358–360]. This distribution into multiple cellular regions could help explain the various functions of Orf7 (namely regulation of cell cycle progression and apoptosis) and the diversity of its interaction partners[357–360]. The role of Orf7a in apoptosis was highlighted by an increase in caspase-3 protease activity that resulted in a significant induction of apoptosis[358]. Orf7a expression has also been shown to downregulate cyclin D3, resulting in the accumulation of retinoblastoma protein (Rb) and ultimately cell cycle arrest in G0/G1 phase[359]. Further, Orf7a has been shown to both localize in the perinuclear region and colocalize with ER and ER-Golgi intermediate compartment (ERGIC) markers during infection[360]. Interestingly, this cellular localization coincides with that of high-confidence protein interactors identified in this study, MDN1 and HEATR3, as well as a factor that was just below our MIST threshold, TNPO1. While it did not satisfy our stringent scoring criteria, TNPO1, or transportin-1, is worth mentioning due to its demonstrated role in other viral infections such as Influenza A virus, HIV-1 and Hepatitis C virus, where it plays a crucial role in mediating nuclear transport of viral proteins and protein complexes[361–364] HEATR3 has also been shown to activate the NF-kB pathway via NOD-2, which has been implicated in a pro-inflammatory response during Crohn's disease[365,366] and therefore Orf7a may target HEATR3 to modulate inflammatory response upon SARS-CoV-2 infection.

**SARS-CoV-2 Orf8**

Orf8 is an accessory protein and is not essential for virus replication *in vitro* and *in vivo*[320,367]. It is one of the most rapidly evolving regions among SARS-CoV genomes and was previously shown to be a recombination hotspot[368–370]. Pairwise comparison of amino acid sequences showed that SARS-CoV-2 Orf8 exhibited 45.3% sequence similarity with SARS-CoV. Orf8 in human isolates from the 2003 epidemic contained a signature 29-nucleotide deletion compared to all civet and bat SARS-related CoVs, which causes the split of full-length Orf8 into two small proteins: 8a and 8b[371]. Orf8 from SARS-CoV-2 encodes a single polypeptide and lacks the aggregation motif VLVVL present in SARS-CoV Orf8b, which was shown to induce ER stress and activate NLRP3 inflammasomes[372]. Further, Orf8b protein has been shown to be modified by N-linked glycosylation on N81 residue, which protects Orf8ab protein from proteasomal degradation[373]. This novel Orf8 likely encodes a secreted protein and has an N-glycosylation site at N78, within the consensus sequence NYT.

We identified 47 high-confidence host protein interactions with Orf8. Several Orf8 interactors are involved in ER stress and ER-associated degradation (ERAD) pathway, including UDP-glucose/glycoprotein glucosyltransferase 2 (UGGT2), ER degradation enhancing alpha-mannosidase like protein 3 (EDEM3), OS9[374], N-glycanase 1 (NGLY1), and FAD-dependent oxidoreductase domain-containing protein 2 (FOXRED2)[375–378]. The ERAD pathway targets unassembled glycoproteins for ubiquitylation and proteasomal degradation. EDEM3 has been shown to increase ubiquitylation of HCV envelope proteins via direct physical interaction and consequently reduce viral production[379]. OS9 and ERLEC1 proteins are also known to be targeted by other virus-encoding proteins from Dengue virus[190], HIV[189], WNV[36], HPV and KSHV[380], suggesting common molecular mechanisms of infection and proliferation used by these different pathogens.

Infection with the SARS virus results in severe inflammation in the lungs, which can lead to respiratory distress and fibrosis during the late stages of infection[381]. Fibroblast activation and overexpression of collagen are two important aspects of the pathogenesis of lung fibrosis. Interestingly, we identified a number of Orf8 interactors implicated in pulmonary fibrogenesis including FKBP10[382], GDF15[383], NEU1[384] and IL17RA[385]. In addition, the expression levels of ADAMTS1[386] and HS6ST2[387] are modulated during lung inflammation and fibrosis and are identified as Orf9b interactors. Growth differentiation factor 15 (GDF15) is a fibroblast-inhibiting cytokine that inhibits the growth and activation of lung fibroblasts by inactivating the TGF–Smad pathway, suggesting this cytokine could be a potential therapeutic for ameliorating interstitial lung fibrosis during severe SARS infection. FK506-binding protein 10 (FKBP10) is a collagen chaperone and inhibition attenuates expression of profibrotic mediators and effectors[382], suggesting that this protein can be targeted to reduce virus-induced lung fibrosis. Activation of the pro-inflammatory cytokine receptor IL17RA in lung tissues is an important host defense mechanism upon fungal, bacterial and viral infections, but its overactivation increases collagen secretion and exacerbates pulmonary fibrosis[385]. Inhibition of IL17RA signaling promotes resolution of pulmonary inflammation and fibrosis in *in vivo* models[388] and may therefore serve as a therapeutic strategy to reduce lung fibrosis during SARS infection.

Though below our stringent scoring threshold, one additional interesting protein identified in Orf8 pull-downs was the cellular guanyl transferase RNGTT/Mce1. This interaction was significant (>0.5 BFDR) but just below our MIST threshold (MIST score = 0.649). Previous studies suggest RNGTT can exist in the cytoplasm[389,390], it is possible that Orf8 recruits RNGTT to viral mRNA to add G to nascent mRNAs after they are acted on by Nsp13 to make GpppN mRNA that is subsequently acted on by Nsp14 and Nsp16 (see section on SARS-CoV Capping Enzymes).

**SARS-CoV-2 Orf9b**

Orf9b is an accessory protein synthesized from an alternative complete reading frame within the viral N gene, which encodes for a 98-aa long protein. Orf9b has been shown to be expressed in SARS-CoV-infected cells and antibodies against Orf9b were detected in the sera from convalescent-phase SARS patients[391,392], however the function of Orf9b is largely unknown. It is known that Orf9b can passively diffuse into the nucleus and is actively exported via Crm1-mediated nucleocytoplasmic export[393]. In addition, Orf9b localizes to mitochondria and causes mitochondrial elongation by inducing ubiquitination-mediated proteasomal degradation of the main pro-fission factor (DRP1) dynamin-like protein 1[394]. Orf9b targets the mitochondrial-associated adaptor molecule MAVS signalosome by utilizing PCBP2 and the HECT domain-containing E3 ligase AIP4, resulting in the degradation of MAVS and therefore limiting host cell interferon responses[394].

We found that SARS-CoV-2 Orf9b interacts with 11 human proteins, including with a mitochondrial import receptor, translocase of outer membrane 70 (TOM70). TOM70 is known to interact with MAVS protein upon RNA virus infection and it acts as a critical adaptor linking MAVS to TBK1/IRF3, resulting in the activation of IRF-3[395]. TOM70 also makes a dynamic protein complex with HSP90/IRF3/BAX and mediates virus-induced apoptosis[396]. Though more studies need to be done to fully flesh out this interaction, it is possible SARS-CoV-2 Orf9b may target TOM70 to modulate IRF3-mediated gene expression or apoptosis upon virus infection. Another mitochondrial protein identified as interacting with Orf9b is BCL2-associated athanogene 5 (BAG5). BAG5 inhibits mitophagy of damaged mitochondria by suppressing recruitment of Parkin to the sites of damage[397]. Several viruses trigger Parkin-dependent mitophagy to promote persistent infection and impair the innate immune response[398]. SARS-CoV-2 Orf9b might act similarly by antagonizing the function of BAG5.

In addition to mitochondrial proteins, SARS-CoV-2 Orf9b was also found to interact with CHMP2A, a member of the endosomal sorting complex required for transport (ESCRT)-III machinery[399]. CHMP2A was shown to contribute to the budding of a variety of viruses, including HIV[400], equine infectious anemia virus (EIAV)[401], and murine leukemia virus[401], suggesting a critical role for virus release. Other Orf9b interactors of interest include microtubule affinity-regulating kinases MARK1, MARK2, and MARK3. These proteins are involved in regulating microtubule dynamics and phosphorylation of tau[402], and MARK2 was also shown to regulate HIV trafficking through phosphorylation of FEZ1[403].

**SARS-CoV-2 Orf9c**

SARS-CoV-2 Orf9c (referred to as Orf9B in Wu et. al.) encodes a short polypeptide that is 70 aa in length[404]. There is some debate over whether Orf9c encodes a functional protein, or what it's function would be, as Orf9c is thought to be dispensable for virus replication[405–407]. Therefore it is unclear how clinically relevant molecular interactions for this bait would be in the context of coronavirus infection. Keeping that in mind, we are however able to express and purify Orf9c in HEK293T cells, identifying 26 high confidence human protein interactors of diverse functional enrichments. These functions include mitochondrial respiratory chain complex assembly (NDUFB9, NDUFAF1, ACAD9, ECSIT, BCS1L)[408–410], GPI-anchor biosynthesis (PIGO, PIGS, GPAA1)[411], and regulation of I-kappaβ kinase and NF-kappaβ signaling (NLRX1, F2RL1, NDFIP2).

F2RL1 is implicated in a variety of cellular processes related to the pathogenesis of respiratory viruses and pulmonary disease, including NFκB activation, cooperativity with Toll-like receptors, innate immune recruitment and activation, and acute lung inflammation[412–416]. In the context of IAV pathogenesis of monocytes and macrophages, F2RL1 activation protects against viral infection through an IFN-gamma-mediated mechanism[417,418]. Importantly, F2RL1 is the target of four known pharmacologic agents AC-55541, AZ8838, GB110 and Z3451. Another interactor linked to pulmonary function is NLRX1, an attenuator of Influenza A virus-induced inflammation[419]. During IAV infection, NLRX1 promotes type I IFN signaling and macrophage survival[419]. It is an essential moderator of macrophage immunity, as it senses the extent of viral replication and maintains a protective balance between antiviral immunity and excessive inflammation within the lungs[419].

In our study, Orf9c is also shown to interact with MRP1 (encoded by *ABCC1*), a multifunctional ATP-binding cassette protein that, among other diverse functions, controls the ATP-dependent efflux of drugs from the cell. It has been implicated in multidrug resistance, viral pathogenesis, and pulmonary disease, and is directly targetable by FDA-approved pharmacologic agents daunorubicin and mitoxantrone[420]. MRP1 has a demonstrated role in both HIV and CMV biology, is found to be differentially expressed in a polarized subset of macrophages during HIV-1 infection, and is associated with CMV latency[421–423]. Interestingly, HIV-1 protease inhibitors saquinavir, rotinavir, and nelfinavir are substrates of MRP1, though it was found that MRP1 did not affect the antiviral activity of these drugs in cell lines[424]. The role of MRP1 in determining the severity of diseases (e.g., COPD, pneumonia, and lung carcinoma) and multidrug resistance in the lung has been well-characterized, which could be particularly relevant given the clinical manifestation and treatment of ARDS and pneumonia during SARS-CoV-2 infection[425–428].

**SARS-CoV-2 Orf10**

SARS-CoV-2 Orf10 codes for a peptide only 38 aa long and does not have a homolog in SARS-CoV. There is no data yet providing evidence that the protein is expressed during SARS-CoV-2 infection, however we found that upon expression in HEK293T cells Orf10 interacts with nine host proteins. Among these are multiple members of the Cullin RING E3 ligase 2 (CRL2) complex, including CUL2, ELOB, ELOC, RBX1 and ZYG11B. Cullin RING E3 ligases play a central role in viral infections, since they are commonly hijacked by viral proteins to ubiquitinate and degrade viral restriction factors. CRL2 has been previously found to be targeted by poxviral ANK/BC via a C-terminal BC box domain resulting in potent suppression of inflammatory cytokines production, including interferon[429]. Similarly, Human Papilloma Virus protein HPV16 E7 binds to an active CRL2 complex, and the association correlates with the ability of HPV16 E7 to transform cells[430]. ZYG11B, a substrate adapter of CUL2, is the highest scoring hit in the Orf10 interactome indicating that Orf10 might bind to the assembled $CUL2^{ZYG11B}$ complex. Interestingly, ZYG11B targets substrates with exposed N-terminal glycines for degradation[431]. Orf10 contains an N-terminal glycine but does not have lysine residues, suggesting a few possible models: (1) Orf10 hijacks the $CUL2^{ZYG11B}$ complex for ubiquitination and degradation of restriction factors, or (2) Orf10 blocks $CUL2^{ZYG11B}$ and prevents the ubiquitination of its targets, or (3) Orf10 is targeted by $CUL2^{ZYG11B}$ for degradation through N-terminal ubiquitination.

**Supplementary Discussion References**

50. Ooi, Y. S., Stiles, K. M., Liu, C. Y., Taylor, G. M. & Kielian, M. Genome-wide RNAi screen identifies novel host proteins required for alphavirus entry. *PLoS Pathog.* **9**, e1003835 (2013).
51. Han, J. *et al.* Genome-wide CRISPR/Cas9 Screen Identifies Host Factors Essential for Influenza Virus Replication. *Cell Rep.* **23**, 596–607 (2018).
52. Li, B. *et al.* Genome-wide CRISPR screen identifies host dependency factors for influenza A virus infection. *Nat. Commun.* **11**, 164 (2020).
53. Wang, S., Tukachinsky, H., Romano, F. B. & Rapoport, T. A. Cooperation of the ER-shaping proteins atlastin, lunapark, and reticulons to generate a tubular membrane network. *Elife* **5**, (2016).
54. Yamamoto, Y., Yoshida, A., Miyazaki, N., Iwasaki, K. & Sakisaka, T. Arl6IP1 has the ability to shape the mammalian ER membrane in a reticulon-like fashion. *Biochem. J* **458**, 69–79 (2014).
55. Diaz, A., Wang, X. & Ahlquist, P. Membrane-shaping host reticulon proteins play crucial roles in viral RNA replication compartment formation and function. *Proc. Natl. Acad. Sci. U. S. A.* **107**, 16291–16296 (2010).
56. Chen, Y.-J., Williams, J. M., Arvan, P. & Tsai, B. Reticulon protects the integrity of the ER membrane during ER escape of large macromolecular protein complexes. *J. Cell Biol.* **219**, (2020).
57. Haenssler, E., Ramabhadran, V., Murphy, C. S., Heidtman, M. I. & Isberg, R. R. Endoplasmic Reticulum Tubule Protein Reticulon 4 Associates with the Legionella pneumophila Vacuole and with Translocated Substrate Ceg9. *Infect. Immun.* **83**, 3479–3489 (2015).
58. Hafirassou, M. L. *et al.* A Global Interactome Map of the Dengue Virus NS1 Identifies Virus Restriction and Dependency Host Factors. *Cell Rep.* **21**, 3900–3913 (2017).
59. Singh, C. R. *et al.* Mechanisms of translational regulation by a human eIF5-mimic protein. *Nucleic Acids Res.* **39**, 8314–8328 (2011).
60. Cheng, D.-D. *et al.* Downregulation of BZW2 inhibits osteosarcoma cell growth by inactivating the Akt/mTOR signaling pathway. *Oncol. Rep.* **38**, 2116–2122 (2017).
61. Jin, X., Liao, M., Zhang, L., Yang, M. & Zhao, J. Role of the novel gene BZW2 in the development of hepatocellular carcinoma. *J. Cell. Physiol.* (2019) doi:10.1002/jcp.28331.
62. Sato, K. *et al.* Novel oncogene 5MP1 reprograms c-Myc translation initiation to drive malignant

phenotypes in colorectal cancer. *EBioMedicine* **44**, 387–402 (2019).

106. Gregersen, L. H. *et al.* MOV10 Is a 5' to 3' RNA helicase contributing to UPF1 mRNA target degradation

by translocation along 3' UTRs. *Mol. Cell* **54**, 573–585 (2014).

117. Graham, R. L., Sims, A. C., Brockway, S. M., Baric, R. S. & Denison, M. R. The nsp2 replicase proteins of murine hepatitis virus and severe acute respiratory syndrome coronavirus are dispensable for viral

replication. *J. Virol.* **79**, 13399–13411 (2005).

153. Garrus, J. E. *et al.* Tsg101 and the vacuolar protein sorting pathway are essential for HIV-1 budding. *Cell*

**107**, 55–65 (2001).

176. Brockway, S. M., Clay, C. T., Lu, X. T. & Denison, M. R. Characterization of the expression, intracellular localization, and replication complex association of the putative mouse hepatitis virus RNA-dependent

RNA polymerase. *J. Virol.* **77**, 10515–10527 (2003).

188. Nchoutmboube, J. A. *et al.* Increased long chain acyl-Coa synthetase activity and fatty acid import is linked to membrane synthesis for development of picornavirus replication organelles. *PLoS Pathog.* **9**,

e1003401 (2013).

237. Wang, B., Zhang, X. & Zhao, Z. Validation-based insertional mutagenesis for identification of Nup214 as a

host factor for EV71 replication in RD cells. *Biochem. Biophys. Res. Commun.* **437**, 452–456 (2013).

249. Porter, F. W., Brown, B. & Palmenberg, A. C. Nucleoporin phosphorylation triggered by the encephalomyocarditis virus leader protein is mediated by mitogen-activated protein kinases. *J. Virol.* **84**,

12538–12548 (2010).

261. Feng, P., Everly, D. N., Jr & Read, G. S. mRNA decay during herpes simplex virus (HSV) infections:

protein-protein interactions involving the HSV virion host shutoff protein and translation factors eIF4H and eIF4A. *J. Virol.* **79**, 9651–9664 (2005).

273. Joseph, J. S. *et al.* Crystal structure of nonstructural protein 10 from the severe acute respiratory

syndrome coronavirus reveals a novel fold with two zinc-binding motifs. *J. Virol.* **80**, 7894–7901 (2006).

298. Subramanian, A. *et al.* Auto-regulation of Secretory Flux by Sensing and Responding to the Folded Cargo

Protein Load in the Endoplasmic Reticulum. *Cell* **176**, 1461–1476.e23 (2019).

323. Yue, Y. *et al.* SARS-Coronavirus Open Reading Frame-3a drives multimodal necrotic cell death. *Cell*

*Death Dis.* **9**, 904 (2018).

336. Chen, L.-H., Liao, C.-Y., Lai, L.-C., Tsai, M.-H. & Chuang, E. Y. Semaphorin 6A Attenuates the Migration

Capability of Lung Cancer Cells via the NRF2/HMOX1 Axis. *Sci. Rep.* **9**, 13302 (2019).

348. Kopecky-Bromberg, S. A., Martínez-Sobrido, L., Frieman, M., Baric, R. A. & Palese, P. Severe acute

respiratory syndrome coronavirus open reading frame (ORF) 3b, ORF 6, and nucleocapsid proteins function as interferon antagonists. *J. Virol.* **81**, 548–557 (2007).

360. Fielding, B. C. *et al.* Characterization of a unique group-specific protein (U122) of the severe acute

respiratory syndrome coronavirus. *J. Virol.* **78**, 7311–7318 (2004).

381. Ye, J. *et al.* Molecular pathology in the lungs of severe acute respiratory syndrome patients. *Am. J.*

*Pathol.* **170**, 538–545 (2007).

393. Sharma, K. *et al.* SARS-CoV 9b protein diffuses into nucleus, undergoes active Crm1 mediated

nucleocytoplasmic export and triggers apoptosis when retained in the nucleus. *PLoS One* **6**, e19436 (2011).

406. Shukla, A. & Hilgenfeld, R. Acquisition of new protein domains by coronaviruses: analysis of overlapping

genes coding for proteins N and 9b in SARS coronavirus. *Virus Genes* **50**, 29–38 (2015).
