## Supplementary Information Methods for "A SARS-CoV-2-Human Protein-Protein Interaction Map Reveals Drug Targets and Potential Drug-Repurposing"

**Supplemental methods**

**Supplemental Figure 1 Method:** All MS runs were compared and clustered using standard artMS (https://github.com/biodavidjm/artMS) procedures on observed feature intensities computed by MaxQuant. Supplemental Figure 1 shows all Pearson's pairwise correlations between MS runs, and are clustered according to similar correlation patterns.

**Supplemental Figure 2 Method:** See main text.

**Supplemental Figure 3 Method: PFAM domain enrichment analysis.** The enrichment of individual PFAM domains (or PFAM clans)(El-Gebali et al. 2019) was calculated with a hypergeometric test where success is defined as number of domains, and the number of trials is the number of individual preys pulled-down with each viral bait. The population values were the numbers of individual PFAM domains and clans in the human proteome. To make sure that the p-values that signify enrichment were meaningful, we only considered PFAM domains that have been pulled-down at least three times with any SARS-CoV-2 protein, and which occur in the human proteome at least five times. In SI Figure 3 we show PFAM domains/clans with the lowest p-value for a given viral bait protein.

**Supplemental Figure 4 and 5 Method: Expression analysis of interacting genes.** We used GTEx (version 8, median gene-level transcripts per million (TPM) by tissue), which consisted of 17382 samples (578 lung samples)(Melé et al. 2015) to examine the mRNA expression of all interacting proteins (n=323). The comparison gene group was all RefSeq genes (n=24,491). The lung expression values represent the median expression of each gene across the GTEx lung samples. The lung enrichment values are calculated by dividing the median expression of each gene in lung tissue by the median expression of each gene across all tissues (including lung). A value of greater than one indicates that the gene expression is enriched in lung tissue. Values were plotted on a log10 scale. All figures and statistics were produced in Python3 and code and reference tables can be found at: (https://github.com/stephaniewanko/Fraser_Lab/tree/master/QCRG_COVID19).

**Supplemental Figure 6 Method: Conservation analysis of interacting genes.** We used gnomAD version 2.1(Karczewski et al. 2019), which consists of 125,748 exomes and 15,708 genomes, to determine human genetic variation observed in the interacting proteins (n=323) versus all Refseq genes (n=24,491). Briefly the observed/expected ratio per gene indicates the number of observed variants of that type divided by the number of expected mutations of that type, with a lower observed/expected ratio indicating strong intolerance toward mutation. The number of expected variants were estimated based on the number of CpG and non-CpG transitions observed across the genome(Karczewski et al. 2019). All figures and statistics were produced in Python3 and code and reference tables can be found at:
(https://github.com/stephaniewanko/Fraser_Lab/tree/master/QCRG_COVID19).

**Supplemental Figure 7 Method: Nsp5 main protease (3Clpro) cleavage prediction.** We used sequence specificity data for SARS nsp5(Goetz et al. 2007) (98.7% identical to SARS-CoV-2 nsp5) and NetCorona(Kiemer et al. 2004) to predict cleavage sites within interacting factors. PDB ID: 1UJ1 served as template for peptide docking which was performed using the predicted P4-P1 residues (BioLuminate, Schrödinger, LLC). Illustration of the docked model was generated in PyMol (Schrödinger, LLC).

**Supplemental Figure 8 Method: Orf6 consensus sequence analysis.** Orf6 sequence homologs were identified using the BLAST tool(Johnson et al. 2008) (accession number YP_009724394.1), run with the default settings: gap opening and extension costs of 11 and 1, respectively, BLOSUM62 as the scoring matrix, and an e-value threshold of 10. The search yielded 34 homologous sequences. The multiple sequence alignment was visualized using the MView web server: https://www.ebi.ac.uk/Tools/msa/mview/ (Brown, Leroy, and Sander 1998) and the WebLogo server (https://weblogo.berkeley.edu/logo.cgi)(Crooks et al. 2004).

**Supplemental Table 1 Method:** See main text.

**Supplemental Table 2 Method:** See main text.

### Supplementary Methods References


Brown, N. P., C. Leroy, and C. Sander. 1998. "MView: A Web-Compatible Database Search or Multiple Alignment Viewer." *Bioinformatics* 14 (4): 380–81. https://doi.org/10.1093/bioinformatics/14.4.380.

Crooks, Gavin E., Gary Hon, John-Marc Chandonia, and Steven E. Brenner. 2004. "WebLogo: A Sequence Logo Generator." *Genome Research* 14 (6): 1188–90. https://doi.org/10.1101/gr.849004.

El-Gebali, Sara, Jaina Mistry, Alex Bateman, Sean R. Eddy, Aurélien Luciani, Simon C. Potter, Matloob Qureshi, et al. 2019. "The Pfam Protein Families Database in 2019." *Nucleic Acids Research* 47 (D1): D427–32. https://doi.org/10.1093/nar/gky995.

Goetz, D. H., Y. Choe, E. Hansell, Y. T. Chen, M. McDowell, C. B. Jonsson, W. R. Roush, J. McKerrow, and C. S. Craik. 2007. "Substrate Specificity Profiling and Identification of a New Class of Inhibitor for the Major Protease of the SARS Coronavirus." *Biochemistry* 46 (30): 8744–52. https://doi.org/10.1021/bi0621415.

Johnson, Mark, Irena Zaretskaya, Yan Raytselis, Yuri Merezhuk, Scott McGinnis, and Thomas L. Madden. 2008. "NCBI BLAST: A Better Web Interface." *Nucleic Acids Research* 36 (Web Server issue): W5–9. https://doi.org/10.1093/nar/gkn201.

Karczewski, Konrad J., Laurent C. Francioli, Grace Tiao, Beryl B. Cummings, Jessica Alföldi, Qingbo Wang, Ryan L. Collins, et al. 2019. "Variation across 141,456 Human Exomes and Genomes Reveals the Spectrum of Loss-of-Function Intolerance across Human Protein-Coding Genes." *bioRxiv*. https://doi.org/10.1101/531210.

Kiemer, Lars, Ole Lund, Søren Brunak, and Nikolaj Blom. 2004. "Coronavirus 3CLpro Proteinase Cleavage Sites: Possible Relevance to SARS Virus Pathology." *BMC Bioinformatics* 5 (June): 72. https://doi.org/10.1186/1471-2105-5-72.

Melé, Marta, Pedro G. Ferreira, Ferran Reverter, David S. DeLuca, Jean Monlong, Michael Sammeth, Taylor R. Young, et al. 2015. "Human Genomics. The Human Transcriptome across Tissues and Individuals." *Science* 348 (6235): 660–65. https://doi.org/10.1126/science.aaa0355.
